# Chimeric Origins and Dynamic Evolution of Central Carbon Metabolism in Eukaryotes

**DOI:** 10.1101/2024.05.29.596406

**Authors:** Carlos Santana-Molina, Tom A. Williams, Berend Snel, Anja Spang

**Author notes:** equal contribution, co-correspondence, alphabetic order (Berend Snel, Anja Spang).

## Abstract

The origin of eukaryotes was a key event in the history of life. Current leading hypotheses propose that a symbiosis between an asgardarchaeal host cell and an alphaproteobacterial endosymbiont represented a crucial step in eukaryotic origins and invoke a central role for syntrophic interactions - that is, metabolic cross-feeding between the partners as the basis for their subsequent evolutionary integration. A major unanswered question is whether the metabolism of modern eukaryotes bears any vestige of this ancestral interaction. To investigate this question in detail, we systematically analyze the evolutionary origins of the eukaryotic gene repertoires mediating central carbon metabolism. Our phylogenetic and sequence analyses reveal that this gene repertoire is chimeric, with ancestral contributions from Asgardarchaeota and Alphaproteobacteria operating predominantly in glycolysis and TCA, respectively. Furthermore, our analyses reveal diverse additional contributions from other prokaryotic sources as well as the extent to which this ancestral metabolic interplay has been remodeled via gene loss, transfer, and subcellular retargeting in the >2Ga since the origin of eukaryotic cells. Together, our work demonstrates that in contrast to previous assumptions, the eukaryotic metabolism preserves information about the nature of the original asgardarchaeal-alphaproteobacterial interactions, and supports syntrophy scenarios on the origin of the eukaryotic cell.

## Introduction

The origin of eukaryotes represents a defining event in life’s history that occurred between 2.6-1.2 Ga years ago^1–5^, possibly coinciding with rising oxygen levels in Earth’s atmosphere^6–8^. One of the predicted key steps during eukaryogenesis involved the symbiotic integration between a member of the Asgardarchaeota^9–12^ and a bacterial partner related to the Alphaproteobacteria that evolved to become the mitochondrion^13,14^.

To explain the evolutionary driving force underlying eukaryogenesis, a multitude of models have been proposed^15–29^ that differ with regard to predictions on the identity and number of partners involved as well as the nature of interactions ranging from syntrophy to phagocytosis and parasitism. Since the discovery of the Asgardarchaeota, symbiogenetic models invoking a syntrophic relationship between at least one archaeal and bacterial partner have been increasingly articulated^26,28^. Specifically, metabolic capabilities inferred for the asgardarchaeal ancestor of eukaryotes, have inspired updated hypotheses on potential syntrophic interactions between the archaeal and bacterial partners that may have played a role in the early stages of eukaryogenesis^24,25,30^. These syntrophy-based hypotheses suggest that one partner may have been dependent on the other as an external electron sink, but make distinct predictions about the types of metabolites being exchanged, the transition of lipid membranes, the timing of mitochondrial acquisition, the mechanism of mitochondrial uptake, and the origin of the nucleus^26,28,31–33^. However, testing these scenarios with current data is challenging, in part because the evolutionary origin of eukaryotic metabolism remains understudied.

Previous genomic analyses have suggested that ‘informational’ genes (those involved in translation, replication and transcription) have archaeal origins, while ‘operational’ genes (those involved in metabolism) derive from bacteria, particularly from the pre-mitochondrial endosymbiont^34–39^. This has led to the hypothesis that during eukaryogenesis, the archaeal host metabolism was replaced by counterparts from the endosymbiont^15^. However, considering that syntrophy relies on metabolic repertoires from both partners, archaeal gene contributions to eukaryotic metabolisms might be expected.

To assess current models on the microbial origins of eukaryotic cells as well as the evolution of plastids, we here analyze the origins of the eukaryotic central carbon metabolism (CCM), comprising four main pathways: the Embden–Meyerhof–Parnas (EMP) and the Entner-Doudoroff glycolytic pathways, the pentose phosphate pathway including pyruvate/acetate conversions and the tricarboxylic acid (TCA) cycle (**Supplementary Text**). While it has previously been suggested that many TCA cycle enzymes, as well as enzymes involved in pyruvate conversions, were present in LECA and trace their origin back to Alphaproteobacteria^13,40–42^, the evolutionary origins of eukaryotic enzymes of glycolysis and PPP remain unresolved^20,40^. Furthermore, a number of metabolic genes in eukaryotes appear to have origins unrelated to either symbiotic partner, potentially reflecting independent horizontal acquisitions either before or after the radiation of the extant eukaryotic lineages^43–45^, further complicating phylogenetic analyses। The metagenomics-based discovery of various new archaeal and bacterial lineages during the past decades, including the Asgardarchaeota, has provided a wealth of new information to address the origins and evolution of CCM enzymes of Eukaryotes within the context of a more broadly sampled tree of life^9,46–49^. Our comprehensive phylogenetic analyses reveal a much more complex pattern of evolution than previously anticipated and identify a chimeric CCM that notably includes contributions of archaeal origin to the LECA proteome. The distributions of CCM enzymes across the eukaryotic tree of life illuminate the subsequent highly dynamic evolution of these enzymes shaped by gene loss, endosymbiotic gene transfers, as well as paralogous and analogous gene replacements.

## Results

### Eukaryotic Tree of Life diversified into three major supergroups: Excavata, Amorphea and Diaphoretickes

To be able to compare gene trees of CCM enzymes with a eukaryotic tree of life, we selected a balanced and representative set of 207 eukaryotic proteomes. These cover currently known taxonomic diversity and lifestyles, including anaerobic eukaryotes and eukaryotes with primary and higher-order plastid organelles (**Fig. 1A** and **Table S1**). We reconstructed a species tree using various approaches, including heterogeneous-site trimming (*see methods*) to evaluate the robustness of the support for major groups of eukaryotes (**Fig. 1B, S1** and **Supplementary Text**). The resulting tree comprises three major supergroups (**Fig. 1A**): Excavata, Amorphea and Diaphoretickes. Although we rooted the tree between Excavata and the other groups for visualization, the placement of the root remains under debate^48,50–54^ and our interpretations of gene family origins do not assume a particular root position. Excavata include Discoba which appear to have retained ancestral traits such as a larger set of mitochondrial genes^55–57^, and include the anaerobic Metamonada, members of which usually form long branches in phylogenetic trees. The placement of Anaeramoebae, which has recently been shown to belong to Metamonada^58^, was unstable, alternatively branching with Amorphea clades (**Fig. S1B**). Amorphea includes Obozoa (Fungi, Metazoa, and various protists), Amoebozoa, and other putative relatives like CRuMs, Malawimonada and Ancyromonada. The monophyly of Amorphea is not stable: while Ancyromonadida most consistently places within Amorphea, we also observed its clustering with Diaphoretickes (**Fig. 1B** and **Supplementary Text**). Diaphoretickes form a stable group composed of two well supported monophyletic clades: the Cryptista-Archaeplastida (the later including Chloroplastida, Rhodophyta, Glaucophyta, Picozoa and Rhodelphis), and SAR (Stramenopile, Alveolata and Rhizaria). Apart from these, Haptista, Telonemia, Hemimastigophora and Aconracysta (the latter now classified as Provora phylum^59^) showed varying placements within Diaphoretickes, depending on site filtering and phylogenetic methods (**Fig. S1**). Overall, the inferred eukaryotic tree of life, and the inference of the three major supergroups, Excavata, Amorphea and Diaphoretickes, provide a solid framework for interpreting individual gene trees, defining LECA versus post-LECA clades and thereby determining the relative timing of gene acquisitions.

**Fig. 1.**
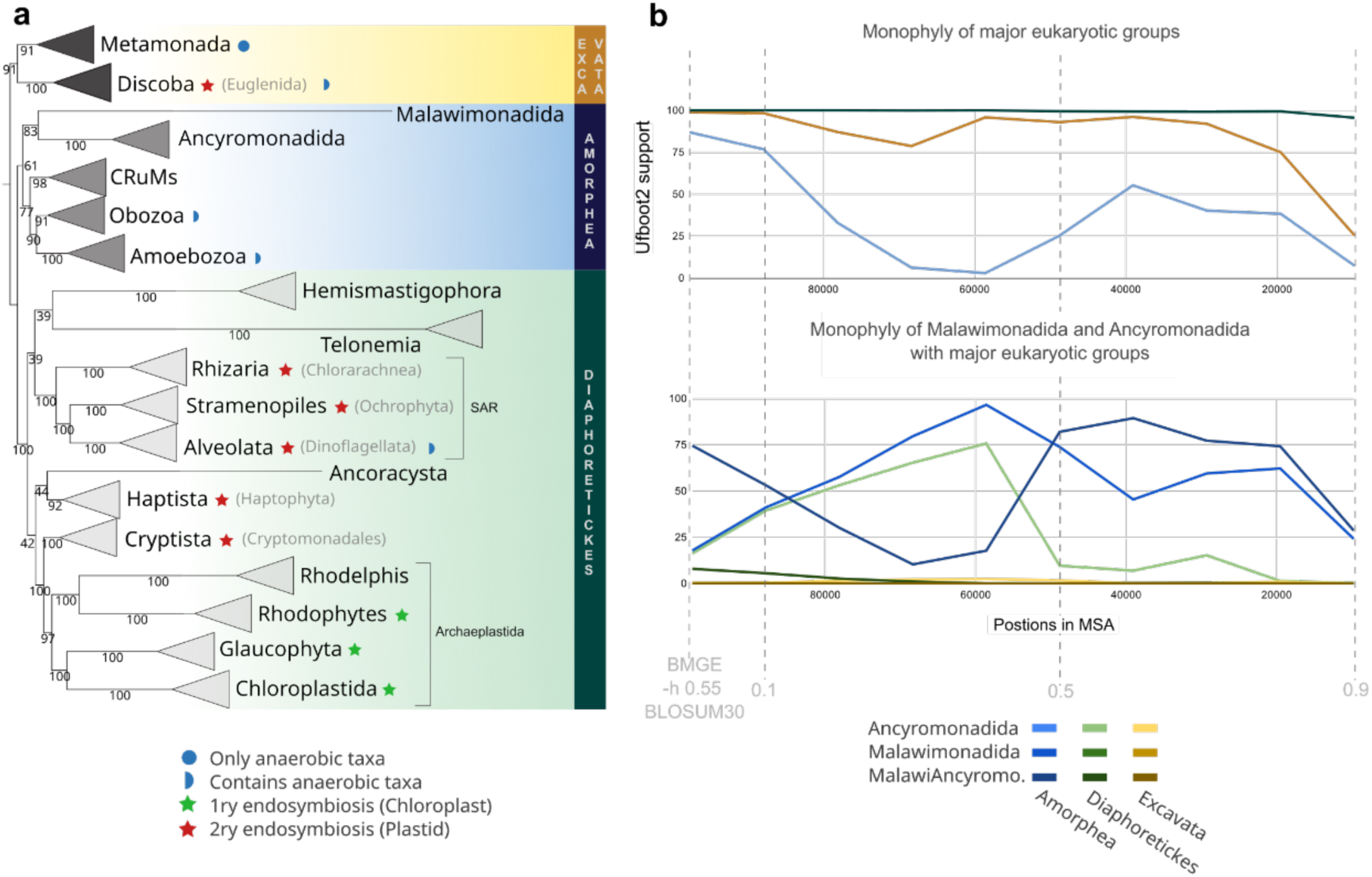
Phylogenetic reconstruction of the eukaryotic tree of life. **A)** Maximum-likelihood phylogeny of the eukaryotic tree of life based on the concatenation of 317 phylogenetic markers. The tree is unrooted, but drawn with Excavata at the root for ease of visualisation. The concatenated MSA consisted of 207 taxa, 97,680 positions, and the tree was built with IQTREE.2.1.2 under LG+C60+G model and using optimized ultrafast bootstrap (Ufboot2 -bnni). Annotation corresponds to characteristic traits (see legend). **B)** Chart depicting Ufboot2 of major groups (Excavata, Amorphea and Diaphoretickes, upper panel) and unstable groups (Ancyromonadida and Malawimonadida, lower panel) on intervals along removal of heterogeneous sites. Note that Amorphea in the upper panel only includes Obozoa, Amoebozoa and CRuMs. Extended tree and additional phylogenetic analyses in **Fig. S1** .

### Central Carbon Metabolism was established in the Last Eukaryotic Common Ancestor

We inferred the phylogenies of 64 gene families encoding enzymes involved in the CCM of eukaryotes and evaluated the evolutionary origins of eukaryotic clades in each phylogeny (**Fig. S2-41** and **Supplementary Text**). Specifically, we performed careful and iterative phylogenetic analyses that included an extensive taxonomic sampling (*see methods,* **Fig. S2**) to gain resolution in determining the taxonomic identity of sister groups.

Our phylogenetic analyses suggest that 9 out of 10 EMP (**Fig. S3-14**) and 7 out of 8 pentose-phosphate pathways (**Fig. S15-24,** and **Supplementary Text**) enzymes comprise putative LECA clades, respectively, suggesting that these pathways are ancestral in eukaryotes. By contrast, phylogenies of Entner-Doudoroff glycolytic enzymes seem to indicate that these enzymes have been acquired later during eukaryotic evolution in photosynthetic eukaryotes and representatives with secondary endosymbionts^60^ (**Fig. S23**). For pyruvate/acetate conversions, we observed that the pyruvate dehydrogenase complex (PDH formed by PDHA/B/C/D subunits) as well as acetyl-CoA synthetase (ACS) and lactate dehydrogenases (LDH), were present in LECA, while their respective counterparts, i.e. pyruvate formate-lyase (PFL), pyruvate-ferredoxin/flavodoxin oxidoreductase (POR), and ADP-forming acetyl-CoA synthetase (ACDA/B), are present in anaerobic Metamonada Archamoebae, and Breviatea among others including aerobic organisms (**Fig. S25-29** and **Supplementary Text**). Most of these likely reflect post-LECA acquisitions with subsequent transfer among eukaryotes, while others such as (POR) may have been present in LECA^45^. The phylogenies of the reverse TCA cycle defined by ATP-citrate lyase subunits (ACLA/B) or its fused version (ACLY; **Fig. S30**), suggested that both versions were likely present in LECA as suggested previously^61^. Finally, for the TCA cycle, all phylogenies for the 10 enzymatic steps showed clear LECA clades (**Fig. S31-40** and **Supplementary Text**), indicating that the TCA was present in LECA. Yet, the TCA was nearly absent in Metamonada, Archaemoeba, Microsporidia and partially in Breviatea in line with suggested secondary losses in these eukaryotic clades^62^. As previously observed^58,62–64^, in the case of two mitochondrial genes found in metamonads, PDHD and FUMC, their sequences form a monophyletic clade with those of other eukaryotes (**Fig. S25,37B**), consistent with the hypothesis that these organisms secondarily lost mitochondria, rather than that they never had them^54^.

Notably, the phylogeny of some enzymatic steps associated with these metabolic pathways revealed the presence of independent orthogroups coding for enzymes predicted to perform the same reactions. Examples include phosphoglycerate mutases (in EMP), aconitases (ACO) and isocitrate dehydrogenases (IDH, in TCA), suggesting that LECA harbored metabolic redundancy for these enzymatic steps. Altogether, these phylogenies unambiguously show that LECA had the complete set of enzymes needed for a complete CCM.

### Diverse origins for the assembly of Central Carbon Metabolism in eukaryotes

We classified our phylogenies into distinct categories according to tree topology and the inferred donor lineage for each eukaryotic clade (see **Fig. 2** for representative examples and **Fig. 3, S40** for the distribution and potential origins of all CCM enzymes considered in this study).

**Fig. 2.**
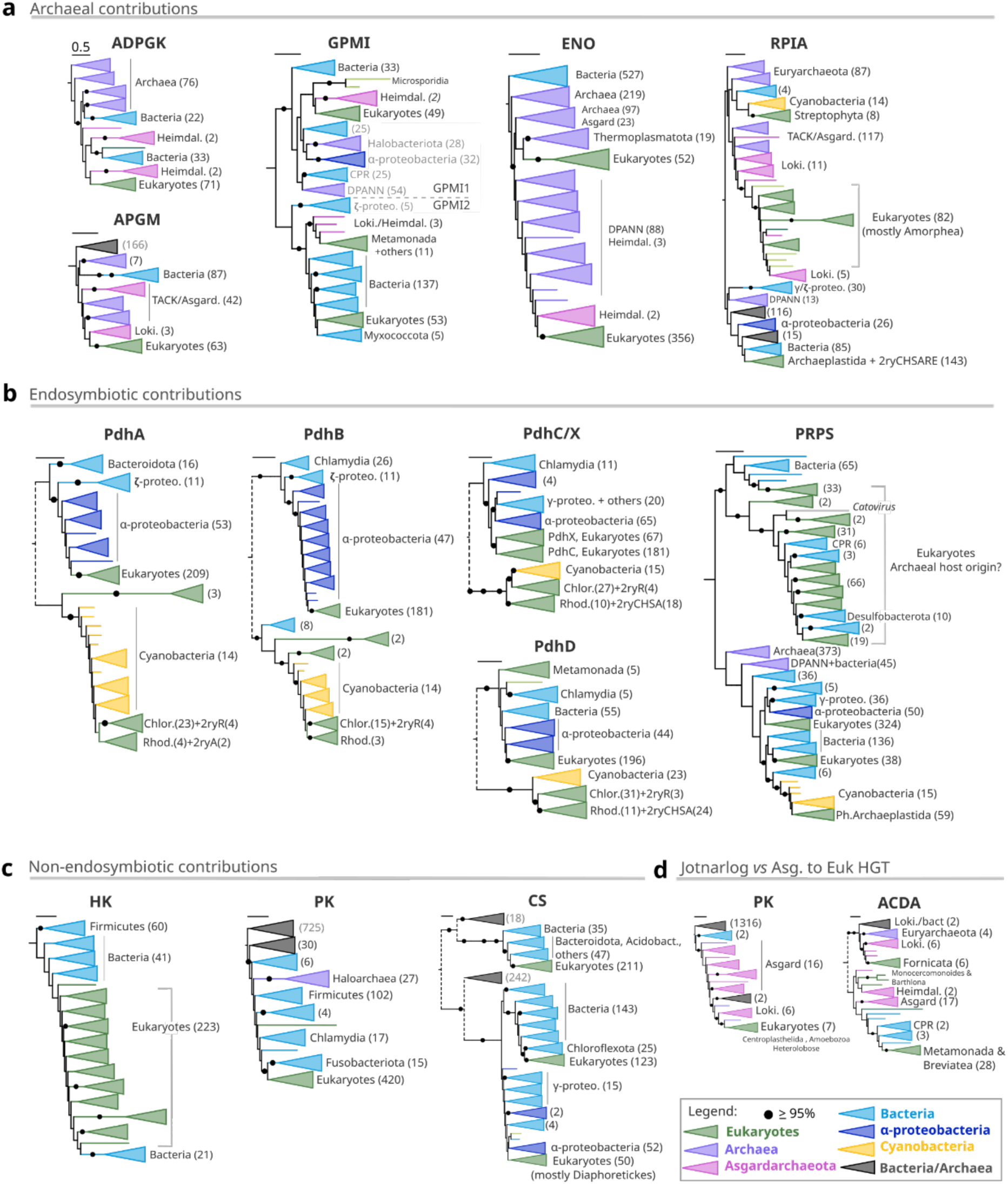
Maximum-likelihood phylogenies of selected enzymes representing diverse evolutionary origins of the eukaryotic central carbon metabolism (CCM) in eukaryotes. Examples of potential **A)** archaeal contributions from Asgardarchaeota, **B)** contributions from Alphaproteobacteria and Cyanobacteria, **C)** other prokaryotic origins (from known or unknown donor), and **D)** jotnarlogs *versus* Asgardarchaeota to eukaryote horizontal gene transfer. Discontinuous lines indicate simplified tree topology after pruning the relative branches of interest. Number between brackets denotes the number of sequences in the respective clades. 2ryCHSRA, denotes secondary endosymbionts including Cryptista (C), Haptista (H), Stramenopile (S), Rhizaria (R) and Alveolata (A). Phylogenies were built with IQTREE.2.1.2 under LG+C20+G+F model and using optimized ultrafast bootstrap (NNI UfBoot2). Extended trees provided in **Fig. S3-39**. Complete enzyme gene names: ADP-dependent glucokinase (ADPGK), 2,3-bisphosphoglycerate-independent phosphoglycerate mutase (APGM and GPMI), enolase (ENO), ribose 5-phosphate isomerase A (RPIA), pyruvate dehydrogenase complex subunits (PDHA/B/C/D), ribose-phosphate pyrophosphokinase (PRPS), hexokinase (HK), pyruvate kinase (PK), citrate synthase (CS) and ADP-forming acetate---CoA (ACDA).

**Fig. 3.**
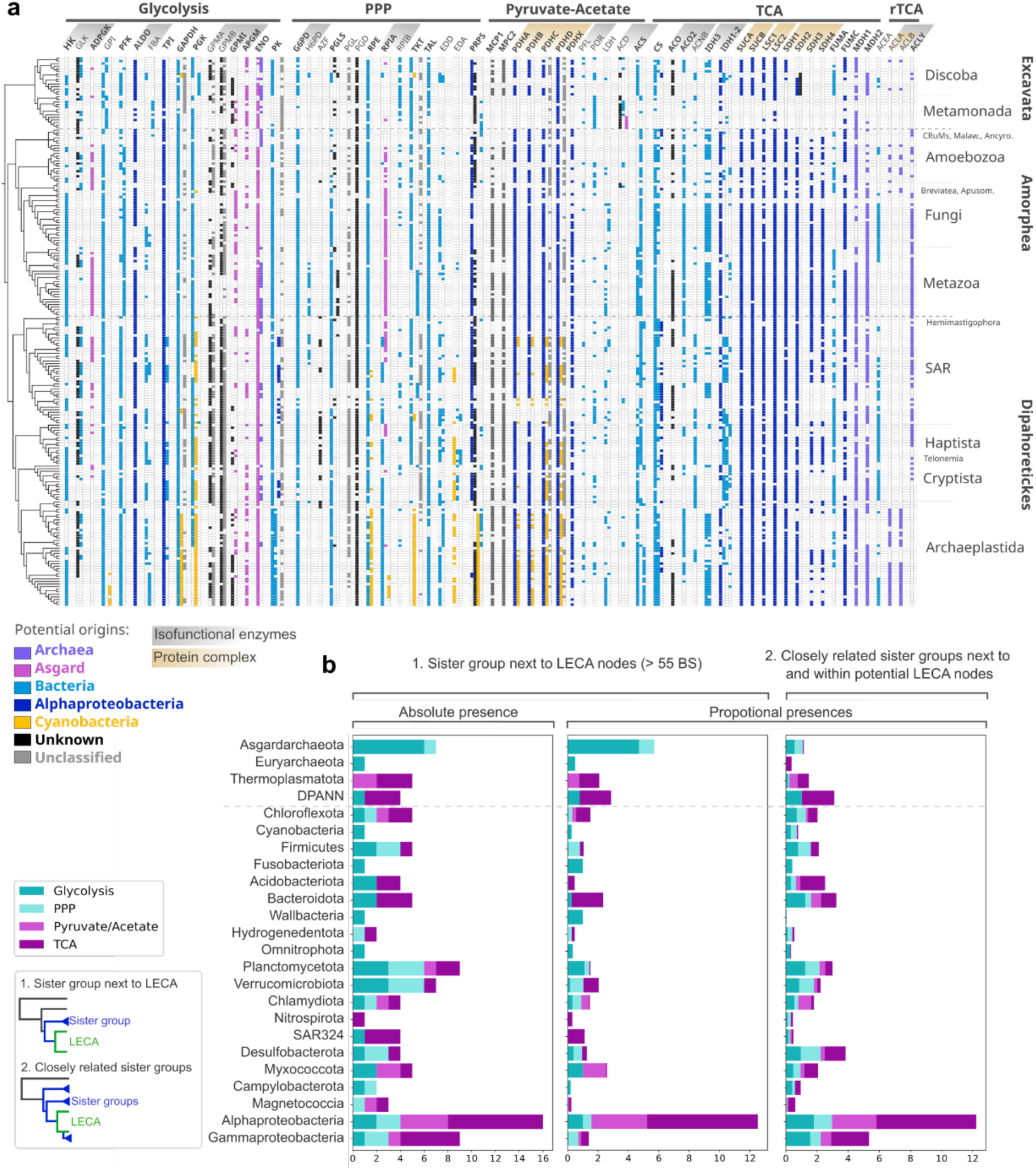
Prokaryotic origins and distribution of eukaryotic CCM. **A)** Phylogenetic profile of eukaryotic orthogroups manually selected from phylogenetic trees. Coloured cells indicate presence of genes with a proposed origin, see legend. Unknown origins (black cells), refer to those unresolved phylogenies. Gray cells column indicate sequences not considered as orthogroups (*unclassified*), and they are shown if they are present in more than 80 eukaryotes. Bold enzyme names are those hypothesized to be present in LECA, and gray and light-brown denote isofunctional and protein complex enzymes, respectively. Eukaryotic tree of life includes fast-evolving and low genome completeness taxa like Microsporidia and Picozoa (see extended Fig. in **Fig. S40**). Raw data for absence/presence profile provided in **Table S5**. **B)** General taxonomic composition of sister group(s) of selected potential LECA clades. First and second panels (1) depict the taxonomic composition of the first/direct sister group to a LECA clade, while third panel (2) depicts the composition of the extended LECA sister groups (see legend for visual explanation). Bar length is the sum of the respective presences of a taxon. ‘Absolute presence’ counts the presence of a specific taxon in the sister group, while ‘proportional presences’ represent the proportion of a taxon respective to the size of the sister group(s). Those taxa whose proportional presence in the single sister group was >= 1, were shown in the plot. Different colored stacked bars refer to different pathways, and prokaryotic phyla were sorted according to phylogenetic relationships.

*Asgardarchaeal host contributions*. In four EMP phylogenies, we observed clusters of sequences from diverse eukaryotic LECA clades potentially branching sister to Asgardarchaeota (**Fig. 2A, S2,12,13**): ADP-dependent glucokinase (ADPGK, acting in the first and third step of EMP^65^), two 2,3-bisphosphoglycerate-independent mutases (APGM and GPMI, analogous enzymes) and enolase (ENO). Among these, three phylogenies unequivocally support an archaeal origin (ADPGK, APGM and ENO), while the topology of GPMI_1 shows a sister relationship between a small number of asgardarchaeal and eukaryotic sequences. In the PPP, the ribose 5-phosphate isomerase A (RPIA) phylogeny shows a large eukaryotic cluster branching next to Asgardarchaeota (**Fig. 2A, S18**) although this cluster mainly contains species from the Amorphea and few Discoba (**Fig. 3A**). Thus, this indicates that Asgardarchaeota have contributed gene families to the CCM of eukaryotes.

Other cases of archaeal origins remain more speculative. Eukaryotic ATP-citrate lyase subunits (ACLA/B) or its fused version (ACLY), appear to be present in various Asgardarchaeota and might therefore have been part of the archaeal FECA proteome. However, the respective eukaryotic ACLA/B and ACLY clades branch with DPANN and Thermoplasmatota-E3 respectively, suggesting lack of signal, independent HGTs from different archaeal groups, or HGTs among archaea (**Fig S30** and **Supplementary Text**). Eukaryotic malate dehydrogenase LECA paralogs operate in the cytoplasm (MDH1) and in the mitochondria (MDH2)^66^. The phylogeny of the MDH family in combination with shared introns suggest that MDH1 and MDH2 originated by duplication in LECA, with a clade containing TACK and Baldarchaeota sequences as sister groups (**Fig. S38** and **Supplementary Text**). However the long branches characterizing these phylogenetic relationships render the archaeal origin of MDH1/2 tentative.

Lastly, in three phylogenies, we observed the clustering of a limited number of eukaryotes with Asgardarchaeota (**Fig 2D**). The pyruvate kinase (PK) phylogeny contains a phylogenetic group, PK_5, mainly composed of Amoebozoa (plus a few others), nested within an asgardarchaeal clade (**Fig. 2D and S14**). Furthermore, the phylogeny of GPMI showed, in addition to a clear LECA clade (GMPI_1), a small monophyletic group of Asgardarchaeota with some eukaryotes (labeled as GPMI_2, **Fig. 2A, S12** and **Supplementary Text**). The third case is the acetate-CoA ligase (ADP-forming) enzyme family (ACDA/B) which is involved in the reversible conversion of acetate into acetyl-CoA using ATP (counterpart to ACS). We found a fully supported monophyletic group (100% optimized ultrafast bootstrap, ACD3) containing Lokiachaeia and Fornicata, a subclade of Metamonda (**Fig. 2D, S29**). The clustering of these eukaryotic clades with Asgardarchaeota would be consistent with eukaryotic sequences being LECA clades that underwent massive loss later on, i.e jotnarlogs^67^ (eg. PK). Alternatively, this clustering might reflect a post-LECA transfer from Asgardarchaeota to the ancestor of one of these eukaryotic groups with subsequent transfer between eukaryotes. Overall, these cases show a previously unappreciated role of the asgardarchaeal host cell in shaping the eukaryotic proteome involved in the CCM.

#### Alpha- and cyanobacterial endosymbiotic contributions

The next category involves putative endosymbiotic contributions to LECA and the Archaeplastida ancestor, i.e. genes that were acquired through endosymbiotic gene transfer (EGT^68^). Indeed, many CCM phylogenies recovered Alphaproteobacteria as sister clades to eukaryotes (**Fig. 3A**). We found two alphaproteobacterial contributions to the EMP (fructose-bisphosphate aldolase, ALDO, and triosephosphate isomerase, TPI), one to the PPP (PRPS), 4 in pathways related to pyruvate conversions (PDHA/B/C/D) and 10 to the TCA cycle (IDH1, 2-oxoglutarate dehydrogenase subunits, SUCA/B, succinyl-CoA synthetase alpha/beta subunits, LSC1/2, succinate dehydrogenase subunits, SDH1/2/3/4, and fumarate dehydrogenase C, FUMC, **Fig. S7A,8,24,25,33-37**). Thus, most of the potential contributions from alphaproteobacteria seem to operate in the mitochondria, i.e. in 5 out of 10 steps of the TCA.

We also found several likely cyanobacterial contributions to the CCM of photosynthetic eukaryotes (**Fig. 3A**). Specifically, 4 enzymes of the EMP (glucose-6-phosphate isomerase, GPI, fructose-bisphosphate aldolase class II, FBA, glyceraldehyde 3-phosphate dehydrogenase, GAPDH, and phosphoglycerate kinase, PGK), 6 of the PPP (ribulose-phosphate 3-epimerase, RPE, RPIA, PRPS, transketolase, TKT, phosphogluconate dehydratase, EDD and 2-dehydro-3-deoxyphosphogluconate aldolase, EDA) and 4 among pyruvate conversions (PDHA/B/C/D), have topologies consistent with being derived by EGT from cyanobacteria (**Fig. S5,7B,9,10,19-25**). These phylogenetic clusters also contain photosynthetic eukaryotes with higher level plastids (e.g. secondary and tertiary endosymbioses), with PDH phylogenies clearly supporting distinct secondary endosymbiosis events involving green and red algae, respectively (**Fig. 2B**). The subsequent evolution in photosynthetic eukaryotes seems to have been highly dynamic involving losses, duplications and retargeting signals (**Fig. 3B, 4**; *see below*). It is important to acknowledge that the phylogenetic signal for the sister relationship between Alphaproteobacteria (or Cyanobacteria) and eukaryotes is not always unequivocal^13^ (*see* **Supplementary Text)**: in phylogenies of TPI and PRPS, eukaryotes are sister to Alpha-/Gammaproteobacteria, while trees of LSC2, and ACS, recover genes of other prokaryotic clades interspersed between the alphaproteobacterial/LECA clades. Furthermore, the PDHD and FUMC phylogenies include divergent eukaryotic sequences that branch within the Alphaproteobacteria clade rather than within the LECA clade (especially Excavata taxa; **Fig. S25,36**). Similarly, cyanobacterial contributions are not always highly supported or form well defined topologies (eg. RPE, EDD and EDA; **Fig. S19,23**). Nevertheless, our analysis shows that alphaproteobacterial contributions (LECA) mainly operate in the TCA pathway, and that cyanobacterial contributions (in Archaeplastida ancestor) often comprise enzymes of the EMP glycolysis and pentose-phosphate pathways. The PDH complex and PRPS phylogenies revealed contributions derived from both Alphaproteobacteria and Cyanobacteria (**Fig. 2B**), potentially illustrating the importance of pyruvate and ribose phosphate metabolisms in these endosymbioses.

#### Contributions to LECA CCM from other prokaryotic lineages

Besides contributions of the asgardarchaeal host and the alphaproteobacterial endosymbiont to the LECA proteome, several phylogenetic trees indicate donations from other prokaryotic lineages, some with good support (i.e optimized UfBoot2 support >95%). In some cases, these donations included enzyme families that lacked respective homologs in Asgardarchaeota and Alphaproteobacteria. Examples comprise phylogenies of glycolytic enzymes (such as hexokinase, HK, 6-phosphofructokinase 1, PFKA, GAPDH, PGK, and PK), enzymes involved in the PPP (glucose-6-phosphate 1-dehydrogenase, G6PD, 6-phosphogluconolactonase, PGLS, RPE, TKT and TAL), as well as TCA cycle enzymes (citrate synthase, CS, IDH1, FUMA/B). Potential donors that we identified included Chlamydia (TAL), Planctomycetota-Verrucomicrobiota (PGK, G6PD and RPE), Fusobacteriota (PK), Cyanobacteria, Dependinate (for two independent donations of LDH), and Chloroflexota (CS) (**Fig. S14,15,19,31,38**). However, a number of phylogenies displayed a mixed composition of sister groups which hindered the identification of the donor for the respective clade (eg. GAPDH **Fig. 9**). In spite of this difficulty, we observed the presence of recurrent phyla in the sister groups, including Myxococota-Desulfobacterota, Bacteroidota and Acidobacteriota among others (**Fig. 3B**). These examples suggest that prokaryotes other than Asgardarchaeota and Alphaproteobacteria have contributed to the assembly of CCM during and/or after eukaryogenesis.

#### Gene families of unresolved origins

In several (at least 12) phylogenetic reconstructions, it was not possible to clearly denote LECA clades due to paraphyletic branching of eukaryotic and prokaryotic sequences resulting in unresolved sister groups. While the phylogenetic signal was limited in some cases (GPMA, GMPB or PGL), others recovered consistent and robust topologies across a range of datasets and analyses i.e. glucokinase, GLK, GPI, EDA/D, PRPS1, and ACS, **Fig. S3,5,18,24,28**). For example, our phylogeny of GPI is consistent with previous work that also resolved paraphyletic clades for eukaryotic homologs^69,70^ (**Fig. S5**). Our investigation of the conservation of introns in eukaryotic GPI in light of phylogenetic signals across its MSA, suggested that GPI was present in LECA (**Fig. S5C** and **Supplementary Text**). In addition, the GPI full length MSA phylogeny showed a paraphyletic group of Cryptophycea and Chlamydiota branching between a plastid and the aforementioned tentative LECA clade. However, this intermediary position changed when considering different part of the alignment: a region that surrounds a shared intron between Cryptophyceae and plastids provided a monophyletic topology of these species sharing the intron, while the rest of the C-terminus provided a monophyletic branching of Cryptophycea/Chlamydiota and the ‘LECA’ clade. (**Fig. S5D**). This indicates that some CCM enzymes of eukaryotes with distinct prokaryotic origins underwent recombination events (**Supplementary Text**). The PRPS phylogeny recovers an unresolved eukaryotic group (PRPS1), that might be derived from archaeal PRPS. However, this is speculative due to the presence of interspersed bacterial groups (**Fig. 2B, S24** and **Supplementary Text**). Thus, the evolutionary origins of some of these eukaryotic genes remain unresolved, likely due to a combination of events including gene exchange during and after eukaryogenesis, different evolutionary rates in prokaryotes and eukaryotes, loss of signal, and recombination.

### Functional and dynamic evolution of CCM enzymes during eukaryotic diversification

We next investigated post-LECA evolution of CCM enzymes including their distribution across the eukaryotic tree and their predicted organellar localisation as inferred from organelle targeting sequences. The analysis of CCM enzyme distribution across the tree revealed that orthogroup repertoires vary between distinct eukaryotic clades (**Fig. 3A**). For example, we identified both cases of replacement and differential retainment of isofunctional enzymes (eg. HK/Glucokinase, GLK/ADPGK, ALDO/FBA, PGMA/PGMB/GPMI/APGM, RPIA/RPIB, PDH/POR, ACS/ACDAB, ACO/ACO2, IDH1/IDH2/IDH3, FUMAB/FUMC), as well as metabolic re-adaptations (eg. loss of the PDH complex and TCA in Metamonada). The evolutionary history of enzymes of inferred Asgardarchaeota origin, like ADPGK, APGM and RPIA, suggests that these genes were present in LECA but subsequently replaced by horizontally acquired paralogous or analogous enzymes of bacterial origin (HK, PGMA/B, RPIB respectively) in some eukaryotic lineages. Others, like PDHA/B/C/D, IDH3 (alpha and gamma subunits), and ACLA/B presented high correlative distributions in line with their forming protein complexes (**Table S5**). A correlation network (+/-0.5 phi-coefficient cut-off) between orthogroups and lifestyle characteristics (e.g. anaerobic, primary and secondary endosymbiosis) suggests that at least some of the dynamics of CCM enzymes reflect eukaryotic lifestyles (**Fig. 4AB, S41** and **Supplementary Text**). Correlated distributions between photosynthetic eukaryotes (by/of primary and secondary endosymbioses) are mainly related to EMP/EDP and PPP, while correlations regarding aerobic/anaerobic lifestyle usually involved pyruvate/acetate conversions and TCA cycle enzymes. Phylogenies of POR, ACDA/B, GPI, FBA, and RPIB, displayed orthogroups involving anaerobic eukaryotes, suggesting adaptations to anoxygenic conditions (**Fig. 4AB**).

**Fig. 4.**
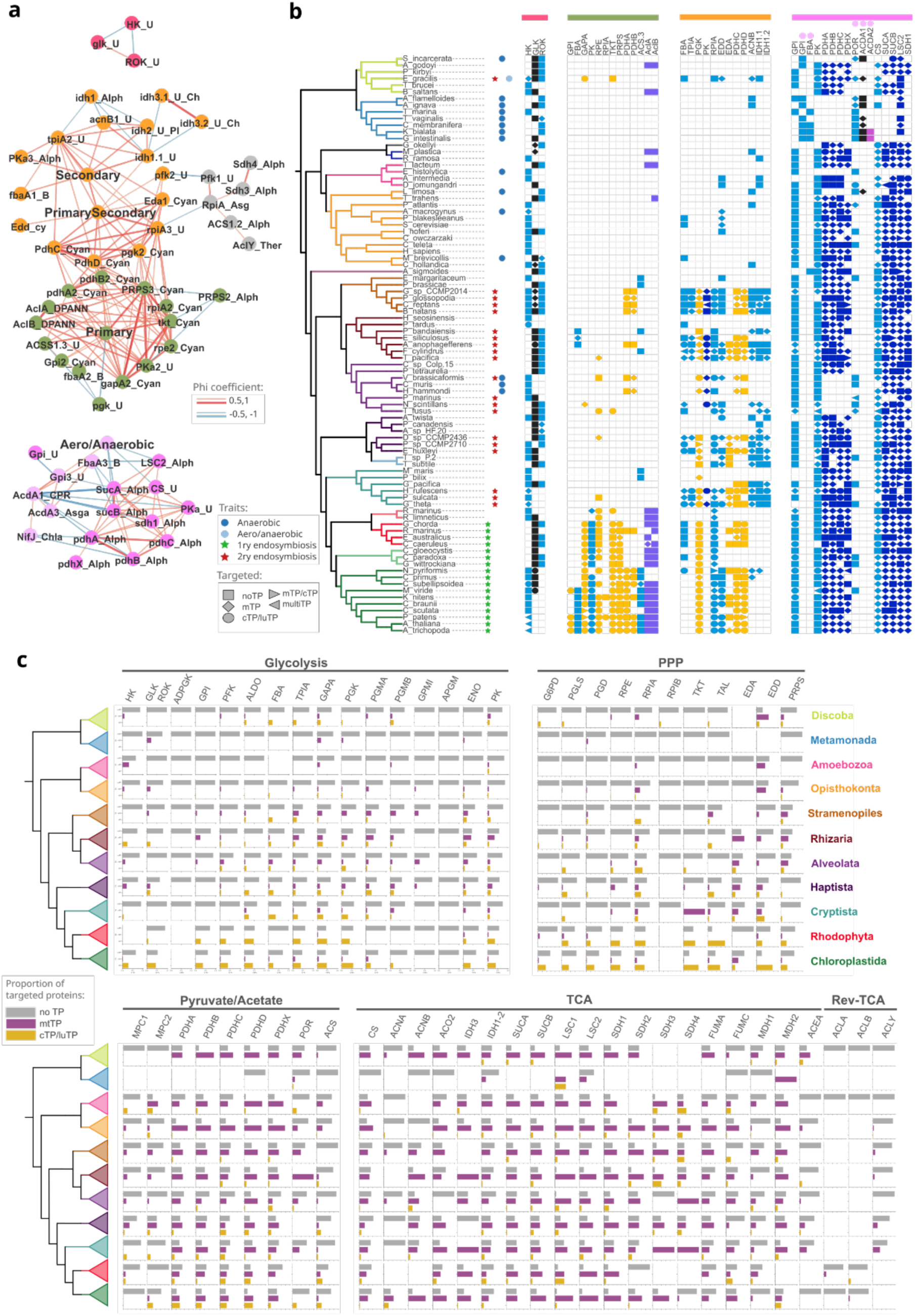
Correlative distributions of CCM enzymes across eukaryotes and their targeting in eukaryotic cells. **A)** Correlative networks for the distribution of those orthogroups with higher (red edges) or lower (blue edges) phi-coefficients than 0.5 and -0.5 respectively. Light pink indicates orthogroups including anaerobic eukaryotes. Clusters were obtained by modularity using Gephi. **B)** Phylogenetic profile of the respective correlated orthogroups indicating their evolutionary origins (cell color) and targeting signal (cell shape). Taxonomic tree is a subselection of representatives and is annotated with characteristic traits of the respective taxa (*see legend*). **C)** Distribution of targeted proteins along the eukaryotic tree of life and the central carbon metabolism. Bars represent the proportion of sequences with the respective targeting (see legend). Only sequences from selected orthogroups (Fig. 3A) were used for this analysis. Raw data for these plots provided in **Table S5**.

The majority of enzymes of the EMP, and PPP as well as the key enzymes of the reverse-TCA enzymes, do not encode a targeting signal, whereas most of the enzymes involved in pyruvate conversions and the TCA appear to be targeted to the mitochondria based on TargetP annotations (**Fig. 4C**). Nevertheless, exceptions exist, indicating potential sub- or neofunctionalizations of certain enzymes. For instance, PGMA and PGMB are typically found in both the cytoplasm and mitochondria/chloroplast, whereas their analogous enzymes, GPMI and APGM, do not exhibit mitochondrial targeting sequences (**Fig. 4C**). Likewise, in agreement with their general targeting patterns, MDH1 is generally associated with mitochondrial functions, whereas MDH2 tends to be associated with cytoplasmic activities^66^, although the reverse is true in some taxa (**Fig. 4C, S40**). On the other hand, not all proteins of alphaproteobacterial origin are targeted to the mitochondria (eg. ALDO, TPI), and conversely some enzymes of non-alphaproteobacterial origin appear to have mitochondrial targeting signals (eg. CS, ACNB, IDH1-2). Therefore, the distribution of targeted proteins across the CCM enzymes illustrate the general compartmentalization of these pathways in eukaryotic cells. However, the targeting of these is not always in agreement with their origins, suggesting an ongoing process of retargeting during the evolution of eukaryotes^71,72^.

Two cases exemplify the complexity of post-LECA retargeting of CCM: the chloroplast and mitochondrial glycolysis respectively (**Fig. 4C, S42**). In Archaeplastida, the genes coding for EMP (and PPP) enzymes are targeted to the cytoplasm and chloroplast, respectively^73^, and have originated by nuclear gene duplications, EGT, HGTs, retargeting and differential loss (summarized in **Fig. S43** and **Supplementary Text**). Specifically, our results highlight the frequent duplication and subsequent relocation of ‘nuclear’ genes to the photosynthetic organelle. Similarly, the parallel glycolysis in cytoplasm and mitochondria described in SAR^74,75^, appears to be specific to secondary endosymbionts involving the lower enzymatic steps, between TPI and PK (**Fig. S42,44** and **Supplementary Text**). Therefore, CCM enzymes are subject to organelle retargeting promoting their functional diversification in eukaryotic cells.

## Discussion

Our comprehensive phylogenetic analyses demonstrate that a complete set of eukaryotic CCM enzymes was likely present in LECA. These enzymes have complex evolutionary histories with eukaryotic genes originating from a variety of sources, including not only contributions from the alphaproteobacterial symbiont but also from the asgardarchaeal host and other prokaryotic donor lineages (**Fig. 5**). We found 6 putative contributions from Asgardarchaeota to the CCM of LECA, within the EMP and PPP: ADPGK, GPMI, APGM, ENO, PK, and RPIA, which is in contrast to previous work postulating that Asgardarchaeota did not contribute to eukaryotic CCM^12^, or that ENO was the only eukaryotic enzyme within carbon metabolism to be of archaeal origin^76,77^. Even more salient is the potential archaeal affiliation of MDH1/2 and ACLA/B/Y which are involved in the TCA and reverse-TCA cycles, and which might therefore represent archaeal host contributions that became integrated into the mitochondrial TCA cycle. With the exception of ENO, these Asgard host contributions are patchily distributed in extant eukaryotes, apparently due to independent horizontal replacement events from a variety of sources. This, combined with limited taxon sampling of prokaryotes and microbial eukaryotes in previous studies, might explain why these contributions previously went undetected. Our findings on asgardarchaeal contributions to the CCM strengthen the idea that eukaryotic metabolism is a product of integrating gene repertoires from symbiotic partners, rather than being derived solely from the mitochondrial progenitor^77^.

**Fig. 5.**
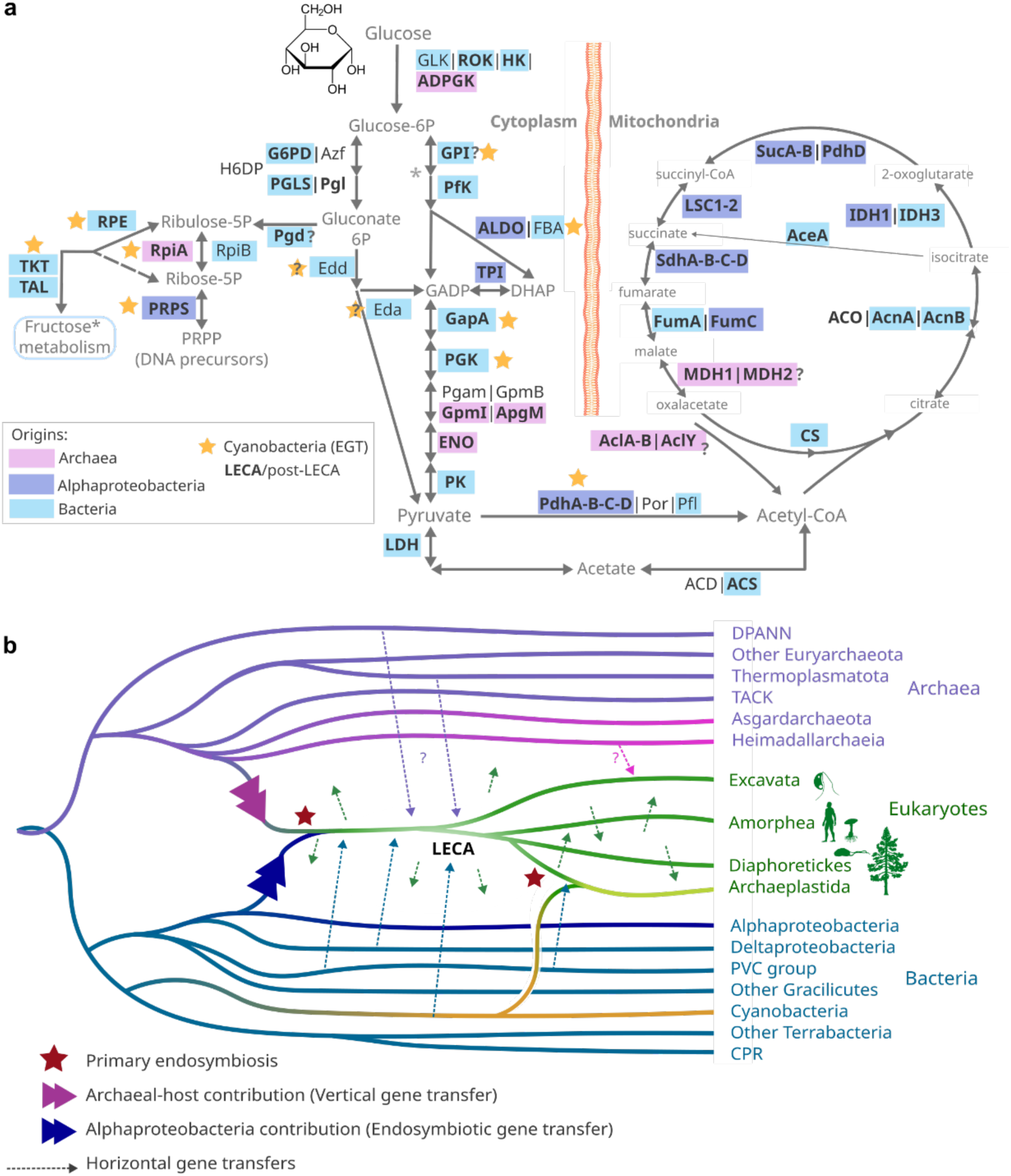
Evolutionary model for the origin of CCM during eukaryogenesis. **A)** Central carbon metabolic pathways highlighting the proposed origins by colors. Enzyme names in bold denote those enzymes potentially present in LECA. Interrogations indicate tentative inferences. **B)** Schematic representation of tree of life illustrating the evolutionary history for the assembly of CCM in eukaryotes. Please note that we only depict selected major routes of HGT, which is pervasive in both eukaryotic, archaeal and bacterial evolution. Eukaryotic icons were downloaded from PhyloPic.

We found 17 putative alphaproteobacterial contributions, most of which are predicted to operate in the mitochondria (except for TPI, ALDO and PRPS; **Fig. 5**). This finding is reminiscent of the evolutionary mosaicism previously reported for another essential process in eukaryotes, iron-sulphur cluster biosynthesis, in which the mitochondrial steps are predominantly alphaproteobacterial in origin, while the cytosolic steps are carried out by enzymes of varying evolutionary affinities^78^.

While eukaryotes appear to have several CCM genes acquired from different individual bacterial taxa other than alphaproteobacteria (**Fig. 3, S3-40**), no additional dominant source of gene donations is apparent (**Fig. 3B**). Thus, our analysis of the origin of eukaryotic CCM enzymes does not provide strong evidence for an additional endosymbiotic partner besides the alphaproteobacterial and cyanobacterial ancestors of mitochondria and plastids, respectively. It is noteworthy however, that a substantial number of phylogenies displayed a mixed composition of sister groups (**Fig. 3B**). This may be due to lack of phylogenetic signal, undersampling (i.e. lack of sequence data) of relevant prokaryotic taxa, and the ongoing evolution and HGT within and between both archaea, bacteria and eukaryotes^79–84^. Nevertheless, we identified certain lineages within the sister groups, which previously have been suggested to have exchanged genes with early eukaryotes, such as Chlamydiota^45^, and Myxococcota^85^. Together, the diversity of potential donors, highlights the mosaicism of the CCM in LECA, including contributions from additional prokaryotic sources.

The cyanobacterial contributions representing EGTs from the chloroplast to the Archaeplastida ancestor operate in the EMP and PPP (**Fig. 5**), which are connected to the Calvin Cycle^73^. The evolutionary origins of both chloroplast and cytoplasmic versions of the EMP and PPP in Archaeplastida shows a general prevalence of nuclear gene duplications over the genes originating from chloroplast. The predominant process appears to be one in which ‘nuclear’ genes were duplicated, with one copy relocated to the photosynthetic organelle, which might have promoted the genome reduction of the endosymbiont^71,86^. Similarly, while the targeted localization of glycolytic enzymes to the mitochondria has previously led to the suggestion of an endosymbiotic origin of glycolysis ^74,75^, our work does not support this conclusion. Instead our data indicates that CCM enzymes have been retargeted between cytosol, mitochondrion, and plastid many times independently during the evolution of eukaryotes, revealing an ongoing and highly dynamic remodeling of eukaryotic CMM.

Our results show that investigating the origin of the eukaryotic metabolism is crucial to inform our understanding of eukaryogenesis and the impact of the two primary endosymbiotic events that occurred during the origin and diversification of eukaryotes. The archaeal contributions we identify are not consistent with the view that eukaryotic metabolism is predominantly of bacterial origin^34–39^; instead, we show that the CCM is the result of contributions from host and symbiont, with the original pattern partially obscured through remodeling by HGT during subsequent eukaryotic evolution. The observation that most enzymes of archaeal ancestry are cytosolic and operate in the EMP/PPP, while genes of alphaproteobacterial origin function in the TCA within the mitochondrial organelle, support symbiogenetic models of eukaryogenesis: i.e. those models that invoke two partners with an archaeal origin of the eukaryotic cytoplasm and an alphaproteobacterial origin of the mitochondrium^15,21,24,26,28^. While we could not identify a third dominant donor lineage, it seems clear that some CCM enzymes present in LECA have other phylogenetic origins among prokaryotes. Given the lineage-specific remodeling observed in eukaryotic lineages post-LECA, an explanation for this pattern may be that the continuous remodeling process was already underway in “stem” eukaryotes, that is before the radiation of the modern eukaryotic lineages. Our observations are in agreement with hypotheses that emphasise syntrophy between host and endosymbiont, in which the archaeal partner produced reducing equivalents by the degradation of organic substrates via glycolysis which, in the absence of a suitable electron acceptor, were shuttled to a bacterial symbiont which contributed a TCA cycle and an electron transport chain^24–26,30,87^, with the phylogenetic affinities of modern CCM enzymes retaining a vestige of the ancestral contribution from both host and symbiont lineages. Given our results and the previously undetected asgardarchaeal host contributions to the eukaryotic CCM, we expect future studies analyzing the gene origins of additional metabolic pathways, to allow further refining symbiogenetic models on the origin of the eukaryotic cell.

## Methods

### Dataset construction

#### Initial proteome selection, annotation, and redundant filtering: core dataset

We assembled a representative and balanced dataset of selected proteomes comprising 483 archaea, 487 bacteria (5 archaeal and 95 bacterial phyla) and 224 eukaryotic proteomes, that we refer to as *core database* (**Table S1** Supplementary Tables CCM). Each proteome was annotated with eggNOG-mapper v.2.1.4-2 (MMseqs search mode^88,89^), KOFAM_SCAN v.1.3.0^90^ (-f mapper-one-line, -E 1e-3), and HMMSEARCH (HMMER.3.2.3 ^91^, e-value < 1e-3, selecting best i-value hit) against KO.hmm database^92^. We also performed DIAMOND v.2.0.6^93^ protein sequence searches against NCBI_nr release 244. To identify sequences for metabolic gene trees, we primarily used Kegg Orthology (KO) annotations, prioritizing KOFAM classifications. In instances where KOFAM annotation was absent, we relied on HMMSEARCH annotations. The respective sequences were additionally annotated with TargetP v.2.0^94^.

Eukaryotic proteomes were downloaded and manually selected from EukProt v3 ^95^. As this selection includes a variety of sequencing methods (genomes, transcriptomes and single-cell genomes), redundant and truncated sequences were filtered out uniformly. For each proteome, we first used MMseqs2 (options easy-cluster, --cluster-mode 2, --cov-mode 1, -c 1 --min-seq-id 0.95 ^88^), and then, used a custom script (*read_clusters_mmseqs_declusterization.py*) to redefine flawed clusters.

#### Curation of proteomes from eukaryotic contaminations

We performed phylogenies of eukaryotic phylogenetic markers (*see below*) and identified prominent contaminations in the proteomes of some taxa in our dataset (**Table S2**). Among others, these seem to be a result from difficulties in obtaining axenic cultures (eg. Telonemia^96^). To detect and filter out these contaminant sequences, we implemented the following workflow: first, we clustered protein families using Broccoli v1.2.1^97^, using representative nonredundant eukaryotic proteomes. For each orthogroup (OG), we aligned sequences with MAFFT-auto v7.453^98^, and trimmed the MSA with trimAl 1.4.22^99^ (-gt 0.2), removing sequences with coverage below 35% (custom script). We then used FastTree v2.1.11^100^ (-lg) for inferring the phylogeny of each OG. Finally, we used a custom ETE script ^101^ to identify contaminations defined as cases in which certain eukaryotic taxa formed a monophyletic group together with the known contaminants. Specifically, following contaminants were removed: Kinetopalstids sequence data was detected in several eukaryotic proteomes including *Lapot gusevi*, Colponemids and Telonemia among others, and Apusomonadida sequences were detected in proteomes of *Choanocystis* sp. and *Colponema vietnamica*. For kinetoplastid contamination, truncated contaminant sequences remained after this filtering, and thus, we additionally filtered out those sequences that were taxonomically assigned to kinetoplastids given NCBI and EggNog annotations (**Supp table 1.2**).

#### KO homologies

Single KO families are not always sufficient for inferring deep evolutionary history of enzymes because they are sometimes defined on relatively shallow levels. Therefore, we inferred the homology across KO families and combined homologous families when necessary (*see* **Table S3**). We clustered all sequences from the *core dataset* by KO annotation, and further analyzed those KO families with more than 10 sequences. Specifically, sequences for each KO were aligned with MAFFT-auto v7.453^98^, trimmed using trimAl 1.4.22^99^ (-gt 0.35) and again curated with trimAl (-maxidentity 0.85 -seqoverlap 80 -resoverlap 0.5). Next, we made individual HMMs with the HH-suite 3.1.0 package^102^, using HHMAKE (-M 50). We combined all the resulting KO.hhm (14,744) into a single HH-suite database. Then, we performed HHSEARCH of KO.hhm of interest against our HH-suite database (**Table S4**). Finally, we merged those KOs that were relevant for inferring the evolutionary history of certain families.

#### Investigating the origins of LECA clades using expanded dataset

To improve identification of prokaryotic origins of eukaryotic KO families, we searched potential LECA gene families (preliminarily identified from initial trees, *see below*) against a broader set of prokaryotic (NCBI-GTDB) and virus (NCBI) proteomes. We assembled a local dataset including all translated genomes from NCBI that have GTDB^49^ annotation and whose genome completeness was higher than 75% and genome contamination was lower than 5% (a total of 187,681 prokaryotic proteomes which were overrepresented in phyla like Proteobacteria, Firmicutes and Actinobacteria among others; **Table S1**). Additionally, we added viruses from NCBI (a total of 44,889 viral proteomes). We refer to this database as the *expanded dataset.* The workflow was as following: we first screened potential LECA clades across the preliminary phylogenies of CCM enzymes (*see below*) and performed respective HMM protein models using exclusively eukaryotic sequences. Then, we performed HMMSEARCHES (-E 1e-5) of these eukaryotic HMMs against the *expanded prokaryotic and viral datasets*. To avoid overrepresentation of taxa, for each HMMSEARCH we selected the top 15 sequences for each taxonomic class until we collected a total of 150 sequences. Then, we added these sequences to our original set of sequences from the core *dataset* (removing redundant sequences at 97% of identity threshold using cd-hit). These extended searches provided potential donors that were overlooked in the core dataset (eg. LDH phylogeny).

### Phylogenetic analyses

#### Eukaryotic Tree of Life phylogenies

The eukaryotic tree of life was reconstructed by the concatenation of the alignments of phylogenetic markers whose suitability as marker and marker assignment for each species were obtained through iterative phylogenetic reconstruction supervised by hand. We first assembled protein HMMs (MAFFT-auto v7.453^98^, trimAl 1.4.22^99^ -gt 0.4, HMMBUILD^91^) using the sequences for 320 markers provided in^103^. For each phylogenetic marker HMM, we made HMMSEARCH (-E 1e-15) against the eukaryotic proteomes and extracted the top 10 sequences of each taxon sorted by individual-evalue per domain. We performed an initial phylogeny using MAFFt-auto v7.453^98^, trimming with trimAl 1.4.22^99^ (-gt 0.70) and FastTree v2.1.11^100^ (-lg) to identify the orthogroup in question and remove spurious and/or long-branching sequences. Then, we performed two other rounds of phylogenies using MAFFT-L-INS-i v7.453^98^, BMGE 1.12^104^ (-h 0.55), MSA-cover > 35%, and built the gene tree with IQ-TREE 2.1.2 using ultrafast bootstrap with the best fitting empirical or mixture model^105,106^ (-bb 1000 -mset LG -madd LG+C10,LG+C20,LG+C10+R+F,LG+C20+R+F). These two rounds were used to identify and remove contaminating sequences and select a single ortholog per taxon based on the phylogenetic position and the sequence length relative to the total alignment length (note that 3 phylogenetic markers were excluded due to low phylogenetic resolution). We finally concatenated 317 markers that were individually aligned with MAFFT-L-INS-i v7.453^98^ and trimmed with BMGE 1.12^104^ (-h 0.55). Phylogenetic analyses were based on IQ-TREE 2.1.2^106^ (*see below*). Taxa with a concatenation coverage below 50% as well as fast-evolving taxa like Microsporidia were excluded for analyses focusing on the eukaryotic tree of life (**Fig. 1 and Table S1**).

We first reconstructed a phylogeny using corrected UBFoot2 and the LG+C60+G mixture model (-mset LG -madd LG+C60+G --score-diff all -bb 1000 -bnni) with IQ-TREE 2.1.2^105,106^. We then gradually removed heterogeneous sites using *alignment_pruner.pl*script (--chi2_prune 0-0.9, https://github.com/novigit/davinciCode/blob/master/perl), and followed by phylogenetic inferences using IQ-TREE 2.1.2^105,106^ (-mset LG -madd LG+C60 -bb 1000). We additionally reduced the MSA to 148 selected eukaryotes and performed a bayesian phylogeny using PhyloBayes 3^107^ (-catfix C60, -gtr) although the chains did not converged (11,700 generations, max_dif=1, meandif=0.03).

#### Central carbon metabolism enzyme phylogenies

We used the metabolic maps of Glycolysis, Pentose Phosphate pathway, Entner-Doudoroff pathway, Pyruvate metabolisms and Citrate cycle provided by KEGG (https://www.genome.jp/kegg/pathway.html), and determined their distribution across our core dataset to selected those KOs that were present in eukaryotes (**Table S3**). Instances like GAPOR, KGD, KOR among other enzymes, were not found in eukaryotes and excluded in downstream analyses.

To reconstruct refined gene tree phylogenies, we performed three main steps (**Fig. S2**). In the *preliminary phase,* we built an initial and curated phylogeny using the *core dataset*, by using a strict MSA covering threshold (> 80%), visual inspection of the MSA, and removing terminal long branches (longer than six times the mean of all the terminal branch lengths, *read_terminalbranchlength.py*). Final trees were obtained using IQ-TREE 2.1.2 using the best model s and optimized UfBoot2 (-mset LG -madd LG+C20+G+F -bb 1000 -bnni -alrt). Then, we manually inspected the trees and identified those potential LECA clades (including cases like GPI and PGD) to build a eukaryotic-specific HMM. These HMM were then used for the *extension phase* in which we made HMMSEARCHES of each ‘LECA’ HMM against our local NCBI database including prokaryotes and viruses, and add the 150 top sequences to our final set of sequences obtained in the previous phase (*see above, expanded dataset*).

The final phase consisted in two kinds of reconstructions. One was based on the strict trimming (MSA cover > 80%) to get consistent sister group relationship and definition of LECA clades while the other was based on inclusive trimming (MSA cover > 20%) in order to include truncated sequences in the absence/presence profiles across eukaryotes. Final phylogenies are based on Mafft-L-INS-i alignments trimmed with trimAl 1.4.22 (-gt 0.7), and IQ-TREE 2.1.2 using empirical and mixture models (-mset LG | -m LG+C20+G+F -bb 1000 -bnni - alrt -nstop 500 -pers 0.2). Trees were manually rooted as described in supplementary information. We preferentially used outgroup rooting, but when this was not possible, we chose an arbitrary root to ease visualization of sister groups of interest. In addition, some phylogenies required further refinements including addition of an outgroup (GLK, HK, ADPGK, ACO, MDH, LDH, SDH, LSC), extraction and phylogeny of single Pfam domains (PFK, H6PD, PGLS, ACDAB, ACLAB/Y), subselection of sequences for refined phylogenies (TPI, PK, EDD, EDA, TAL, TKT, PDHD, POR, FUMAB/C), and conservation of introns (GPI, PGD, ACS, MDH/LDH). All trees were annotated and visualized with iToL^108^. All alignments and raw tree files can be found on Zenodo repository 10.5281/zenodo.10991068.

#### Domain extraction for phylogenetic reconstructions

For PFK, H6PD and PGLS phylogenies, we extracted the respective Pfam domain of interest inferred with HMMSCAN. For the case of ACLAB/Y and ACDAB, we first built the respective MSAs including all homologs (fused and separate genes) using MAFFT-L-INS-i v7.453 and trimAl 2.1.2 (-gt 0.4). Then, we split the MSA into the respective subunits (ACLA/ACLB, and ACDA/ACDB) and built a HMM with HMMBUILD. Then, we aligned the set of sequences to the HMM using HMMALIGN (--trimm), and converted the *sto* alignment into unaligned sequences which were used for final phylogeny. Similar approach was used for separated mitochondrial pyruvate carrier subunits (MPC1/2). Note that phylogeny of AcdB/AclA subunit in **Fig. S29A** consists of extraction of ATP-grasp Pfam domain. Final phylogenies were conducted as described above.

#### Analyses for shared introns

We investigated the shared intron for GPI, PGD, ACS and MDH/LDH gene families in order to investigate the potential monophyly of eukaryotic sequences. We search the HMM of interest against a set of proteomes previously selected for which genome data is available^109^. Then, we made preliminary trees using MAFFT v7.453 (default), trimAl 1.4.22 (-gt 0.7) and FastTree v2.1.11 (-lg), from which we selected the eukaryotic orthogroups of interest to investigate shared introns. We realigned the selected sequence using MAFFT-L-INS-i v7.453 and used imapper (https://github.com/JulianVosseberg/imapper) to infer the table of shared intron positions. Finally, we made a phylogeny including closely related prokaryotic sequences previously obtained, using MAFFT-L-INS-i v7.453 , trimAl 1.4.22 (-gt 0.7) and IQ-TREE 2.1.2 (-m LG+C20+G+F -B 1000 -alrt 1000 -bnni) and mapped the intron positions with sufficient conservation. For putative single gene families like GPI and PGD we mapped those positions with more than 4 sharing introns in the same phase relative to their codon, while for putative paralogous gene families like ACS and MDH, we mapped those positions that shared intron between two subfamilies, and one of them contain at least 4 taxa sharing an intron.

#### Topology support along GPI’s MSA partitions

In GPI’s MSA, we identified a shared intron between Cyptophyceae and chloroplast clade. We made the phylogeny of 20 positions down and upstream the intron position (at position 1915), and the rest of C-terminus, providing different topologies and suggesting a recombination event between nuclear and plastid paralogs. To verify that topology is specific to the recombined region, we made a subselection of 67 representative sequences, and aligned them L-INS-i v7.453 and trimmed with BMGE 1.12 (-h 0.55). Then, we split the MSA into partitions of 14 positions, in order to span the potential recombined region identified by eye. We made phylogenies of each partition using IQ-TREE (-bb 1000) with LG+G+F and C20+G+F models. Finally, we read the *ufboot files with a custom ETE script, to investigate the Ufboot2 of the topologies of interest: Cryptophyceae+Chlamydia or Cryptophyceae-only sequences branching monophyletically with plastid (Archaeplastida+Cyanobacteria) or with potential LECA paralog.

### Orthogroup definition and correlations

For the definition of orthogroups, we manually selected the sequences forming a monophyletic clade in the respective tree built using inclusive trimming (see above). Note that we also contrast it with strict trimming trees to include sequences that were not correctly placed with eukaryotic sequences, eg. Metamonada in PDHD phylogeny. These manually selected orthogroups were used for plotting the phylogenetic profile (in **Fig 3A**) as well as for plotting the presence of targeting signals (**Fig. 4C**). To infer the correlative distribution of these orthogroups, we converted the orthogroup distribution table into absences (0) and presences (1), and inferred Phi correlation coefficient using *sklearn.metrics.matthews_corrcoef* python function.

## Acknowledgements

A.S. and B.S. have received support from an initiative of Utrecht University (UU) to foster collaborations between UU and NIOZ (NZ4543.11: “The origin and diversification of eukaryotic metabolisms”) and thank additional collaboration partners of this project, Emmanuelle J. Javaux, Paul Mason, and Rick Hennekam. A.S. has received funding from the European Research Council (ERC) under the European Union’s Horizon 2020 research and innovation programme (grant agreement No. 947317, ASymbEL), the Moore–Simons Project on the Origin of the Eukaryotic Cell, Simons Foundation 735929LPI (https://doi.org/10.46714/735929LPI), and a Gordon and Betty Moore Foundation’s Symbiosis in Aquatic Systems Initiative (GBMF9346). T.A.W. and A.S. have received funding by the Gordon and Betty Moore Foundation (GBMF9741). We want to thank Courtney Stairs, Nina Dombrowski, Maximilian Raas, Savvas Tzavellas, and Daniel Tamarit for helpful discussions on eukaryotic and archaeal/bacterial taxon selection, redundancy filtering, intron analysis, and phylogenetic analyses, respectively.

## Author contributions

A.S. and B.S. conceived the study. C.S.M. performed analyses, wrote code, and made figures. C.S.M., A.S., and B.S. analyzed and interpreted data. T.A.W. contributed expertise and helped to interpret results. C.S.M. with B.S. and A.S. wrote the manuscript and supplementary materials and all authors (C.S.M., T.A.W., B.S., A.S.) contributed to the final version of the submitted manuscript.

## Code availability

Workflows for annotations and phylogenies, and custom python scripts to analyze and parse annotation data for figure generation have been deposited in our data repository at Zenodo (10.5281/zenodo.10991067). We used the following published codes: https://github.com/takaram/kofam_scan/tree/master, https://github.com/novigit/davinciCode/blob/master/perl/alignment_pruner.pl, and https://github.com/JulianVosseberg/imapper.

## Data availability

All genomic data of Archaea and Bacteria analyzed are available at NCBI, while all eukaryotic proteomes were downloaded from EukProt V3, and are provided together with the annotations in our data repository at Zenodo (10.5281/zenodo.10991067). Data generated in this study including single gene tree analyses, concatenated phylogenies and manual annotations (i.e. sequence files, alignments, and treefiles, compositions of orthogroups and sister groups, etc) have also been deposited in our data repository at Zenodo (10.5281/zenodo.10991067). Public databases are available as follows: EggNog annotations were obtained with Eggnog-mapper 2.1.4-2 (https://github.com/eggnogdb/eggnog-mapper), KOFAM annotations and KO profiles downloaded from the KEGG Automatic Annotation Server in 2021 (https://www.genome.jp/tools/kofamkoala/), the NCBI proteomes were downloaded in November 2021 (https://ftp.ncbi.nlm.nih.gov/genomes/) using taxonomic annotations from GTDB (https://data.gtdb.ecogenomic.org/) and eukaryotic proteomes were downloaded from EukProt V3 (https://doi.org/10.6084/m9.figshare.12417881.v3).

## Ethics statement

This research did not involve animals or humans and no new data has been generated. Furthermore, the information provided here does not pose a threat to public health, safety or security, animals, plants or the environment.

## References

1. Parfrey, L. W., Lahr, D. J. G., Knoll, A. H. & Katz, L. A. Estimating the timing of early eukaryotic diversification with multigene molecular clocks. Proc. Natl. Acad. Sci. U. S. A. 108, 13624–13629 (2011).

2. Eme, L., Sharpe, S. C., Brown, M. W. & Roger, A. J. On the Age of Eukaryotes: Evaluating Evidence from Fossils and Molecular Clocks. Cold Spring Harb. Perspect. Biol. 6, a016139 (2014).

3. Betts, H. C. et al. Integrated genomic and fossil evidence illuminates life’s early evolution and eukaryote origin. *Nat*. Ecol. Evol. 2, 1556–1562 (2018).

4. Gueneli, N. et al. 1.1-billion-year-old porphyrins establish a marine ecosystem dominated by bacterial primary producers. Proc. Natl. Acad. Sci. U. S. A. 115, E6978–E6986 (2018).

5. Mahendrarajah, T. A. et al. ATP synthase evolution on a cross-braced dated tree of life. Nat. Commun. 14, 7456 (2023).

6. Lyons, T. W., Reinhard, C. T. & Planavsky, N. J. The rise of oxygen in Earth’s early ocean and atmosphere. Nature 506, 307–315 (2014).

7. Mills, D. B. et al. Eukaryogenesis and oxygen in Earth history. *Nat*. Ecol. Evol. 6, 520–532 (2022).

8. Craig, J. M., Kumar, S. & Hedges, S. B. The origin of eukaryotes and rise in complexity were synchronous with the rise in oxygen. Front. Bioinforma. 3, (2023).

9. Spang, A. et al. Complex archaea that bridge the gap between prokaryotes and eukaryotes. Nature 521, 173–179 (2015).

10. Zaremba-Niedzwiedzka, K. et al. Asgard archaea illuminate the origin of eukaryotic cellular complexity. Nature 541, 353–358 (2017).

11. Liu, Y. et al. Expanded diversity of Asgard archaea and their relationships with eukaryotes. Nature 593, 553–557 (2021).

12. Eme, L. et al. Inference and reconstruction of the heimdallarchaeial ancestry of eukaryotes. Nature 618, 992–999 (2023).

13. Roger, A. J., Muñoz-Gómez, S. A. & Kamikawa, R. The Origin and Diversification of Mitochondria. Curr. Biol. 27, R1177–R1192 (2017).

14. Martijn, J., Vosseberg, J., Guy, L., Offre, P. & Ettema, T. J. G. Deep mitochondrial origin outside the sampled alphaproteobacteria. Nature 557, 101–105 (2018).

15. Martin, W. & Müller, M. The hydrogen hypothesis for the first eukaryote. Nature 392, 37–41 (1998).

16. Cavalier-Smith, T. The phagotrophic origin of eukaryotes and phylogenetic classification of Protozoa. Int. J. Syst. Evol. Microbiol. 52, 297–354 (2002).

17. Martijn, J. & Ettema, T. J. G. From archaeon to eukaryote: the evolutionary dark ages of the eukaryotic cell. Biochem. Soc. Trans. 41, 451–457 (2013).

18. Baum, D. A. & Baum, B. An inside-out origin for the eukaryotic cell. BMC Biol. 12, 76 (2014).

19. Guy, L., Saw, J. H. & Ettema, T. J. G. The archaeal legacy of eukaryotes: a phylogenomic perspective. Cold Spring Harb. Perspect. Biol. 6, a016022 (2014).

20. Wang, Z. & Wu, M. Phylogenomic reconstruction indicates mitochondrial ancestor was an energy parasite. PloS One 9, e110685 (2014).

21. Koonin, E. V. Origin of eukaryotes from within archaea, archaeal eukaryome and bursts of gene gain: eukaryogenesis just made easier? Philos. Trans. R. Soc. B Biol. Sci. 370, 20140333 (2015).

22. Martin, W. F., Garg, S. & Zimorski, V. Endosymbiotic theories for eukaryote origin. Philos. Trans. R. Soc. B Biol. Sci. 370, 20140330 (2015).

23. Moreira, D. & López-García, P. Evolution of viruses and cells: do we need a fourth domain of life to explain the origin of eukaryotes? Philos. Trans. R. Soc. B Biol. Sci. 370, (2015).

24. Spang, A. et al. Proposal of the reverse flow model for the origin of the eukaryotic cell based on comparative analyses of Asgard archaeal metabolism. Nat. Microbiol. 4, 1138–1148 (2019).

25. Imachi, H. et al. Isolation of an archaeon at the prokaryote–eukaryote interface. Nature 577, 519–525 (2020).

26. López-García, P. & Moreira, D. The Syntrophy hypothesis for the origin of eukaryotes revisited. Nat. Microbiol. 5, 655–667 (2020).

27. Speijer, D. Debating Eukaryogenesis-Part 1: Does Eukaryogenesis Presuppose Symbiosis Before Uptake? BioEssays News Rev. Mol. Cell. Dev. Biol. 42, e1900157 (2020).

28. Donoghue, P. C. J. et al. Defining eukaryotes to dissect eukaryogenesis. Curr. Biol. 33, R919–R929 (2023).

29. Dacks, J. B. et al. The changing view of eukaryogenesis - fossils, cells, lineages and how they all come together. J. Cell Sci. 129, 3695–3703 (2016).

30. Sousa, F. L., Neukirchen, S., Allen, J. F., Lane, N. & Martin, W. F. Lokiarchaeon is hydrogen dependent. Nat. Microbiol. 1, 16034 (2016).

31. Martin, W. & Koonin, E. V. Introns and the origin of nucleus–cytosol compartmentalization. Nature 440, 41–45 (2006).

32. Burns, J. A., Pittis, A. A. & Kim, E. Gene-based predictive models of trophic modes suggest Asgard archaea are not phagocytotic. *Nat*. Ecol. Evol. 2, 697–704 (2018).

33. Baum, B. & Spang, A. On the origin of the nucleus: a hypothesis. Microbiol. Mol. Biol. Rev. MMBR 87, e0018621 (2023).

34. Jain, R., Rivera, M. C. & Lake, J. A. Horizontal gene transfer among genomes: The complexity hypothesis. Proc. Natl. Acad. Sci. 96, 3801–3806 (1999).

35. McInerney, J. O., O’Connell, M. J. & Pisani, D. The hybrid nature of the Eukaryota and a consilient view of life on Earth. Nat. Rev. Microbiol. 12, 449–455 (2014).

36. Rochette, N. C., Brochier-Armanet, C. & Gouy, M. Phylogenomic Test of the Hypotheses for the Evolutionary Origin of Eukaryotes. Mol. Biol. Evol. 31, 832–845 (2014).

37. Pittis, A. A. & Gabaldón, T. Late acquisition of mitochondria by a host with chimaeric prokaryotic ancestry. Nature 531, 101–104 (2016).

38. Méheust, R. et al. Formation of chimeric genes with essential functions at the origin of eukaryotes. BMC Biol. 16, 30 (2018).

39. Knopp, M., Stockhorst, S., van der Giezen, M., Garg, S. G. & Gould, S. B. The Asgard Archaeal-Unique Contribution to Protein Families of the Eukaryotic Common Ancestor Was 0.3. Genome Biol. Evol. 13, evab085 (2021).

40. Canback, B., Andersson, S. G. E. & Kurland, C. G. The global phylogeny of glycolytic enzymes. Proc. Natl. Acad. Sci. 99, 6097–6102 (2002).

41. Schnarrenberger, C. & Martin, W. Evolution of the enzymes of the citric acid cycle and the glyoxylate cycle of higher plants. A case study of endosymbiotic gene transfer. Eur. J. Biochem. 269, 868–883 (2002).

42. Szklarczyk, R. & Huynen, M. A. Mosaic origin of the mitochondrial proteome. PROTEOMICS 10, 4012–4024 (2010).

43. Alsmark, C. et al. Patterns of prokaryotic lateral gene transfers affecting parasitic microbial eukaryotes. Genome Biol. 14, R19 (2013).

44. Stairs, C. W. et al. Microbial eukaryotes have adapted to hypoxia by horizontal acquisitions of a gene involved in rhodoquinone biosynthesis. eLife 7, e34292 (2018).

45. Stairs, C. W. et al. Chlamydial contribution to anaerobic metabolism during eukaryotic evolution. Sci. Adv. 6, eabb7258 (2020).

46. Hug, L. A. et al. A new view of the tree of life. Nat. Microbiol. 1, 16048 (2016).

47. Castelle, C. J. & Banfield, J. F. Major New Microbial Groups Expand Diversity and Alter our Understanding of the Tree of Life. Cell 172, 1181–1197 (2018).

48. Burki, F., Roger, A. J., Brown, M. W. & Simpson, A. G. B. The New Tree of Eukaryotes. Trends Ecol. Evol. 35, 43–55 (2020).

49. Parks, D. H. et al. GTDB: an ongoing census of bacterial and archaeal diversity through a phylogenetically consistent, rank normalized and complete genome-based taxonomy. Nucleic Acids Res. 50, D785–D794 (2022).

50. Hampl, V. et al. Phylogenomic analyses support the monophyly of Excavata and resolve relationships among eukaryotic “supergroups”. Proc. Natl. Acad. Sci. 106, 3859–3864 (2009).

51. Rogozin, I. B., Basu, M. K., Csürös, M. & Koonin, E. V. Analysis of rare genomic changes does not support the unikont-bikont phylogeny and suggests cyanobacterial symbiosis as the point of primary radiation of eukaryotes. Genome Biol. Evol. 1, 99–113 (2009).

52. He, D. et al. An Alternative Root for the Eukaryote Tree of Life. Curr. Biol. 24, 465–470 (2014).

53. Cerón-Romero, M. A., Fonseca, M. M., de Oliveira Martins, L., Posada, D. & Katz, L. A. Phylogenomic Analyses of 2,786 Genes in 158 Lineages Support a Root of the Eukaryotic Tree of Life between Opisthokonts and All Other Lineages. Genome Biol. Evol. 14, evac119 (2022).

54. Al Jewari, C. & Baldauf, S. L. An excavate root for the eukaryote tree of life. Sci. Adv. 9, eade4973 (2023).

55. Gray, M. W. et al. The draft nuclear genome sequence and predicted mitochondrial proteome of Andalucia godoyi, a protist with the most gene-rich and bacteria-like mitochondrial genome. BMC Biol. 18, 22 (2020).

56. Pyrih, J. et al. Vestiges of the Bacterial Signal Recognition Particle-Based Protein Targeting in Mitochondria. Mol. Biol. Evol. 38, 3170–3187 (2021).

57. Galindo, L. J., Prokina, K., Torruella, G., López-García, P. & Moreira, D. Maturases and Group II Introns in the Mitochondrial Genomes of the Deepest Jakobid Branch. Genome Biol. Evol. 15, evad058 (2023).

58. Stairs, C. W. et al. Anaeramoebae are a divergent lineage of eukaryotes that shed light on the transition from anaerobic mitochondria to hydrogenosomes. Curr. Biol. 31, 5605–5612.e5 (2021).

59. Tikhonenkov, D. V. et al. Microbial predators form a new supergroup of eukaryotes. Nature 612, 714–719 (2022).

60. Chen, X. et al. The Entner–Doudoroff pathway is an overlooked glycolytic route in cyanobacteria and plants. Proc. Natl. Acad. Sci. 113, 5441–5446 (2016).

61. Gawryluk, R. M. R., Eme, L. & Roger, A. J. Gene fusion, fission, lateral transfer, and loss: Not-so-rare events in the evolution of eukaryotic ATP citrate lyase. Mol. Phylogenet. Evol. 91, 12–16 (2015).

62. Stairs, C. W., Leger, M. M. & Roger, A. J. Diversity and origins of anaerobic metabolism in mitochondria and related organelles. Philos. Trans. R. Soc. Lond. B. Biol. Sci. 370, 20140326 (2015).

63. Karnkowska, A. et al. A Eukaryote without a Mitochondrial Organelle. Curr. Biol. 26, 1274–1284 (2016).

64. Novák, L. V. F. et al. Genomics of Preaxostyla Flagellates Illuminates the Path Towards the Loss of Mitochondria. PLOS Genet. 19, e1011050 (2023).

65. Verhees, C. H. et al. The unique features of glycolytic pathways in Archaea. Biochem. J. 375, 231–246 (2003).

66. Gietl, C. Malate dehydrogenase isoenzymes: cellular locations and role in the flow of metabolites between the cytoplasm and cell organelles. Biochim. Biophys. Acta 1100, 217–234 (1992).

67. More, K., Klinger, C. M., Barlow, L. D. & Dacks, J. B. Evolution and Natural History of Membrane Trafficking in Eukaryotes. Curr. Biol. 30, R553–R564 (2020).

68. Martin, W. F. Is something wrong with the tree of life? BioEssays 18, 523–527 (1996).

69. Stechmann, A., Baumgartner, M., Silberman, J. D. & Roger, A. J. The glycolytic pathway of Trimastix pyriformis is an evolutionary mosaic. BMC Evol. Biol. 6, 101 (2006).

70. Grauvogel, C., Brinkmann, H. & Petersen, J. Evolution of the Glucose-6-Phosphate Isomerase: The Plasticity of Primary Metabolism in Photosynthetic Eukaryotes. Mol. Biol. Evol. 24, 1611–1621 (2007).

71. Martin, W. Evolutionary origins of metabolic compartmentalization in eukaryotes. Philos. Trans. R. Soc. B Biol. Sci. 365, 847–855 (2010).

72. A. von der Dunk, S. H. & Snel, B. Recurrent sequence evolution after independent gene duplication. BMC Evol. Biol. 20, 98 (2020).

73. Maeda, H. A. & Fernie, A. R. Evolutionary History of Plant Metabolism. Annu. Rev. Plant Biol. 72, 185–216 (2021).

74. Río Bártulos, C., et al. Mitochondrial Glycolysis in a Major Lineage of Eukaryotes. Genome Biol. Evol. 10, 2310–2325 (2018).

75. Liaud, M. F., Lichtlé, C., Apt, K., Martin, W. & Cerff, R. Compartment-specific isoforms of TPI and GAPDH are imported into diatom mitochondria as a fusion protein: evidence in favor of a mitochondrial origin of the eukaryotic glycolytic pathway. Mol. Biol. Evol. 17, 213–223 (2000).

76. Hannaert, V. et al. Enolase from Trypanosoma brucei, from the Amitochondriate Protist Mastigamoeba balamuthi, and from the Chloroplast and Cytosol of Euglena gracilis: Pieces in the Evolutionary Puzzle of the Eukaryotic Glycolytic Pathway. Mol. Biol. Evol. 17, 989–1000 (2000).

77. Martin, W. & Russell, M. J. On the origins of cells: a hypothesis for the evolutionary transitions from abiotic geochemistry to chemoautotrophic prokaryotes, and from prokaryotes to nucleated cells. Philos. Trans. R. Soc. Lond. B. Biol. Sci. 358, 59–83; discussion 83-85 (2003).

78. Freibert, S.-A. et al. Evolutionary conservation and in vitro reconstitution of microsporidian iron–sulfur cluster biosynthesis. Nat. Commun. 8, 13932 (2017).

79. Archibald, J. M. Gene transfer in complex cells. Nature 524, 423–424 (2015).

80. Ku, C. et al. Endosymbiotic origin and differential loss of eukaryotic genes. Nature 524, 427–432 (2015).

81. Leger, M. M., Eme, L., Stairs, C. W. & Roger, A. J. Demystifying Eukaryote Lateral Gene Transfer (Response to Martin 2017 DOI: 10.1002/bies.201700115). BioEssays 40, 1700242 (2018).

82. Wu, F. et al. Unique mobile elements and scalable gene flow at the prokaryote–eukaryote boundary revealed by circularized Asgard archaea genomes. Nat. Microbiol. 7, 200–212 (2022).

83. Filée, J. et al. Bacterial origins of thymidylate metabolism in Asgard archaea and Eukarya. Nat. Commun. 14, 838 (2023).

84. Keeling, P. J. Horizontal gene transfer in eukaryotes: aligning theory with data. Nat. Rev. Genet. 1–15 (2024) doi:10.1038/s41576-023-00688-5.

85. Santana-Molina, C., Rivas-Marin, E., Rojas, A. M. & Devos, D. P. Origin and Evolution of Polycyclic Triterpene Synthesis. Mol. Biol. Evol. 37, 1925–1941 (2020).

86. Coale, T. H. et al. Nitrogen-fixing organelle in a marine alga. Science 384, 217–222 (2024).

87. Speijer, D. Alternating terminal electron-acceptors at the basis of symbiogenesis: How oxygen ignited eukaryotic evolution. BioEssays 39, 1600174 (2017).

88. Steinegger, M. & Söding, J. MMseqs2 enables sensitive protein sequence searching for the analysis of massive data sets. Nat. Biotechnol. 35, 1026–1028 (2017).

89. Huerta-Cepas, J. et al. eggNOG 5.0: a hierarchical, functionally and phylogenetically annotated orthology resource based on 5090 organisms and 2502 viruses. Nucleic Acids Res. 47, D309–D314 (2019).

90. Aramaki, T. et al. KofamKOALA: KEGG Ortholog assignment based on profile HMM and adaptive score threshold. Bioinformatics 36, 2251–2252 (2020).

91. Potter, S. C. et al. HMMER web server: 2018 update. Nucleic Acids Res. 46, W200–W204 (2018).

92. Kanehisa, M., Sato, Y., Kawashima, M., Furumichi, M. & Tanabe, M. KEGG as a reference resource for gene and protein annotation. Nucleic Acids Res. 44, D457–462 (2016).

93. Buchfink, B., Reuter, K. & Drost, H.-G. Sensitive protein alignments at tree-of-life scale using DIAMOND. Nat. Methods 18, 366–368 (2021).

94. Almagro Armenteros, J. J., et al. Detecting sequence signals in targeting peptides using deep learning. Life Sci. Alliance 2, e201900429 (2019).

95. Richter, D. J. et al. EukProt: A database of genome-scale predicted proteins across the diversity of eukaryotes. Peer Community J. 2, (2022).

96. Strassert, J. F. H., Jamy, M., Mylnikov, A. P., Tikhonenkov, D. V. & Burki, F. New Phylogenomic Analysis of the Enigmatic Phylum Telonemia Further Resolves the Eukaryote Tree of Life. Mol. Biol. Evol. 36, 757–765 (2019).

97. Derelle, R., Philippe, H. & Colbourne, J. K. Broccoli: Combining Phylogenetic and Network Analyses for Orthology Assignment. Mol. Biol. Evol. 37, 3389–3396 (2020).

98. Katoh, K. & Standley, D. M. MAFFT multiple sequence alignment software version 7: improvements in performance and usability. Mol. Biol. Evol. 30, 772–780 (2013).

99. Capella-Gutiérrez, S., Silla-Martínez, J. M. & Gabaldón, T. trimAl: a tool for automated alignment trimming in large-scale phylogenetic analyses. Bioinforma. Oxf. Engl. 25, 1972–1973 (2009).

100. Price, M. N., Dehal, P. S. & Arkin, A. P. FastTree 2--approximately maximum-likelihood trees for large alignments. PloS One 5, e9490 (2010).

101. Huerta-Cepas, J., Dopazo, J. & Gabaldón, T. ETE: a python Environment for Tree Exploration. BMC Bioinformatics 11, 24 (2010).

102. Steinegger, M. et al. HH-suite3 for fast remote homology detection and deep protein annotation. BMC Bioinformatics 20, 473 (2019).

103. Strassert, J. F. H., Irisarri, I., Williams, T. A. & Burki, F. A molecular timescale for eukaryote evolution with implications for the origin of red algal-derived plastids. Nat. Commun. 12, 1879 (2021).

104. Criscuolo, A. & Gribaldo, S. BMGE (Block Mapping and Gathering with Entropy): a new software for selection of phylogenetic informative regions from multiple sequence alignments. BMC Evol. Biol. 10, 210 (2010).

105. Hoang, D. T., Chernomor, O., von Haeseler, A., Minh, B. Q. & Vinh, L. S. UFBoot2: Improving the Ultrafast Bootstrap Approximation. Mol. Biol. Evol. 35, 518–522 (2018).

106. Minh, B. Q. et al. IQ-TREE 2: New Models and Efficient Methods for Phylogenetic Inference in the Genomic Era. Mol. Biol. Evol. 37, 1530–1534 (2020).

107. Lartillot, N., Lepage, T. & Blanquart, S. PhyloBayes 3: a Bayesian software package for phylogenetic reconstruction and molecular dating. Bioinformatics 25, 2286–2288 (2009).

108. Letunic, I. & Bork, P. Interactive Tree Of Life (iTOL) v5: an online tool for phylogenetic tree display and annotation. Nucleic Acids Res. 49, W293–W296 (2021).

109. Vosseberg, J., Schinkel, M., Gremmen, S. & Snel, B. The spread of the first introns in proto-eukaryotic paralogs. *Commun*. Biol. 5, 1–9 (2022).

