## Supplementary Information for "Chimeric Origins and Dynamic Evolution of Central Carbon Metabolism in Eukaryotes"

**Abstract**

The origin of eukaryotes was a key event in the history of life. Current leading hypotheses propose that a symbiosis between an asgardarchaeal host cell and an alphaproteobacterial endosymbiont represented a crucial step in eukaryotic origins and invoke a central role for syntrophic interactions - that is, metabolic cross-feeding - between the partners as the basis for their subsequent evolutionary integration. A major unanswered question is whether the metabolism of modern eukaryotes bears any vestige of this ancestral interaction. To investigate this question in detail, we systematically analyze the evolutionary origins of the eukaryotic gene repertoires mediating central carbon metabolism. Our phylogenetic and sequence analyses reveal that this gene repertoire is chimeric, with ancestral contributions from Asgardarchaeota and Alphaproteobacteria operating predominantly in glycolysis and TCA, respectively. Furthermore, our analyses reveal diverse additional contributions from other prokaryotic sources as well as the extent to which this ancestral metabolic interplay has been remodeled via gene loss, transfer, and subcellular retargeting in the >2Ga since the origin of eukaryotic cells. Together, our work demonstrates that in contrast to previous assumptions, the eukaryotic metabolism preserves information about the nature of the original asgardarchaeal-alphaproteobacterial interactions, and supports syntrophy scenarios on the origin of the eukaryotic cell.

### Central carbon metabolism

Central carbon metabolism (CCM) is a pivotal process in eukaryotic cells providing building blocks for the cell as well as electrons that are channeled into the respiratory chain for energy conservation and biosynthesis. The eukaryotic CCM, as defined here, comprises four main pathways: the Embden–Meyerhof–Parnas glycolytic pathway (canonical glycolysis; EMP; (Kresge et al. 2005)), the pentose phosphate pathway (PPP, including also the Entner-Doudoroff glycolytic pathway; EDP; (Horecker 2002)), pyruvate/acetate conversions and the tricarboxylic acid (TCA, or Krebs(Krebs and Johnson 1937)) cycle. Of specific relevance with regard to eukaryogenesis models is the observation that Glycolysis and PPP, which are involved in the metabolisms of glucose, ribose and pyruvate among other important metabolites, take place in the cytoplasm and in the chloroplast in photosynthetic organisms (Plaxton 1996; Río Bártulos et al. 2018a). By contrast, pyruvate conversions and the TCA cycle generally operate in the mitochondria being efficiently coupled to the respiratory chain (oxidative phosphorylation) which, in aerobic eukaryotes, uses oxygen as final electron acceptor. Several anaerobic protists harbor mitochondria-related organelles (MROs) which originated through the reductive evolution of mitochondria as an adaptation to an anaerobic lifestyle (Stairs et al. 2015; Burki 2016; Stairs et al. 2021; Novák et al. 2023). This resulted in the loss of metabolic processes that take place in the mitochondrion like TCA and pyruvate conversions as well as the use of different cofactors and electron acceptors (Müller et al. 2012; Stairs et al. 2015; Karnkowska et al. 2016; Gawryluk and Stairs 2021).

### Phylogenomic analyses of the eukaryotic tree of life

We performed phylogenomic analyses by concatenating 317 phylogenetic markers (Strassert et al. 2021) in 207 eukaryotes. Our taxon selection spans all known eukaryotic diversity with a balanced taxonomic composition and includes recently sequenced organisms such as early branching Cryptista (*Hemiarma marina*) and Metamonada (Anaeromoebae) among others. We performed a first maximum-likelihood (ML) reconstruction on the concatenated dataset, with IQ-TREE (Minh et al. 2020) using LG+C60+G mixture model and corrected ultrafast bootstraps (UfBoot2) (**Figure 1, S1A**). In agreement with previous studies (Hampl et al. 2009), the phylogeny retrieved Excavata as monophyletic (91%) , full support for Diaphoretickes monophyly (100%), and moderate support for Amorphea as supergroup (77%, **Figure 1**). Note that Malawimonadida and Ancyromonadida branch sister to Amorphea (61%), though their phylogenetic location was unstable across other reconstructions (see below). We also constructed a Bayesian (BY) phylogeny using PhyloBayes (Lartillot et al. 2009) with a reduced set of eukaryotes but the analysis did not converge (C60, -gtr, max_dif = 1 and meandiff = 0.03 after 11,700 generations, **Figure S1B**). Despite the lack of convergence, ML and BY phylogenies provided very similar topologies except for Anaeromoebae taxa, which branched with Amorphea taxa in our BY reconstruction.

In addition, we iteratively removed heterogeneous sites (from 0.1 to 0.9 cut-offs), and built the respective phylogenies using UfBoot2 (Hoang et al. 2018) (**Figure S1C**). We then made a survey of the UfBoot2 values for the monophyly of major eukaryotic groups (Excavata, Amorphea excluding Malawimonadida and Ancyromonadida, and Diaphoretickes, upper panel in **Figure 1B**), and for the monophyly of Malawimonadida and/or Ancyromonadida with the major groups (lower panel in **Figure 1B**). The analysis consistently recovers the Diaphoretickes and Excavata clades, while the definition of Amorphea is less clear. Ancyromonadida shows a monophyletic relationship with Diaphoretickes (0.4 cut-off, 76% Ufboot2). However, the relationship of Ancyromonadida and Malawimonadida with Amorphea had better support (0.4 cut-off, 97% UfBoot2 for Malawimonadida and 0.6 cut-off, 90% UfBoot2 for Malawimonadida and Ancyromonadida), supporting the phylogenetic association of these. The phylogenetic association of Ancyromonadida with Excavata (Tikhonenkov et al. 2022), was not recovered in our analyses. Therefore, as previously pointed out (Atkins et al. 2000; Heiss et al. 2018), Amorphea might be composed of Obozoa, Amoebozoa, CRuMs, Malawimonadida and Ancyromonaida. However, the placement of the two latter lineages was unstable which might be due to compositional biases and a lack of closely related taxa in our analyses.

**Evolution of Embden-Meyerhof-Parnas (EMP) glycolytic enzymes**

**Glucose Phosphorylation**

The first step in the glycolysis is the phosphorylation of the glucose carried out by P-loop NTPases superfamily such as Glucokinase (GLK), Hexokinase (HK) (**Figure S3A-E**; (Stoddard et al. 2020)). Phylogenetic reconstruction combining homologous of all these protein families, plus other closely related ones, shows that GLK, ROK and HK form three distinct families within the large P-loop NTPase superfamily phylogeny, with HK being the most divergent (**Figure S3F**). The GLK/ROK subfamily appears to be the most widespread group in prokaryotes, while HK has a more restricted distribution in Bacteria (**Figure S3A-C**). Besides, glucose can also be phosphorylated by the non-homologous ADP-dependent glucokinase (ADPGK), which belongs to the ADP-dependent phosphofructokinase/glucokinase superfamily (**Figure S4**, PfkC is involved in prokaryotic glycolysis and pentose phosphate pathway (PPP) (Verhees et al. 2003)). Based on the phylogenetic clustering of eukaryotic and prokaryotic sequences described below, we infer that the eukaryotic GLK, ROK, and HK appear to be of bacterial origin, while eukaryotic ADPGK seems to be of archaeal origin.

**Glucokinase and Repressor ORF Kinase — GLK/ROK — K00844**

Although GLK and ROK do not form a monophyletic cluster in the P-loop NTPase superfamily phylogeny, they are assigned to the same orthologous family, K00844. This allowed us to reciprocally root the GLK and ROK family trees, respectively (**Figure S3A,B**).

The internal branches in the GLK family phylogeny are poorly supported, with eukaryotic homologs being separated into distinct clusters. In one of these clusters, a small number of eukaryotic sequences mainly from Excavata and Amoebozoa, place sister to among others DPANN and Spirochaetes. However it is unclear whether these eukaryotic sequences represent a LECA family or have been acquired later on and transferred between eukaryotes. The other major cluster mainly comprises Diaphoretickes branching next to a large bacterial clade including Cyanobacteria and Alphaproteobacteria among others (**Figure S3A**). We also found two basally branching Metamonoda sequences, but considering that Metamonada sequences often form long branches in phylogenetic reconstructions (eg. in the eukaryotic tree of life; **Figure 1**), it is possible that this placement is an artifact. Irrespective, the phylogeny of GLK shows two taxonomically non-overlapping clades, which suggest two potential post-LECA donations, one of them most likely in Diaphoretickes ancestor. However, due to low overall support of this GLK phylogeny and complexity of sister groups, it is not possible to discern the donor of the respective eukaryotic GLKs. Please note that while we found Asgard ROK branching basally to the GLK family (**Figure S3F**), this signal was not recovered in alternative reconstructions including only GLK-ROK (**Figure S3E**) further emphasizing the phylogenetic instability of these GLKs.

In contrast to the complex history of the GLK subfamily, the eukaryotic ROK subfamily forms a monophyletic group, with deep branches being largely consistent with the eToL topology (**Figure**

**S3B**) indicating that it has been present in LECA. Note that this enzyme is not found in well studied eukaryotes such as bilateria, embryophytes, ascomycetes, hence the biological role of the ROK enzyme in eukaryotic biochemistry is unclear. Basally to this potential LECA group, we found alphaproteobacterial sequences from Sphingomonadales and Caulobacterales and a few other bacteria, pointing to a putative alphaproteobacterial origin, but the scarcity of these sequences in extant Alphaproteobacteria limits the certainty of this inference.

**Hexokinase — HK — K00845**

The hexokinase phylogeny (K00845) was rooted with Firmicutes including Clostridia, Negativicutes, Desulfotomaculia and Synthrophomonadia (**Figure S3D**) as suggested by the extended phylogeny of the P-loop NTPase superfamily (**Figure S3F**) and shows a mostly monophyletic group of eukaryotes that separates Amorphea and Diaphoretickes, with few exceptions (**Figure S3D**). Therefore, it is likely that this Hexokinase family was present in LECA. Note however, that a clade comprising Metamonada clusters with a mixed bacterial clade either within this larger eukaryotic LECA clade (**Figure S3D**) or basal to it (protein family reconstruction, asterisk in **Figure S3F**). This suggests an LBA artifact or alternatively independent acquisition. While we cannot pinpoint a specific donor lineage, this family likely has bacterial origins with no indication for the acquisition from Asgardarchaeota nor Alphaproteobacteria.

**ADP-dependent glucokinase — ADPGK — K08074**

In contrast to GLK/ROK, HK and PPGK which belong to the P-loop NTPase superfamily, ADPGK belongs to a distinct protein family called the ribokinase-like superfamily. The closest KO to the eukaryotic ADPGK is the PFKC (ADP-dependent phosphofructokinase/ glucokinase, K00918), which was added to the ADPGK set for the final reconstruction (including sequences from extended searches, **Figure S4**). ADPGK/PFKC is a bifunctional enzyme that can phosphorylate glucose as well as fructose-6-P (first and third steps of the glycolysis; (Verhees et al. 2003)), and was previously suggested to be acquired in Metazoa by HGT from archaea (Ronimus and Morgan 2004). Eukaryotic ADPGK (with the unique K08074) form a monophyletic group including Amophea and Excavate taxa plus other Diaphoretickes protists, suggesting that ADPGK was present in LECA (**Figure S4**) and the respective sister group is composed of two Heimdallarchaeota representatives (moderate support of 76% optimized UfBoot2 ). In turn, considering our phylogenetic data and the widespread distribution of this enzyme in archaea, we suggest an archaeal origin of this family in eukaryotes. However, please note that PfkC evolution was characterized by gene transfers to bacteria including to the gut alphaproteobacterial RF32 lineage, among others (**Figure S4**).

**Glucose-6-phosphate isomerase — GPI — K01810**

The phylogeny of the glucose-6-phosphate isomerase, GPI, is in agreement with previous reconstructions based on a smaller number of sequences (Stechmann et al. 2006; Grauvogel et al. 2007) (**Figure S5A**) and recovers three main distinct eukaryotic clades (described in same order as in **Figure S5B**). The first group comprises anaerobic eukaryotes from the Excavates for which there is no clear sister group (**Figure S5B**). Closely related to this group, we found a bacterial group including Cyanobacteria and photosynthetic unicellular eukaryotes from Streptophyta, with Gloemargarita as a sister taxa, in agreement with a primary EGT and supporting previous observations (Grauvogel et al. 2007). The third group is formed by diverse paraphyletic eukaryotic subgroups which are interspersed with bacterial sequences, including Alpha- and Gammaproteobacteria and Magnetococcia, among other bacteria. One of these subgroups is formed exclusively by Diaphoretickes taxa, being a well defined phylogenetic subgroup. This subgroup includes photosynthetic and non-photosynthetic eukaryotes and shows some secondary EGTs like in Euglenozoa and Ochrophyta. There is another main subgroup comprising Amorphea and Excavata and others composed of Haptista, Cryptista, SAR, Amoebozoa organisms. Additionally, there is another intermediary group including Cryptista branching sister to Chlamydiae and other bacteria (**Figure S5B**).

These paraphyletic groups may be a result of independent origins through HGTs from bacteria. Alternatively, they may indicate the ancestral presence of GPI in LECA combined with HGT with prokaryotes (during and after eukaryogenesis). The large diversity of sequences from Alphaproteobacteria, Gammaproteobacteria and Magnetococcia, branching in-between the Diaphoretickes and the Amorphea-Excavate-Diaphoreticke groups may suggest that these eukaryotic clades represent a LECA clade with a mitochondrial origin as previously pointed out (Grauvogel et al. 2007). Note however, that this phylogeny contains another distant cluster of Proteobacteria (**Figure S5A**), suggesting diverse origins of GPI, even within this bacterial lineage.

One possible way to test the monophyly of eukaryotic sequences is by investigating shared introns between potential monophyletic sequences. Investigating the distribution of introns across the multiple sequence alignment (MSA) of GPI (using a previously generated dataset; (Vosseberg et al. 2022)), we detect two intron positions shared between Diaphoretickes and Opisthokonta which would be in agreement with the monophyly of these eukaryotic sequences (positions 2117 and 2146; (**Figure S5C**). We found three additional introns with one position displaced that could represent remnants of LECA introns (1554/55, 1772/73 and 1979/80) (**Figure S5C**). We also observed potential parallel insertions like those in positions 1915, 2282 and 2373 between the potential LECA and chloroplast paralog. The shared introns in 1915 position, in combination with the intermediary placement of Cryptophyceae-Chlamydia group in the phylogeny (**Figure S5C**), was suspicious and led us to build phylogenies of 20 amino acids up and downstream of the 1915 position and the rest of the C-terminus. This resulted in two different topologies (**Figure S5D**): the Cryptophyceae-Chlamydia group branches close to chloroplast paralog in 1915-intron phylogeny, while in the C-terminus phylogeny, this group is branching with the Diaphoretickes group. Topology searches along GPI's MSA partitions shows that this signal is exclusive to this region, being supported by both evolutionary models (LG/C20+G+F) and amino acid conservation in the MSA (bin_9, **Figure S5E**). Together, this may suggest that the seemingly 'parallel inserted' intron in Cryptophyceae is due to a recombination between the nuclear and the plastid paralog acquired by secondary endosymbiosis, as well as putative gene exchange between Cryptophyceae (or Archaeplastida) and Chlamydia. In contrast, the introns in Opisthokonta at this same position represent a potential parallel insertion. Thus, while we can speculate that GPI was present in LECA, solving the origin of eukaryotic GPI remains highly uncertain due to potential HGT and recombination events.

**6-phosphofuctokinase — PFKA — K00850**

The reconstruction of PFKA shows two distant groups (PFKA_1 and PFKA_2, **Figure S6**) with little taxonomic overlap (exceptions include anaerobic amoebozoa and basal opisthokonts) (**Figure 3A**). PFKA_1 contains a tandem duplication of the PFK Pfam domain and it is mainly found in Amorphea, with few exceptions like Rhodelphis, and Heterolobosea-Tetramitia as well as in Bacteria (**Supp. Figure S6A,C**). PFKA_2 by contrast, mainly comprises eukaryotic homologs and is found in Diaphoretickes and Excavata, including a small and taxonomically mixed bacterial clade (**Figure S6D**). Based on a larger phylogeny that includes related enzyme families, i.e. PFK and PFP, it seems that PFKA_2 and PFKA_1 are monophyletic (**Figure S6B**, left panel). Phylogenetic trees based on alignments that include individual PFKA_1 N- and C-terminal domains, suggest instead that the PFKA_1 is derived from a tandem duplication of PFK domain, while PFKA_2 places at a different place in the tree i.e PFKA_1 and PFKA_2 are not monophyletic (**Figure S6B,** right panel). This may indicate that the initial placement of PFKA_2 with PFKA_1 is a result of LBA or that the alignment as a whole does not contain enough information to resolve the placements accurately. PFKA_2 might have been encoded by LECA but its origin, though likely bacterial, is unclear. In contrast, based on the limited distribution of PFKA_1 in eukaryotes, it seems likely that this family was not encoded by LECA. The congruent topologies of PFKA_1 N- and C- terminus subtrees, in which Amoebozoa are closer to the bacterial clade (*extended search* in **Figure S6C,** inset tree), suggests gene exchange between this eukaryotic lineage with bacteria. In line with this, the exact prokaryotic origin and timing of the tandem domain duplication for the origin of PFKA_1 is somewhat tentative.

**Fructose-biphosphate aldolases — ALDO, FBA**

**Fructose-biphosphate aldolases, Class I — ALDO — K01623**

The fructose-bisphosphate aldolase class I, ALDO, is widespread in eukaryotes, present in Bacteria, and apparently less frequent in Archaea (**Figure S7A,C**). The widespread distribution and the monophyletic branching of eukaryotic sequences suggest that ALDO was present in LECA. The unrooted view of the ALDO reconstruction shows diverse alphaproteobacteria taxa as closest relatives of the eukaryotic ALDO (**Supp. Figure S7A**) indicating that ALDO is of alphaproteobacterial origin. Note that we observed potential indications of HGTs from eukaryotes to bacteria and/or viruses (**Figure S7C**). Archaeaplastida (except Glaucophyta and Rhodelphis) encode two copies of ALDO, but it is unclear whether they represent lineage-specific or ancestral duplications in Archaeplastida.

**Fructose-biphosphate aldolases,  Class II — FBA — K01624**

The phylogeny of the fructose-bisphosphate aldolase class I, FBA, shows one main eukaryotic group composed of Amorphea, Diaphoretickes and Euglenozoa, suggesting its potential ancestral presence in LECA (**Figure S7B,D**). However, please note the scarce representation of this enzyme in Amorphea. The sister group of this potential LECA clade is composed of a larger diversity of bacteria with mixed taxonomy. Therefore, the donor of this eukaryotic FBA group remains unknown but appears to be bacterial. In addition, we found other smaller clades of  eukaryotic homologs including photosynthetic eukaryotes. One comprising Chloroplastida, secondary endosymbiotic eukaryotes and Dikarya Fungi with few Cyanobacteria, and another comprising Glaucophytes nested within Cyanobacteria (**Figure S7D**). While the former eukaryotic group might be derived by HGT from Cyanobacteria, the latter group might be the result of EGT with homologs being retained in Glaucophyta and lost in the rest of Archaeplastida. Finally, there are diverse anaerobic eukaryotes interspersed within diverse bacteria most likely indicating independent later acquisitions (**Figure S7D**).

**Triosephosphate isomerase — TPI — K01803**

The phylogeny of TPI shows a deep separation between archaeal homologs on one side and eukaryotic and bacterial homologs on the other side (**Figure S8A**). Eukaryotes form one main monophyletic group comprising sequences from all eukaryotic clades (**Figure S8B**) suggesting that TPI was present in LECA. Basal to this LECA group, we found another eukaryotic group comprising Chloroplastida and secondary endosymbionts whose phylogenetic locations differ when excluding archaea (**Figure S8A,B**). This shows the phylogenetic instability of this specific group which likely has an independent origin, or alternatively, increased sequence divergence. Nevertheless, the basal location of Alphaproteobacteria (and more closely related Gammaproteobacteria), to the LECA group of TPI (topology obtained when only excluding archeal sequences from the reconstruction, **Figure S8B,C**), suggests that TPI is of alphaproteobacterial origin, as previously suggested (Keeling and Doolittle 1997). Note that Wallbacteria was also found basally to eukaryotes, but with a long basal branch, making this relationship uncertain.

**Glyceraldehyde 3-phosphate dehydrogenase — GAPDH — K00134**

The phylogeny of GAPDH shows four distantly related major eukaryotic groups that comprise diverse bacterial sister groups (**Figure S9**). One clade comprises Archaeplastida and is likely of cyanobacterial origin, though some secondary endosymbionts are also closely related. Another group is formed by a large diversity of eukaryotic homologs with bacterial clades interspersed. Thus, this later group suggests that GAPDH was present in LECA and followed independent HGT from eukaryote-to-bacteria and subsequent HGTs between bacteria, as previously pointed out (Takishita and Inagaki 2009). The sister group to this large eukaryotic clade is composed of Chlamydia and Cyanobacteria (plus other bacteria and Euglenozoa) and therefore the donor lineage is unclear, though most likely bacterial (Martin et al. 1993). Finally, two additional eukaryotic clades appear to be of bacterial origins. While one of these might be derived from alphaproteobacteria, the acquisition might post-date LECA.

**Phosphoglycerate kinase — PGK — K00927**

The phylogeny of PGK seems to be widespread in prokaryotes (**Figure S10A**) with eukaryotic homologs forming three main clusters affiliated with bacterial sequences (**Figure S10B**). The first group is composed of Archaeplastida and secondary endosymbionts, which are branching together with Cyanobacteria and suggesting primary and secondary EGTs. However, basal to this group we also found non-photosynthetic Diaphoretickes, maybe suggesting more complex evolution within this group. The second group is composed by a large diversity of eukaryotes from all the major groups, suggesting that PGK was present in LECA (note that the topology between eukaryotes does not strictly follow the eToL evolution, possibly due to lack of phylogenetic signal as indicated by low branch supports). Planctomycetota sequences are branching basal to this eukaryotic group, suggesting a potential acquisition from this phylum (97% optimized Ufboot2). Finally, the third group suggests an independent acquisition in Discoba-Euglenozoa organisms from bacteria as it branches next to Verrucomicrobiota.

**2,3-bisphosphoglycerate phosphoglycerate mutases**

There are several distinct phosphoglycerate mutase families that are non-homologous isofunctional enzymes with independent evolutionary origins and no similarity in primary sequence, 3D structure, or catalytic mechanism. Cofactor-dependent phosphoglycerate mutases require 2,3-bisphosphoglycerate for activity while cofactor-independent PGM (iPGM) do not (Foster et al. 2010).

**GPMA — K01834 (cofactor-dependent)**

The phylogeny of GPMA shows the two main groups of eukaryotes with poorly defined sister groups due to limited phylogenetic signal (**Figure S11A**). One of these eukaryotic GPMA clades branches with some bacteria and as well as Asgardarchaeota (Lokiarchaeia and Hemindallarchaeia). However, given the low resolution in the basal nodes in GPMA reconstruction, it is unclear whether those eukaryotic homologs are derived from the archaea.

**GPMB — K15634 (cofactor-dependent)**

The phylogeny of GPMB shows its widespread distribution in prokaryotes and in eukaryotes. There is one main eukaryotic clade although other 'minor' eukaryotic clades are found with unclear sister groups (**Figure S11B**). The lack of phylogenetic resolution for defining sister groups hinders the inference about the origin of GPMB in eukaryotes.

**GPMI — K15633 (cofactor-independent)**

The reconstruction of GPMI shows a peculiar evolutionary history regarding the relationship of eukaryotes and Asgardarchaeota. GPMI is widespread in prokaryotes with indications for frequent HGTs, while eukaryotic sequences form two distant groups (**Figure S12A**). Both groups contain taxa from the major eukaryotic clades, but with a patchy distribution (**Figure 3A**). GPMI_1 sequences form a monophyletic group whereas the monophyly of eukaryotic GPMI_2 is less clear. GPMI_2 forms two eukaryotic subclades: one highly supported clade is composed of mainly anaerobic eukaryotes like Metamonada and some Amoebozoa with Heimdalarchaeia sequences as sister group, while the other main subgroup comprises mainly Diaphoretickes, Discoba, CRuMs and Breviatea with diverse bacteria, including Myxoccocota as sister group. In turn, the origin(s) of eukaryotic GPMI_2 remain unclear, since the phylogeny is compatible with various scenarios: one single origin in LECA with subsequent HGT to prokaryotes, two post-LECA acquisitions, or one LECA (with Asgardarchaeota origin) and another post-LECA (bacterial origin) acquisition. Eukaryotic GPMI_1 in contrast, appears to have been encoded by LECA and may have been derived from Heimdalarchaeia, as they are branching in the sister group. Remarkably, GPMI_1 and GPMI_2, do not present taxonomic overlap in eukaryotes (**Figure 3A**), and distinct paralogues are encoded by some closely related eukaryotic taxa including Metamonada and Rhodophytes (no evident HGT of GPMI_1 and GPMI_2 between eukaryotes is found **Figure S12A**). This could potentially indicate that LECA harbored both GPMI_1 and GPMI_2 with subsequent differential loss in the various eukaryotic taxa. Alternatively, LECA had GPMI1 and the presence of GPM2 in eukaryotes might be explained with post-LECA transfers, followed by HGTs between eukaryotes.

**APGM — K15635 (cofactor-independent)**

APGM shares a homologous region to GPMI and both carry out the same reversible enzymatic reaction, from 3-phosphoglycerate to 2-phosphoglycerate. APGM is widespread in prokaryotes, showing large monophyletic groups of Bacteria and Archaea, but also a mix of them indicative of various HGTs (**Figure S12B**). The APGM phylogeny recovered one clade of eukaryotes encompassing Excavates, Diaphoretickes as well as Amorphea including Fungi, CRuMs and Breviatea, suggesting that APGM was present in LECA. Considering that the closest sister group of this eukaryotic clade comprises TACK and Asgardarchaeota, it seems likely that eukaryotic APGM was acquired from the archaeal-host.

**Enolase — K01689**

Enolase is widespread in all three domains of life (**Figure S13A**). Eukaryotic homologs form two major clades (here referred to as enolase 1 and 2), each of which seems to have been present in LECA and is phylogenetically related to archaeal sequences as previously shown (Hannaert et al. 2000). One of these clades comprises a large diversity of eukaryotic homologs from Excavates, Diaphoretickes and Amorphea (**Figure S13B**) and clusters with a clade of Hemindalarchaeia and DPANN sequences, though, with low support (39% optimized UfBoot2). It is unclear whether the placement of those DPANN homologs is due to LBA artifacts or indicates a history of gene transfer among Archaea. Nonetheless, given that eukaryotes are closely related to archaeal sequences, it is likely that this Enolase family is derived from the archaeal ancestor of eukaryotes. The other eukaryotic enolase clade (Enolase 2) comprises sequences from major eukaryotic groups, although these appear to have long terminal branches (note that some sequences of this group were removed in filtering processes, **Figure S13C**) and it is unclear whether this clade corresponds to a LECA group. The closest archaeal sister group is composed of Methanomassiliicoccales (89% optimized UfBoot2), which among others have been found associated with animals. In turn, it remains to be elucidated whether this placement reflects a more recent transfer to Eukaryotes, or is indicative of phylogenetic artifacts.

**Pyruvate kinase — PK — K00873**

The phylogeny of PK indicates a complex history of gene transfers across the domains (**Figure S14A**). Eukaryotic sequences form six distincts phylogenetic groups (PK_1-6) probably suggesting multiple origins. PK_1 has a sparse composition of diverse eukaryotes from the three major groups, and groups with DPANN, euryarchaeal and bacterial sequences, though these placements are not supported (**Figure S14B**). PK_2 represents a clear ancestral duplication in Archaeplastida with potential origin in Patescibacteria (**Figure S14B**). PK_3 is restricted to diverse Diaphoretickes and Amorphea taxa, with a taxonomically diverse (mostly bacterial) sister group (**Figure S14C**). PK_4 is composed of a mix of secondary endosymbionts from Cryptista and SAR, but also contains a paraphyletic group consisting of Metamonada-Fornicata, which are branching within Alphaproteobacteria (**Figure S14C**). Due to the paraphyletic branching and limited taxonomic composition, these two groups suggest post-LECA independent HGT acquisitions from Alphaproteobacteria. PK_5 is composed of a few distant eukaryotes including Amoebozoa, Discoba-Heterolobosea, and Haptista-Centroplasthelida, whose sequences are branching within a large group of Asgardarchaeota (**Figure S14D**). One possible interpretation is that PK_5 was derived from the asgardarchaeal ancestor followed by massive gene loss across eukaryotic evolution. In this case, eukaryotic PK_5 might be considered as 'jotnarlog' (More et al. 2020). The archaeal-host origin of PK would be in line with the common evolutionary history of the two previous enzymatic steps, ENO and APGM, both of archaeal origin (*see above*). Alternatively, PK_5 might also suggest a post-LECA transfer from Asgardarchaeota probably to Amoebozoa, with subsequent HGT between eukaryotes. PK_6 is the largest eukaryotic group comprising seq uences from the three major groups suggesting ancestral presence in LECA, showing well conserved duplication in Chloroplastida (**Figure S14E**). The closest and well supported (100% optimized UfBoot2) sister clade is mainly composed of members of Fusobacteria indicating the potential donor to this eukaryotic orthogroup. Altogether, the phylogeny of PK enzymes suggests the possibility of one or more paralogues in LECA that were derived from distinct prokaryotic sources including the Alphaproteobacteria, Asgardarchaeota and potentially Fusobacteriota.

**Evolution of pentose-phosphate pathway (PPP) enzymes**

**Glucose 6-phosphate dehydrogenases**

**Glucose-6-phosphate 1-dehydrogenase — G6PD, zwf — K00036**

The first step in the PPP as well as in the Etner-Doudoroff glycolysis is the conversion of glucose-6P into gluconate-6P, through the combination of two enzymes G6PD and 6-phosphogluconolactonase, PGLS, which in some cases are fused (**Figure S15**). G6PD is widespread in Bacteria and in Eukaryotes, but not in Archaea (except for some DPANN, **Figure S15A**). The phylogeny of G6PD revealed one major clade comprising a large diversity of eukaryotic homologs (**Figure S15B**), suggesting that G6PD was present in LECA. Furthermore, the phylogeny indicated a bacterial origin of this LECA protein family which branches sister to a bacterial clade consisting predominantly of homologs of Verrucomicrobiota, Planctomycetota and Hydrogenedentota, suggesting potential origin of PVC-related clade.

**Hexose-6-phosphate dehydrogenase — H6PD — K13937**

The eukaryotic G6PD family comprised a clade of long branching H6PD homologues. While the phylogenetic placement of H6PD within the G6PD group is unclear, this suggests that H6PD might be an eukaryotic invention. Several genes encoding H6PD homologues (but also some G6PD sequences) are fused to a gene for PGLS characterised by a glucosamine_iso Pfam domain (PF01182) (**Figure S15B**). This gene fusion allows the direct conversion of glucose-6P into gluconate-6P and seems to have occurred independently in different eukaryotic lineages such as in Chlorarachnea (Rhizaria) and Myzozoa (Alveolates) characterized by the gene fusion at the C- and N-terminus, respectively. H6PD might have originated in Holozoa, with subsequent transfers to diverse protists from Euglenida or SAR.

**6-phosphogluconolactonase — PGLS — K01057**

The PGLS appears to be widespread in Eukaryotes (**Figure S15C**). Eukaryotic homologs form four major clades, suggesting diverse acquisitions from prokaryotes (in addition to other smaller groups; **Figure S15D**). Each of these clades contains independent domain fusions with G6PD and other domains. The first group (fusion_1, **Figure S15D**) represents the eukaryotic H6PD, which seems to be the result of the domain fusion of the eukaryotic G6PD with a PGLS most likely acquired from Planctomycetota (100% optimized UfBoot2). A second eukaryotic clade (fusion_2), is composed of a small number of eukaryotic homologs from diverse lineages but with a notorious representation of Excavata taxa. It is uncertain whether this clade corresponds to a LECA group or was acquired later. Notably, this clade places sister to a group of sequences including homologues from two Heimdallarchaeia, though with low support (**Figure S15D**). The third main clade (fusion_3), comprises most eukaryotic diversity with mixed topology and likely represents a LECA clade, acquired from an unknown bacterial donor. The fourth clade (fusion_4) comprises few Metamonada homologs that branch distantly in both PGLS and G6PD phylogenies, suggesting independent bacterial donations.

**NAD+ dependent glucose-6-phosphate dehydrogenase – Azf – K19234**

Azf is a non-homologous epimerase alternative to the G6PD (and H6PD), which was originally described in Archaea (Verhees et al. 2003). This enzyme seems to be relatively widespread in Archaea, but also in some bacteria forming various paraphyletic groups (**Supp. Figure S16**). Azf is mainly found in Diaphoretickes protists, and given the current phylogeny, the prokaryotic origin of this gene in eukaryotes is unclear.

**6-phosphogluconolactonase — PGL — K07404**

The phylogeny of PGL shows a complex evolutionary scenario in which eukaryotic sequences form one main clade but with diverse prokaryotic groups interspersed (**Figure S17**). The low phylogenetic signal combined with multiple HGTs, hinders the inference of the origin and evolution of PGL in eukaryotes.

**6-phosphogluconate dehydrogenase — PGD — K00033**

The phylogeny of PGD shows two main groups: one only composed by prokaryotes which lack a C-terminus region in the MSA and was used to root the tree, and the other composed by diverse intermixed eukaryotic and prokaryotic homologues suggesting a complex evolutionary history (**Figure S18A**). Broadly, eukaryotic sequences form four main clades, each of them composed of distantly related eukaryotes. The largest eukaryotic group comprises Diaphoretickes as well as Discoba, and Amorphea protists like CRuMs, Filasterea, and Ancyromonadida, without presenting clear evidence of HGT between these. Two other main groups of eukaryotic homologues cluster closely together but are separated by diverse bacteria branching between them, among which, PVC bacteria have basal positions. Both eukaryotic groups are mainly composed of Amorphea, although Fungi and Holozoa are monophyletic together with a paralogous group of Rhodophytes (among others), while Amoebozoa are branching in the other group with Cryptista. The fourth group is subtended by a long branch and composed by a few Diaphoretickes protists and Discoba-Metakinetoplastina, most likely suggesting independent origins and post-LECA HGT between eukaryotes or lack of phylogenetic signal between eukaryotic sequences. Additionally, we also found discrete paraphyletic groups of Metamonada and Fungi like Microsporidia and Dikarya forming long branches along this part of the tree. Given the low resolution in most of basal nodes, and the disparate compositions of each eukaryotic group (**Figure S18B**), it is unclear whether any or all of these paraphyletic eukaryotic clades actually represent a LECA group with subsequent gene transfers to prokaryotes or whether they represent distinct bacterial acquisitions with post-LECA HGT from bacteria, followed by HGT between eukaryotes. The conservation of some introns along the PGD eukaryotic family, are supportive of LECA origin, with subsequent transfer to prokaryotes, (i.e introns at positions 0988, 1488, 1650 and more tentatively 0960/0962) (**Figure S18C**). If anything, the phylogeny suggests significant gene exchange between bacteria and eukaryotes.

**Ribulose-phosphate 3-epimerase — RPE — K01783**

The phylogeny of RPE reveals three major clades of eukaryotes (**Figure S19A**). The largest clade comprises homologs of all eukaryotic taxa and thus likely represents a LECA clade (**Figure S19B**). While this eukaryotic LECA clade includes bacterial sequences (**Figure S19B**), this might be a phylogenetic artifacts in the phylogeny of the whole family because the bacterial homologs branch outside of this LECA clade when the phylogeny is refined (**Figure S19C**). Basal to this LECA group, we find bacterial sequences of diverse taxonomic compositions, suggesting that RPE was a bacterial contribution during eukaryogenesis and not directly derived from the alphaproteobacterial or asgardarchaeal ancestor. Our analysis revealed a Planctomycetota clade positioned basally to the LECA clade. This clade exhibited a relatively stable affiliation with the eukaryotic clade, albeit with low bootstrap support in the refined phylogeny (49% Ufboot2, **Figure S19BC**), suggesting a possible HGT acquisition of RPE from Planctomycetota. A distantly placed eukaryotic clade was composed of primary and secondary endosymbionts and emerged from within Cyanobacteria in phylogenetic reconstructions using the best empirical fitting model, i.e. LG+R10 (**Figure S19D**). However, the topology differed when using a complex model (C20+G+F), in that Cyanobacteria seemed to emerge from within eukaryotic homologs (**Figure S19B**). Yet, based on the phylogeny obtained using the empirical model, it may be hypothesized that RPE was acquired from Cyanobacteria through primary endosymbiosis, followed by secondary endosymbiosis. Finally, we identified a third eukaryotic clade composed of secondary endosymbiotic eukaryotes from SAR, Haptista and Cryptista with Gammaproteobacteria as a potential donor (**Figure S19B**).

**Ribose 5-phosphate isomerase A — RPIA — K01807**

RPIA seems to be widespread in Archaea with the phylogeny showing evidence for various interdomain transfers between Archaea and Bacteria (**Figure S20A**). Eukaryotic homologs form two main distinctive clades here referred to as RpiA_1 and RpiA_2, and a small group composed of sequences from unicellular Streptophyta branching with Cyanobacteria (**Figure S20A**). The eukaryotic RpiA_1 homologs, prevalent in Amorphea, might represent a LECA clade as it branches sister to homologs of Asgardarchaeota and TACK (**Figure S20B**). A clade consisting of lokiarchaeal homologues is placed within eukaryotic RpiA_1, which could be due to gene tree reconstruction artifacts or to HGTs between pre-LECA organisms and Asgardarchaeota (Wu et al. 2022). Eukaryotic RpiA_2 homologs, by contrast, branch distantly to RpiA_1, and cluster with a diverse set of bacterial homologs of various taxonomic affiliation (**Figure S20C**). Note that RpiA_2 includes a diversity of homologs of secondary endosymbionts that form a monophyletic group and are likely derived from secondary EGTs. It is unclear whether eukaryotic RpiA_2 homologs, which seem to be predominantly present in Archaeplastida and secondary endosymbiont, represent a LECA clade. Evolutionary analysis of RpiA suggests its presence in LECA, potentially acquired from the archaeal host. Subsequently, RpiA underwent multiple gene loss events in eukaryotic lineages belonging to the Diaphoretickes and Excavata supergroups. These losses were likely followed by functional replacements with paralogs (RPIA) and analogs (RPIB, *see below*) derived from bacterial contributions.

**Ribose 5-phosphate isomerase B — RPIB — K01807**

RpiB seemed to be more widespread in Bacteria than in Archaea (**Figure S21**). The phylogeny shows three distant and discrete groups of eukaryotic homologs, each with limited support. The biggest group encompasses all the diversity of eukaryotes, including various sequences from anaerobic eukaryotes, which together form a sister group of Spirochaetota, though with low support (69% Ufboot2; **Figure S21B**). Note that most of these eukaryotes do not have the isofunctional enzyme RpiA. Another group of eukaryotes includes homologs of Streptophyta, secondary endosymbiotic eukaryotes and Fungi-Mucoromycota. One cyanobacterial sequence branches basal to this group, but given the sparse distribution of this enzyme in extant members of this lineage, an EGT origin is unlikely. Finally, a third group comprises sequences of some Excavata and Ciliophora as well as Heimdalarchaeaia and Alphaproteobacteria. However, the low support for the clustering of these sequences and the scarce distribution in eukaryotes prevents drawing clear conclusions about their origins.

**Transketalose — TKTA/B — K00615**

The transketolase is involved in producing erythose-4P and heptolulose-7P from different substrates like fructose-6P, ribose-5P, or xylulose-5P, reversibly.

The transketolase revealed several clades of eukaryotic groups (**Figure S22A**), the two largest of which we refer to as Group_1 and Group_2 (**Figure S22C**). Group_1 includes sequences of photosynthetic eukaryotes (Archaeplastida and secondary endosymbionts) and Cyanobacteria, but also one Rhodelphis sequence branching basal to this group. In contrast, Group_2 includes homologs from a wide diversity of eukaryotes suggesting that TKT was present in LECA, but was lost in photosynthetic Archaeplastida. Both LECA and chloroplast clades are closely related and stable in our reconstructions. Thus, it is unclear whether both groups originated from Cyanobacteria or whether they have distinct origins. Basal to these two eukaryotic groups, we found a large diversity of bacteria, including  a group of Alpha/gammaproteoabacteria. However, due to the distant placement of those alphaproteobacterial sequences, the origin of eukaryotic TKT remains unclear, although it appears to be bacterial.

In addition to these, we also found minor clades mainly comprising secondary endosymbiotic eukaryotes, branching sister to Chlamydia and Verrucomicrobiota sequences (root group in **Figure S22C**) and another distant clade comprising Metazoa, Alveolates and deep branching Cryptista, grouping with diverse bacterial sequences (asterisk in **Figure S22AC**).

**Transaldolase — TALA/B — K00616**

Erythrose-4P and heptolulose-7P, are then converted into metabolites of glycolysis like Glyceraldehyde-3P and fructose-6P through the transaldolase.

The phylogeny of transaldolase shows three major distinct phylogenetic groups, one of which contains a large diversity of eukaryotes as well as some bacterial sequences (**Figure S22B**). We selected sequences of this group to perform a refined phylogeny resulting in a single monophyletic group of eukaryotes (**Figure S22D**), most likely suggesting that TAL was present in LECA. The sister group of this clade comprises diverse bacteria mainly including an ancestral clade of Chlamydiota and Desulfobacterota, few Gammaproteobacteria, but the support is moderate (73% optimized UfBoot2) and does not allow to draw conclusions regarding origins of the TAL LECA family, although it appears to be bacterial.

**Entner-Doudoroff Glycolysis**

The Entner-Doudoroff (ED) glycolysis is an alternative glycolytic pathway that overlaps with the PPP, since both pathways share enzymes for producing gluconate-6P. The gluconate-6P is used to produce pyruvate and G3P through the consecutive reactions made by EDA and EDD. The ED pathway has been characterized in Archaeplastida, and has been suggested to be of cyanobacterial origin (Chen et al. 2016). Both EDA and EDD enzymes occur mainly in Archaeplastida and other protists, although EDD is also found in Amorphea taxa, especially in Fungi (Chen et al. 2016). While our results for the evolution of EDA might support the assumption of cyanobacterial origins, the evolution of EDD suggests a more complex evolutionary history in terms of donor(s) and evolution within eukaryotes (**Figure S23**; see below).

**Phosphogluconate dehydratase, dihydroxy-acid dehydratase —  EDD, ILVD — K01690, K01687**

The respective KO of EDD (K01690) is found in few prokaryotes, but its closest KO is the dihydroxyi-acid dehydratase, ILVD (K01687) and has been described to have the same function as EDD in ED glycolysis. The phylogeny of the combination of both KOs shows EDD/ILVD is widespread in prokaryotes and in some eukaryotes (**Figure S23AC**). Eukaryotes form two clades, one of which contain Chloroplastida and the other Rhodophyta, each of which comprises members from Amorphea and some Discoba (**Figure S23C**). Between these two groups of Eukaryotes, we found diverse bacteria among which Cyanobacteria seems to form a monophyletic clade. Based on our analysis, this gene family seems to be more widespread in Eukaryotes than originally assumed and indicates a complex history of evolution with additional bacterial (though not cyanobacterial) donations.

**2-dehydro-3-deoxyphosphogluconate aldolase — EDA — K01625**

The phylogeny of EDA, widespread in prokaryotes, suggests extensive HGTs (**Figure S23BD**). Eukaryotes form three discrete groups, most likely representing different acquisitions from bacteria. One of these clades comprises a monophyletic group of Chloroplastida and Rhodophyta, and secondary endosymbionts, but also Telonemia. This clade is sister to a cyanobacterial clade suggesting an origin of EDA through the plastid ancestor. The other two groups are mainly composed of secondary endosymbionts and Cryptista among others, respectively, with unknown bacterial donors. Remarkably, most of the eukaryotes in these paraphyletic groups (except for Fungi **Figure S23C**), also encode for EDD, illustrating the phylogenetic incongruences between the history of both enzymes in eukaryotes.

**Ribose-phosphate pyrophosphokinase — PRSA — K00948**

PRSA uses ribose-5P and ATP to produce 5-Phosphoribosyl diphosphate (PRPP), which is the precursor for nucleotide and histidine biosynthesis among others. Acknowledging the low phylogenetic signal of this protein family (i.e. limited amino acid conservation and short MSA, ~300 positions), the phylogeny of PRSA separates Bacteria and Archaea domain with some exceptions (**Figure S24A**). Eukaryotic sequences form four main clades: one of them with unclear origin and paraphyletic with bacterial groups (PRSA_1), while the other groups (PRSA_2-4) seem to have bacterial sister-groups (**Figure S24**). Especifically, eukaryotic PRSA_1, which also comprises bacterial homologs, is a divergent group branching between archaeal sequences (**Figure S24AB**). PRSA_2 is found in most eukaryotic taxa including Amorphea, Diaphoretickes and Excavata (**Figure S24C**). PRSA_2 represents a LECA clade that is phylogenetically associated with proteobacterial sequences, suggesting a mitochondrial origin. Note that this topology was only recovered when using empirical models (LG+R10). PRSA_3 is exclusively composed of Archaeplastida and is phylogenetically associated with cyanobacterial sequences, suggesting a chloroplast origin (**Figure S24D**). PRSA_4 is a minor clade that encompasses all eukaryotic diversity and has a bacterial sister group, but with no support to identify the donor (**Figure S24E**).

The topology of eukaryotic PRSA_2 is broadly in agreement with the species tree, illustrating a vertical evolution from LECA (with some post-LECA duplications in Amorphea, and gene losses in Archaeplastida **Figure 3A and S24C**). In contrast, the complex evolution of PRSA_1, which might be derived from the archaeal host, can potentially be explained by it being redundant with PRSA_2. This genetic redundancy might have selection pressure for one of the PRSA copies, including loss and gene exchange with bacteria. The long basal branch of PRSA_1 might be seen as in agreement with studies (Pittis and Gabaldón 2016) suggesting that genes derived from the archaeal host have longer relative branch lengths' than those from the bacterial partner. However, alternative explanations like accelerated evolution cannot be excluded.

**Evolution of pyruvate, acetate and citrate conversions into acetyl-CoA**

**Reversible conversion of pyruvate to acetyl-CoA**

**Pyruvate dehydrogenase complex — PdhABCD(X) — K00161, K00162, K00627, K00382, K13997**

Pyruvate dehydrogenase (PDH) complex is composed of three to four subunits: PdhA/B are pyruvate dehydrogenase subunits (E1), PdhC is a dihydrolipoamide acetyltransferase subunit (E2), and PdhD (and PdhX) is a dihydrolipoamide dehydrogenase subunit (E3). The consecutive reaction of these enzymes converts pyruvate into acetyl-CoA through pyruvate decarboxylation. PdhABCD phylogenies are congruent regarding the placement of eukaryotic LECA groups clustering with alphaproteobacterial homologs (**Figure S25**). PdhC/X are homologous, and their phylogeny suggests that PdhX originated as a result of mitochondrial PdhC duplication during eukaryogenesis (**Figure S25C**).

Eukaryotic sequences form two main congruent phylogenetic groups in the respective PdhABCD phylogenies: one LECA group clustering with alphaproteobacterial sequences, and another group of sequences of photosynthetic eukaryotes clustering with cyanobacterial sequences (**Figure S25**). The alphaprotebacterial PDH complex is found in nearly all eukaryotes except for anaerobic Metamonada and Dinoflagellata secondary endosymbionts, that only encode for PdhD homologs. While Metamonada's PdhD are forming paraphyletic groups that branch within alphaproteobacteria (**Figure S25D**), these sequences did not pass the Chi test (**Figure S25D,** lower panel), suggesting that these sequences may be compositionally biased. A downsampled phylogeny of this mitochondrial group shows that some of these divergent eukaryotic sequences are indeed branching within the PdhD LECA (**Figure S25D**, lower panel, bold labels) supporting the mitochondrial origin of PdhD subunits in Metamonada. On the other hand, the chloroplast PDH complex shows a patchy distribution in photosynthetic eukaryotes (**Figure S25**) and might have been acquired through at least two independent secondary endosymbiosis event: one from Rhodophyta (including Haptista, Cryptista and Stramenopile and Alveolates secondary endosymbionts), and another from Chlorophyta (in Rhizaria-Cercozoa-Chlorarachnea; **Figure S25CD**). Remarkably, Glaucophyta organisms are not found in this chloroplast group. Thus, and as previously pointed out (Schnarrenberger and Martin 2002), the PDH complex was acquired from alphaproteobacteria, and also from the cyanobacterial ancestor of plastids.

**Pyruvate-ferredoxin/flavodoxin oxidoreductase — POR/Nifj — K03737**

Pyruvate-ferredoxin/flavodoxin oxidoreductase (PFOR) is an alternative route for the reversible conversion of pyruvate into acetyl-CoA that reduces ferredoxin, unlike the PDH complex that reduces NAD+. PFOR is often associated with anaerobic/microaerophilic metabolisms in eukaryotes (Stairs et al. 2015) although it also functions in aerobic organisms like Cyanobacteria (Wang et al. 2022). The phylogeny of PFOR recovers eukaryotic homologs as a single monophyletic group that branches with homologs of Thermoplastota-E2, Chlamydia-Anoxychlamydiales (**Figure S26**). A sister relationship of Anoxychlamydiales with anaerobic eukaryotes, has been previously observed in this and other anaerobic-related genes in eukaryotes (Stairs et al. 2020), suggesting that certain genes for the anaerobic metabolism of eukaryotes might have been acquired from this Chlamydia order. Notably, Thermoplasmatota-E2 (Thermoprofundales order; (Sheridan et al. 2022)), are also found sister to eukaryotic groups in AclA/B/Y phylogeny (*see below*), suggesting an alternative/additional putative donor besides the Chlamydiota. Nevertheless, it is unclear whether PFOR was present in LECA and followed multiple losses across eukaryotic evolution, or whether it represents a post-LECA acquisition that was later transferred between eukaryotes, as previously discussed (Stairs et al. 2020).

**Formate C-acetyltransferase — PflD — K00656**

The formate C-acetyltransferase catalyzes the reversible formation of acetyl-CoA and formate from pyruvate and CoA. The phylogeny of PflD defines two distant phylogenetic groups with limited taxonomic distribution (**Figure S27A**). One of them includes a monophyletic group of eukaryotes comprising diverse anaerobic eukaryotes as well as secondary endosymbionts together with Archaeplastida (**Figure S27B**). The scarce and mixed phylogenetic relationship within this eukaryotic clade suggests a post-LECA acquisition, as previously pointed out (Stairs et al. 2011). However, in contrast with previous analyses that inferred a Firmicute origin for PflD in eukaryotes (Stairs et al. 2011), our phylogeny did not resolve a clear sister group or donor lineage, although Firmicutes-Clostridia (but also others) were closely related (**Figure S27B**).

**Reversible conversion of acetate to acetyl-CoA**

**Acetyl-CoA synthetase — ACS – K01895**

Acetyl-CoA synthetase (ACS) catalyzes the reversible formation of acetate and ATP from acetyl-CoA by using ADP and phosphate, although it can also synthesize acetyl-CoA from short-chain fatty acids. Eukaryotic sequences form two main clades that are closely related , but with different bacterial groups interspersed (**Figure S28**). Each eukaryotic clade comprises most eukaryotic lineages, indicating that both paralogous groups may have been present in LECA (note that one group does not contain Archaeplastida, **Figure S28B**). One of these eukaryotic clades, ACS1, branches to diverse bacteria including Planctomycetota, Verrucomicrobiota, Myxococcota and Spirochaetota, among others. Similarly, the other LECA clade forms a sister group with homologs of amongst others Chloroflexota, Planctomycetota and Verrucomicrobiota, Myxococcota and Desulfobacterota. The sister group relationships of each ACS subfamily are well supported (>99% optimized UfBoot2; **Figure S28B**). However, it is important to note that an ancestral group of Proteobacteria, including Alpha/Gammaproteobacteria and Magnetococcia, is branching intermediary between both LECA clades ACS1 and ACS2. In addition, despite the paraphyletic branching of ACS1 and ACS2, both subfamilies share intron positions (**Figure S28C**). However, given the scarcity of these and their unbalanced conservation, it is unclear whether they suggest a single origin of ACS1 and ACS2 followed by a subsequent duplication or distinct origins with parallel insertions of introns. Therefore, although LECA appears to have encoded two ACS paralogs, inferences regarding their evolutionary origins remains speculative.

**Acetate---CoA ligase (ADP-forming) subunit alpha — AcdA,B,AB – K01905, 22224, K01905**

AcdAB catalyzes the reversible formation of acetate and ATP from acetyl-CoA using ADP and phosphate. AcdA and AcdB contain the CoA binding domains and ATPase domain, respectively. The ATPase domain (ATP-grasp_5) is homologous to the ATPase domain of ATP-Citrate lyase (AclA, ATP-grasp_2) involved in the reverse TCA. AcdAB has been shown to be relevant in anaerobic eukaryotes like Metamonada (Williams et al. 2023). Due to the complex evolution of the AcdA/B, we performed a phylogeny of each subunit, AcdA (including only ATP-grasp domains) and AcdB (including CoA bindlind/ligase and Suc_CoA_ligase domains; **Figure S29**). While AcdB (ATP-grasp_5) was rooted at the AclB subfamily (ATP-grasp_2; **Figure S29A**), the deepest branching AcdB sequences were used to root the AcdA phylogeny (**Figure S29B**). Eukaryotic AcdA/B homologues show diverse architectures, i.e. fused or split, additional domains like Acetyltransferase (at the N- or C-terminus), and rearrangements of domains in fused proteins, i.e. AcdA at the C-terminus and AcdB at the N-terminus (Acd_1 group; **Figure S29**). The phylogenies of AcdA/B seem to be widespread in Archaea while the distribution in Bacteria appears more scattered (**Figure S29AB**). We found five groups mainly comprising anaerobic eukaryotes, whose phylogenetic relationship with their closest prokaryotic sequences were congruent in both phylogenies (**Figure S29CD**). Acd_1 is the most divergent group in the AcdB phylogeny and is characterized by a rearrangement of N- and C-terminal domains. Acd_1 is closely related to bacterial CPR sequences and might be ancestral in Excavata. While it is also found in Breviatea, this group most likely suggests a post-LECA acquisition. Acd_2 is found in diverse eukaryotes, including Glaucophyta, Myzozoa, Metamonada, and Breviatea, being closely related to Chlamydia sequences, among others. Note that the signal for the definition of the

Acd_2 group regarding prokaryotic sequences is unclear due to prokaryotic sequences branching within eukaryotic ones. Acd_3 is found in Metamonada-Fornicata, and these are closely related to Lokiarchaeia sequences with full support (100%). Acd_4 is found in Amoebozoa-Mastigamoebida branching with diverse prokaryotes. Finally, Acd_5 is found in Metamonada, *Barthelona* sp. and *Monocercomonoides exilis:* while being fused (AcdAB), it is closely related to the split proteins (AcdA and AcdB) of Heimdallarchaeia (**Figure S29CD**). The placement of Acd_3 and Acd_5 (**Figure S29CD**) raises the question as to whether these cases represents phylogenetic artifacts, HGTs between Asgardarcahea and Metamoanda (supported by the fact that both inhabit anoxygenic environments), or represent ancestral LECA proteins that were only retained in these Metamoanda clades, i.e. jnorlogs.

**Reversible conversion of citrate to acetyl-CoA**

**ATP-citrate lyase alpha/beta-subunit — AclA,B,Y – K15230, K15231, K01648**

ATP-citrate lyase catalyzes the cleavage of citrate into oxaloacetate and acetyl-CoA using ATP, having an important role in the biosynthesis of lipid precursors and histone acetylation among other cellular proceses. Similar to AcdAB, ATP citrate lyase is composed of two subunits, AclA and AclB, which can be fused in eukaryotes, in which case it is referred to as AclY. We performed phylogenetic analyses of AclA, AclB (including the respective AclY regions), and the concatenation of both (**Figure S30**). The tree was rooted with a divergent group of bacteria including Nitrospira and Chlorobi sequences, in agreement with previous inferences based on the rooted ATP-grasp domain phylogeny (ACDB, **Figure S29A**). AclA and AclB are found in various archaea including Asgardarchaeota as well as some bacteria. Eukaryotic sequences form two clades, i.e. AclA/B and AclY (**Figure S30**). The mixed taxonomic compositions and the phylogenetic relationship in the respective eukaryotic clades suggests that both forms (separated and fused) were present in LECA. Notably, a deep-branching clade of Asgardardarchaeota forms a sister group to these eukaryotic families intermixed with diverse other bacteria and archaea. While the split AclA/B clades are closely related to DPANN archaea, eukaryotic AclY has a sister group comprising Thermoplasmatota-E2 archaea (i.e. *Thermoprofundales*) and Desulfobacterota. These sister relationships of DPANN and Thermoplasmatota-E2/Desulfobacterota with their respective eukaryotic clades, were recovered in individual subunits and concatenated reconstructions (**Figure S30**). However, the rooted phylogeny of the single ATP-grasp_2 domain (encoded in AclA subunit) shows that both eukaryotic groups (split and fused) might be more closely related (**Figure S29A**) than observed in the no-outgroup phylogenies (**Figure S30**). In addition, in individual subunit phylogenies, a divergent group of AclY comprising Haptista-Centroplasthelida sequences branch basal to AclY clade (**Figure S30AB**) while in the concatenated phylogeny, this clade branches within the AclA/B clade (**Figure S30C**). We also found AclA/B subunits in Mimiviridae-Tupanvirus viruses, which might have exchanged genes with eukaryotes. The Haptista-Centroplasthelidad class branches with AclY when virus sequences are removed from the concatenated reconstruction (data not shown).

In turn, while AclA/B as well as AclY have been suggested to have arisen early in eukaryotic evolution (Gawryluk et al. 2015), it is unclear whether these two LECA families trace back to a single origin from Asgardarchaeota (a scenario which would indicate a complex history including HGT with other Bacteria and Archaea and/or phylogenetic artifacts) or whether the two main eukaryotic clades were acquired on two occasions from other archaea or bacteria (such as DPANN, Thermplasmatota and Desulfobacterota).

**Evolution of Tricarboxylic acid cycle enzymes**

**Citrate synthase — CS — K01647**

The phylogeny of citrate synthase (CS) shows three distant eukaryotic groups each emerging from within bacterial sequences (**Figure S31A**). CS_1 is widely distributed in eukaryotes suggesting that it was present in LECA (**Figure S31B**). Eukaryotic CS_1 emerges on a long branch with Acidobacteria and Bacteroidetes forming sister groups. CS_2 is also widespread in eukaryotes and the topology might suggest that it was also present in LECA (**Figure S31C**).The eukaryotic CS_2 group forms a sister group to Chloroflexota and some Gammaproteobacteria sequences (100% optimized UfBoot2). CS_3 mainly comprises Diaphoretickes taxa plus Euglenida and some Amorphea like CRuMs and Apusomonadida (**Figure S31D**). CS_3 is monophyletic with Alphaproteobacteria. Given the scarcity of Amorphea and other Excavata taxa, it is unclear whether CS_3 is of mitochondrial origin, or was a post-LECA acquisition by HGT from Alphaproteobacteria. Thus, LECA might have retained up to three paralogs of CS from different bacterial acquisitions, Bacteroidota, Chloroflexota, and Alphaproteobacteria.

**Aconitate hydratase A — AcnA-ACO, ACO2 — K01681, K17450**

Aconitate hydratase A1 (ACO1 or AcnA) is widespread in prokaryotes, including Alphaproteobacteria and Asgardarchaeota among others (**Figure S32AB**). Eukaryotic sequences form a single monophyletic group comprising all eukaryotic diversity and seemingly branch sister to a group consisting of among others TACK, Thermoplasmatota, Euryarchaeota, DPANN and Chlamydiota, which makes unclear the identity of the donor. In addition, the phylogeny would suggest that putative ACO homologs of alphaproteobacterial and asgardarchaeal partners (if present in those lineages at that time), must have been lost prior to LECA

ACO2 is distantly related to ACO1, and is less widespread in prokaryotes (**Figure S32AC**). ACO2 is found in eukaryotes and forms a monophyletic group encompassing all eukaryotic diversity similar to ACO1, suggesting that ACO2 was also present in LECA. Some taxa like Fungi only retained ACO2, while others like Metazoa retained both. The sister group is well supported (98% optimized UfBoot2) and is composed of diverse bacteria including Bacteroidota and Hydrogenedentota among others.

**Aconitate hydratase B — AcnB — K01682**

Aconitate hydratase B (AcnB) is distantly related to AcnB, and has a limited distribution in prokaryotes (**Figure S32D**). Eukaryotic sequences form a monophyletic group, with CRuMs and Breviatea branching basal to this group. Sequences of this eukaryotic group branch sister to gammaproteobacterial sequences (99% optimized UfBoot2), suggesting this eukaryotic AcnB might have been acquired from Gammaproteobacteria though the timing of the origin is unclear.

**Isocitrate dehydrogenase 1,2,3 — Idh1|2|3 — K00030, K00031**

The isocitrate dehydrogenase family encompasses distant paralogs with distinct evolutionary histories (IDH1, IDH1.1 IDH2, IDH3 and IDH1.3, **Figure S33A**). IDH1 is widespread in eukaryotes and was likely present in LECA. The eukaryotic IDH1 group comprises alphaproteobacterial homologs as well as a sequence from Heimdallarchaeum LC3 (now Hodarchaeaum) and other archaeal and bacterial sequences (**Figure S33B**). While the phylogenetic signal appears to be limited, the widespread distribution of this gene in alphaproteobacteria, suggests them as potential donors. IDH1 is related to a distant group, IDH1.1, which mainly comprises Diaphoretickes (Chlorophyta and secondary endosymbionts, **Figure S33C**). IDH2 is a divergent isocitrate dehydrogenase that is exclusively found in secondary endosymbionts and few Metamoanda, and is closely related to bacterial sequences from Planctomycetota and Bdellovibrionota (**Figure S33D**). Eukaryotic IDH3 homologs form a monophyletic group and were likely present in LECA (**Figure S33E**). IDH3 duplicated during eukaryogenesis giving rise to the alpha and gamma subunit of IDH3 complex, which however, is absent in secondary endosymbionts. The closest sister group of this IDH3 LECA clade is composed of Chlorobi sequences (91% Ufboot2) pointing to them as a potential donor for the eukaryotic IDH3 subunits (**Figure S33AE**).

**2-oxoglutarate dehydrogenase complex— SucA, SucB — K00164, K00654**

The 2-oxoglutarate dehydrogenase complex is composed of three subunits, SucA and SucB and PdhD (the latter of which is described in *Pyruvate metabolism* ssection), and catalyzes the irreversible reaction of 2-oxoglutarate to succinyl-CoA, generating NADH and CO2. Eukaryotic SucA and SucB homologs form monophyletic groups comprising Excavata, Amorphea and Diaphoretickes taxa, which each cluster with alphaproteobacterial homologs (**Figure S34AB**). Thus, 2-oxoglutarate dehydrogenase complex (SucA/B) was present in LECA and it is of alphaproteobacterial origin.

**Succinyl-CoA synthetase complex — LSC1, LSC2 — K01899-K01902, K0900-K01903**

Succinyl-CoA synthetase complex is composed of two subunits, LSC1 and LSC2, which are widely distributed in prokaryotes and eukaryotes (**Figure S35AB**). Eukaryotic sequences form a single monophyletic group in LSC1 and LSC2 phylogenies, encompassing all eukaryotic diversity, including Metamonada (**Figure S35CD**). Alphaproteobacterial sequences are closely related to these eukaryotic clades in the LSC1 but not in LSC2 phylogeny. While in the LSC1 phylogeny Alphaproteobacteria form a monophyletic group with eukaryotes (**Figure S35C**), in the LSC2 phylogeny other bacteria (e.g. Verrucomicrobiota, 95% UfBoot2) are placed between eukaryotes and a group of diverse bacterial lineages including Alphaproteobacteria (**Figure S35D**). In turn, while the origin of LSC2 is tentative, we suggest that eukaryotic LCS1/2 have an alphaproteobacterial origin.

**Succinate dehydrogenase complex — SdhABCD — K00234/K00239/K00244, K00235/K00240/K00245, K00236/K00237/K00241/K00246**

Succinate dehydrogenase complex, SdhABCD, is composed of four subunits, which function in the TCA as well as in the electron transport chain of the mitochondria. SdhABCD catalyzes the oxidation of succinate to fumarate, coupled to the reduction of ubiquinone to ubiquinol. The phylogeny of the four Sdh subunits, show a patchy distribution in bacteria and archaea, but are widespread in eukaryotes, whose sequences form a monophyletical cluster branching with alphaproteobacterial sequences (**Figure S36A-C**). SdhC and SdhD subunits are around 150 amino acids long, with both originating from a duplication that can be traced back to prokaryotes (**Figure S36C**). The phylogeny of SdhD is slightly less conclusive but an alphaproteobacterial origin seems possible (**Figure S36C**). The absence of SDHC/D subunits in Diaphoretickes taxa in our dataset might be a result of limitations in homology detection because the presence of a complete SDH complex has been reported for Alveolates; (Silva et al. 2023)). Altogether, the SdhABCD complex was present in LECA and was derived from the alphaproteobacterial partner.

**Fumarate hydratases — FumAB|FumC — K01676, K01679**

Fumarate hydratases are classified as FumAB and FumC which are distant homologs found in prokaryotes and eukaryotes. Eukaryotic FumAB form one main monophyletic clade comprising taxa from all eukaryotic supergroups but with mixed topology, possibly due to poor phylogenetic signal (**Figure S37A**). In fact, Kinetoplastea sequences are branching closely with bacteria, but when we include a divergent outgroup in the phylogeny, these Kinetoplastea sequences branch within the main eukaryotic group (inset tree in **Figure S37A**). Yet the wide composition of this eukaryotic group suggests that FumAB was present in LECA. FumAB has a limited distribution in bacteria, and bootstrap support values do not allow to infer a clear sister group, which comprises bacteria like Planctomycetota, Alphaproteobacteria, Myxococcota, Nitrospirota and Tectomicrobia, among others.

In contrast to FumAB, the phylogeny of FumC shows a wide distribution in prokaryotes, including Asgardarchaeota and Alphaproteobacteria (**Figure S37B**). Eukaryotic sequences form a main monophyletic group comprising all eukaryotic diversity, with Alphaproteobacteria as sister group. Thus, FumC is inferred to have been present in LECA and most likely had an alphaproteobacterial origin.

**Lactate dehydrogenase and Malate dehydrogenases — Ldh|Mdh|Mdh1|Mdh2 — K00016, K00024, K00025, K00026**

The L-lactate dehydrogenase (LDH) and malate dehydrogenases (MDH) are homologous enzymes that were merged in our phylogenetic analyses similar to previous work (Schnarrenberger and Martin 2002; Brochier-Armanet and Madern 2021; Robin et al. 2023).

Our results show that LDH is mainly found in bacteria and a limited number of archaea (**Figure S38A**). Eukaryotic sequences form two main clades with mixed taxonomic compositions. While one group comprises Metazoa, SAR and Chloroplastida-Rhodophyta with Cyanobacteria as sister group, the other group mainly comprise Fungi, SAR and Rhodophytes, being closely related to Dependentiae bacterial sequences (**Figure S38B**). There is no evidence for shared introns between the two eukaryotic clades (**Figure S38C**), suggesting distinct origins. While the sister-relationship with Cyanobacteria has been previously observed (Robin et al. 2023), the Dependentiae sister relationship was undetected. Thus, considering that such sister relationships are well supported (>99% optimized UfBoot2), these eukaryotic clades might represent two LECA acquisitions from Cyanobacteria and Dependintae respectively.

There are different paralogous families of malate dehydrogenases, which are referred to as MDH, MDH1, and MDH2 (LDH, described in *Pyruvate metabolisms*). The phylogeny of this protein superfamily including MDH, MDH1, MDH2 and LDH, shows that MDH is mostly found in prokaryotes and is closely related to the LDH (**Figure S38A**). On the other hand, MDH1 and MDH2, are mainly found in eukaryotes, and distantly related to MDH/LDH (**Figure S38A**). The wide taxonomic composition of MDH1 and MDH2 clades (**Figure S38B**), suggests that both paralogs were present in LECA. Due to the scarcity of prokaryotes in MDH1 and MDH2 clades and their phylogenetic position within eukaryotes, it is possible that both families originated in eukaryotes, and subsequently transferred to bacteria. In fact, both MDH1 and MDH2 subfamilies share introns positions (introns 1511, 1581, 1601, among others, **Figure S38D**), further supporting the LECA duplication. A clade composed of TACK and Baldarchaeota sequences branch basal to the MDH1/2 bifurcation (**Figure S38A**), suggesting that MDH1/2 may be of archaeal origin. However, given the long branches substending both MDH1 and MDH2, this inference remains speculative and would suggest an early duplication event followed by rapid evolution of the two families prior to LECA.

Importantly, among the bacterial sequences branching within the MDH1 family, a group of aerobic Chlamydia clusters together with the Archaeplastida-Chlorophyta clade with full support (**Figure S38B**). This relationship is in agreement with previous observations regarding Archaeplastida and Chlamydiota (Becker et al. 2008). Since Chlorophyta sequences are branching within bacterial clades, but some shared introns suggest LECA origin of Chlorophyta sequences (introns 1581 and 2341 **Figure S38D**), the direction of HGT between Chlamydia and Chlorophyta is unclear. However, given that MDH1 and MDH2 most likely originated by gene duplication in LECA it is possible that bacterial MDH1 sequences originated through eukaryote-to-bacteria HGT. Therefore, MDH phylogeny may illustrate a gene escape during eukaryogenesis, as previously observed for other eukaryotic components like histone-related genes transferred from proto-eukaryotes to viruses (Irwin and Richards 2023).

**Isocitrate lyase — AceA — K01637**

The phylogeny of isocitrate lyase, AceA, is a distinctive eukaryotic clade defined by a long basal branch (**Figure S39**). This clade is composed by a large diversity of eukaryotes suggesting that AceA was present in LECA. The relationship between the eukaryotic clade and the sister group is unclear due to long basal branches. Despite this, it is most likely that eukaryotic AceA was acquired from bacteria.

**Correlated orthogroups of central carbon metabolism enzymes**

To infer the main trends in the evolution of CCM based on absence/presence profiles, we analyzed the correlative distribution between selected orthogroups in combination with the different lifestyles (anaerobic, primary and secondary endosymbiosis; **Figure S41, TABLE**). The correlative network at 0.5 phi-coefficient cut-off, provided three main informative clusters (**Figure 4AB**).

The glucose phosphorylation step is alternately performed by HK or GLK/ROK (**Figure 4AB**). While HK is inferred to be present in LECA, GLK phylogeny suggests independent post-LECA acquisitions that might have replaced HK in certain lineages and potentially explaining the anticorrelation patterns. In addition, the distribution of GLK, correlates with the distribution of ROK which is inferred to be present in LECA. The targeting signal of these reveals specific functionalization between the paralgos specially in Archaeplastida and Rhizaria (**Figure 4B**).

The enzymatic composition of pyruvate/acetate and TCA metabolisms of eukaryotes reflect their anaerobic or aerobic lifestyles, respectively (**Figure 4AB**). Putative anaerobic adaptations seem to have occurred in Metamonada, Breviatea and other protists. POR is anticorrelated with the PDH complex enzymes (except for the PDHD subunit, which is a multifunctional enzyme). Additionally, GPI, FBA, and RPIB and AcdAB phylogenies also displayed gene acquisitions including in anaerobic eukaryotes (**Figure 4AB, S41**).

The third cluster is related with photosynthetic endosymbioses (primary and/or secondary) mainly including enzymes from glycolysis PPP, and PDH complex (**Figure 4AB**). In Archaeplastida, the proteins of the orthogroups forming these clusters are usually targeted to the chloroplast, while in secondary endosymbionts the targeting signal is mixed as they present a mix of chloroplast and mitochondrial targeting (Figure 4B). The chloroplast orthogroups of RPE and TKT contain sequences from Rhodelphis that are targeted to the chloroplast (**Figure 4B**). This result is in agreement with the presence of chloroplast remnants in this group as previously pointed out (Gawryluk et al. 2019). By contrast, we did not find any chloroplast contributions to Picozoa.

**Origins and evolution of parallel EMP glycolysis and PPP in Archaeplastida**

Paralogous glycolytic enzymes that are targeted to the cytoplasm and chloroplast respectively, are found in photosynthetic Archaeplastida clades, and intermittently in SAR protists, with variation between relatives (**Figure S42**), illustrating the heterogeneous conservation of both glycolytic and pentose-phosphate pathways.

Our examination of the evolutionary history of glycolysis and PPP illustrates the complex evolution of these pathways in Archaeplastida (**Figure S43**). First, enzymes originating from LECA such as GPI, PFK, ALDO, TPI, ENO, PK, H6PD, PGLS and PGD, seem to have been subject to duplications (**Figures S5-8,13-15**). However, the phylogenetic evidence supporting the ancestrality of these duplications is not always unequivocal, and therefore, most of these duplications might have arisen independently in specific Archaeplastida lineages. Secondly, according to the potential cyanobacterial acquisitions, GPI, FBA, GAPDH, PGK, RPE, EDD, EDA, TKT, RPIA and PRPS (**Figures S5,7,9,10,19,20,22,23,24**) were likely present in the Archaeplastida ancestor, although cases like GPI, FBA, and RPIA, might have been lost multiple times during Archaeplastida evolution, as these are retained in unique groups like Glaucophyta and Streptophyta. These chloroplast remnants might compensate for the loss of the LECA paralog duplicates. And thirdly, the chloroplast acquisitions of PGK, TKT, and PRPS led to the replacement of LECA paralogs within Archaeplastida clades (**Figure S10,22,24**).

Collectively, Archaeplastida exhibits distinct mechanisms for acquiring paralogous genes, including nuclear gene duplications, EGT, and HGTs, which have subsequently given rise to intricate evolutionary trajectories. These trajectories have resulted in the heterogeneous maintenance of both plastidial and cytoplasmic versions of glycolysis and the PPP, illustrating the general prevalence of duplications of nuclear genes over the genes originating from chloroplast and therefore, highlights the relocation of 'nuclear' genes to the photosynthetic organelle.

**Later stage of mitochondrial glycolysis is specific of Diaphoretickes protists**

Enzymes catalyzing steps in the second half of glycolysis (C3 conversions), have been observed to be targeted to the mitochondria in Stramenopiles (Liaud et al. 2000; Río Bártulos et al. 2018b). We made a survey of targeting signal across all EukProt V3 proteomes (~1,100 taxa) and observed that glycolytic enzymes with paralogs targeted to both the cytoplasm and the mitochondria, are predominantly found in SAR and Haptista (**Figure S42**). These targeting signals are preferentially found in enzymes utilizing C3 substrates (from TPI to PK), although with heterogeneous pattern of conservation between close relatives (**Figure S42,44A**). The targeted enzymes usually belong to LECA clades (eg. PGK, ENO), but their targeting signals distribution in combination with the phylogeny do not indicate a common origin. Additionally, the PGK phylogeny indicates that a gene product that was targeted to the chloroplast switched to mitochondrial targeting after secondary endosymbiosis in the Stramenopiles-Ochorphyta ancestor (**Figure S44B-D**). Therefore, mitochondria glycolysis seems to be specific of Diaphoretickes protist, involving the later steps utilizing triose substrates, and most likely arose independently in certain lineages.

### References

Atkins MS, McArthur AG, Teske AP. 2000. Ancyromonadida: A New Phylogenetic Lineage Among the Protozoa Closely Related to the Common Ancestor of Metazoans, Fungi, and Choanoflagellates (Opisthokonta). *J. Mol. Evol.* 51:278–285.

Becker B, Hoef-Emden K, Melkonian M. 2008. Chlamydial genes shed light on the evolution of photoautotrophic eukaryotes. *BMC Evol. Biol.* 8:203.

Brochier-Armanet C, Madern D. 2021. Phylogenetics and biochemistry elucidate the evolutionary link between l-malate and l-lactate dehydrogenases and disclose an intermediate group of sequences with mix functional properties. *Biochimie* 191:140–153.

Burki F. 2016. Mitochondrial Evolution: Going, Going, Gone. *Curr. Biol.* 26:R410–R412.

Chen X, Schreiber K, Appel J, Makowka A, Fähnrich B, Roettger M, Hajirezaei MR, Sönnichsen FD, Schönheit P, Martin WF, et al. 2016. The Entner–Doudoroff pathway is an overlooked glycolytic route in cyanobacteria and plants. *Proc. Natl. Acad. Sci.* 113:5441–5446.

Foster JM, Davis PJ, Raverdy S, Sibley MH, Raleigh EA, Kumar S, Carlow CKS. 2010. Evolution of Bacterial Phosphoglycerate Mutases: Non-Homologous Isofunctional Enzymes Undergoing Gene Losses, Gains and Lateral Transfers. *PLOS ONE* 5:e13576.

Gawryluk RMR, Eme L, Roger AJ. 2015. Gene fusion, fission, lateral transfer, and loss: Not-so-rare events in the evolution of eukaryotic ATP citrate lyase. *Mol. Phylogenet. Evol.* 91:12–16.

Gawryluk RMR, Stairs CW. 2021. Diversity of electron transport chains in anaerobic protists. *Biochim. Biophys. Acta BBA - Bioenerg.* 1862:148334.

Gawryluk RMR, Tikhonenkov DV, Hehenberger E, Husnik F, Mylnikov AP, Keeling PJ. 2019. Non-photosynthetic predators are sister to red algae. *Nature* 572:240–243.

Grauvogel C, Brinkmann H, Petersen J. 2007. Evolution of the Glucose-6-Phosphate Isomerase: The Plasticity of Primary Metabolism in Photosynthetic Eukaryotes. *Mol. Biol. Evol.* 24:1611–1621.

Hampl V, Hug L, Leigh JW, Dacks JB, Lang BF, Simpson AGB, Roger AJ. 2009. Phylogenomic analyses support the monophyly of Excavata and resolve relationships among eukaryotic “supergroups.” *Proc. Natl. Acad. Sci.* 106:3859–3864.

Hannaert V, Brinkmann H, Nowitzki U, Lee JA, Albert M-A, Sensen CW, Gaasterland T, M M, Michels P, Martin W. 2000. Enolase from Trypanosoma brucei, from the Amitochondriate Protist Mastigamoeba balamuthi, and from the Chloroplast and Cytosol of Euglena gracilis: Pieces in the Evolutionary Puzzle of the Eukaryotic Glycolytic Pathway. *Mol. Biol. Evol.* 17:989–1000.

Heiss AA, Kolisko M, Ekelund F, Brown MW, Roger AJ, Simpson AGB. 2018. Combined morphological and phylogenomic re-examination of malawimonads, a critical taxon for inferring the evolutionary history of eukaryotes. *R. Soc. Open Sci.* 5:171707.

Hoang DT, Chernomor O, von Haeseler A, Minh BQ, Vinh LS. 2018. UFBoot2: Improving the Ultrafast Bootstrap Approximation. *Mol. Biol. Evol.* 35:518–522.

Horecker BL. 2002. The Pentose Phosphate Pathway. *J. Biol. Chem.* 277:47965–47971.

Irwin NAT, Richards TA. 2023. Self-assembling viral histones unravel early nucleosome evolution. :2023.09.20.558576. Available from: https://www.biorxiv.org/content/10.1101/2023.09.20.558576v1

Karnkowska A, Vacek V, Zubáčová Z, Treitli SC, Petrželková R, Eme L, Novák L, Žárský V, Barlow LD, Herman EK, et al. 2016. A Eukaryote without a Mitochondrial Organelle. *Curr. Biol.* 26:1274–1284.

Keeling PJ, Doolittle WF. 1997. Evidence that eukaryotic triosephosphate isomerase is of alpha-proteobacterial origin. *Proc. Natl. Acad. Sci.* 94:1270–1275.

Krebs HA, Johnson WA. 1937. Metabolism of ketonic acids in animal tissues. *Biochem. J.* 31:645–660.

Kresge N, Simoni RD, Hill RL. 2005. Otto Fritz Meyerhof and the Elucidation of the Glycolytic Pathway. *J. Biol. Chem.* 280:e3.

Lartillot N, Lepage T, Blanquart S. 2009. PhyloBayes 3: a Bayesian software package for phylogenetic

reconstruction and molecular dating. *Bioinformatics* 25:2286–2288.
Liaud MF, Lichtlé C, Apt K, Martin W, Cerff R. 2000. Compartment-specific isoforms of TPI and GAPDH are imported into diatom mitochondria as a fusion protein: evidence in favor of a mitochondrial origin of the eukaryotic glycolytic pathway. *Mol. Biol. Evol.* 17:213–223.
Martin W, Brinkmann H, Savonna C, Cerff R. 1993. Evidence for a chimeric nature of nuclear genomes: eubacterial origin of eukaryotic glyceraldehyde-3-phosphate dehydrogenase genes. *Proc. Natl. Acad. Sci. U. S. A.* 90:8692–8696.
Minh BQ, Schmidt HA, Chernomor O, Schrempf D, Woodhams MD, von Haeseler A, Lanfear R. 2020. IQ-TREE 2: New Models and Efficient Methods for Phylogenetic Inference in the Genomic Era. *Mol. Biol. Evol.* 37:1530–1534.
More K, Klinger CM, Barlow LD, Dacks JB. 2020. Evolution and Natural History of Membrane Trafficking in Eukaryotes. *Curr. Biol.* 30:R553–R564.
Müller M, Mentel M, van Hellemond JJ, Henze K, Woehle C, Gould SB, Yu R-Y, van der Giezen M, Tielens AGM, Martin WF. 2012. Biochemistry and Evolution of Anaerobic Energy Metabolism in Eukaryotes. *Microbiol. Mol. Biol. Rev.* 76:444–495.
Novák LVF, Treitli SC, Pyrih J, Hałakuc P, Pipaliya SV, Vacek V, Brzoň O, Soukal P, Eme L, Dacks JB, et al. 2023. Genomics of Preaxostyla Flagellates Illuminates the Path Towards the Loss of Mitochondria. *PLOS Genet.* 19:e1011050.
Pittis AA, Gabaldón T. 2016. Late acquisition of mitochondria by a host with chimaeric prokaryotic ancestry. *Nature* 531:101–104.
Plaxton WC. 1996. The Organization and Regulation of Plant Glycolysis. *Annu. Rev. Plant Physiol. Plant Mol. Biol.* 47:185–214.
Río Bártulos C, Rogers MB, Williams TA, Gentekaki E, Brinkmann H, Cerff R, Liaud M-F, Hehl AB, Yarlett NR, Gruber A, et al. 2018a. Mitochondrial Glycolysis in a Major Lineage of Eukaryotes. *Genome Biol. Evol.* 10:2310–2325.
Río Bártulos C, Rogers MB, Williams TA, Gentekaki E, Brinkmann H, Cerff R, Liaud M-F, Hehl AB, Yarlett NR, Gruber A, et al. 2018b. Mitochondrial Glycolysis in a Major Lineage of Eukaryotes. *Genome Biol. Evol.* 10:2310–2325.
Robin AY, Brochier-Armanet C, Bertrand Q, Barette C, Girard E, Madern D. 2023. Deciphering Evolutionary Trajectories of Lactate Dehydrogenases Provides New Insights into Allostery. *Mol. Biol. Evol.* 40:msad223.
Ronimus RS, Morgan HW. 2004. Cloning and biochemical characterization of a novel mouse ADP-dependent glucokinase. *Biochem. Biophys. Res. Commun.* 315:652–658.
Schnarrenberger C, Martin W. 2002. Evolution of the enzymes of the citric acid cycle and the glyoxylate cycle of higher plants. A case study of endosymbiotic gene transfer. *Eur. J. Biochem.* 269:868–883.
Sheridan PO, Meng Y, Williams TA, Gubry-Rangin C. 2022. Recovery of Lutacidiplasmatales archaeal order genomes suggests convergent evolution in Thermoplasmatota. *Nat. Commun.* 13:4110.
Silva MF, Douglas K, Sandalli S, Maclean AE, Sheiner L. 2023. Functional and biochemical characterization of the Toxoplasma gondii succinate dehydrogenase complex. *PLoS Pathog.* 19:e1011867.
Stairs CW, Dharamshi JE, Tamarit D, Eme L, Jørgensen SL, Spang A, Ettema TJG. 2020. Chlamydial contribution to anaerobic metabolism during eukaryotic evolution. *Sci. Adv.* 6:eabb7258.
Stairs CW, Leger MM, Roger AJ. 2015. Diversity and origins of anaerobic metabolism in mitochondria and related organelles. *Philos. Trans. R. Soc. Lond. B. Biol. Sci.* 370:20140326.
Stairs CW, Roger AJ, Hampl V. 2011. Eukaryotic Pyruvate Formate Lyase and Its Activating Enzyme Were Acquired Laterally from a Firmicute. *Mol. Biol. Evol.* 28:2087–2099.
Stairs CW, Táborský P, Salomaki ED, Kolisko M, Pánek T, Eme L, Hradilová M, Vlček Č, Jerlström-Hultqvist J, Roger AJ, et al. 2021. Anaeramoebae are a divergent lineage of eukaryotes that shed light on the transition from anaerobic mitochondria to hydrogenosomes. *Curr. Biol.* 31:5605-5612.e5.

Stechmann A, Baumgartner M, Silberman JD, Roger AJ. 2006. The glycolytic pathway of Trimastix pyriformis is an evolutionary mosaic. *BMC Evol. Biol.* 6:101.
Stoddard PR, Lynch EM, Farrell DP, Dosey AM, DiMaio F, Williams TA, Kollman JM, Murray AW, Garner EC. 2020. Polymerization in the actin ATPase clan regulates hexokinase activity in yeast. *Science* 367:1039–1042.
Strassert JFH, Irisarri I, Williams TA, Burki F. 2021. A molecular timescale for eukaryote evolution with implications for the origin of red algal-derived plastids. *Nat. Commun.* 12:1879.
Takishita K, Inagaki Y. 2009. Eukaryotic origin of glyceraldehyde-3-phosphate dehydrogenase genes in Clostridium thermocellum and Clostridium cellulolyticum genomes and putative fates of the exogenous gene in the subsequent genome evolution. *Gene* 441:22–27.
Tikhonenkov DV, Mikhailov KV, Gawryluk RMR, Belyaev AO, Mathur V, Karpov SA, Zagumyonnyi DG, Borodina AS, Prokina KI, Mylnikov AP, et al. 2022. Microbial predators form a new supergroup of eukaryotes. *Nature* 612:714–719.
Verhees CH, Kengen SWM, Tuininga JE, Schut GJ, Adams MWW, De Vos WM, Van Der Oost J. 2003. The unique features of glycolytic pathways in Archaea. *Biochem. J.* 375:231–246.
Vosseberg J, Schinkel M, Gremmen S, Snel B. 2022. The spread of the first introns in proto-eukaryotic paralogs. *Commun. Biol.* 5:1–9.
Wang Y, Chen X, Spengler K, Terberger K, Boehm M, Appel J, Barske T, Timm S, Battchikova N, Hagemann M, et al. 2022. Pyruvate:ferredoxin oxidoreductase and low abundant ferredoxins support aerobic photomixotrophic growth in cyanobacteria.Kramer DM, Storz G, Ducat DC, Burnap R, Nitschke W, editors. *eLife* 11:e71339.
Williams SK, Hultqvist JJ, Eglit Y, Salas-Leiva DE, Curtis B, Orr R, Stairs CW, Simpson AGB, Roger AJ. 2023. Extreme mitochondrial reduction in a novel group of free-living metamonads. :2023.05.03.539051. Available from: https://www.biorxiv.org/content/10.1101/2023.05.03.539051v1
Wu F, Speth DR, Philosof A, Crémière A, Narayanan A, Barco RA, Connon SA, Amend JP, Antoshechkin IA, Orphan VJ. 2022. Unique mobile elements and scalable gene flow at the prokaryote–eukaryote boundary revealed by circularized Asgard archaea genomes. *Nat. Microbiol.* 7:200–212.

**Supplementary Table Captions**

**Table S1.** Taxonomy information of the core and extended dataset.

**Table S2.** Eukaryotic contaminations detected, and general statistics of Kinetoplatid contamination using EggNog/NCBI annotations and a custom ETE script.

**Table S3.** Selection of KO identifiers investigated in this study. Sheet 1, *Selected KOs*, contains the KOs that define the central carbon metabolism in eukaryotes. Sheet 2, *Selection of KOs present in eukaryotes*, distribution of all central carbon metabolism KOs in the core dataset. From here, we selected the KOs of interest based on their presence in eukaryotes.

**Table S4.** KO homologies based on comparison of the respective hidden Markov models. HMM comparisons were made through the HH-suite package, and only the homology of those KO of interest are shown. Groups delimited with borders indicate those KOs that were merged in further phylogenetic analyses.

**Table S5.** Data for the phylogenetic profiles and correlative distributions. This table contains the orthogroups manually selected from the phylogenetic trees, including the potential targeting and the respective sequences. Note that the names of the orthogroups are descriptive and should not be interpreted as final inference. Check Supplementary information for additional discussion about the evolution origins of the orthogroups.

**Table S6.** Ultrafast bootstrap surveys for eukaryotic tree of life (sheet 1) and GPI's multiple sequence alignment partitions (sheet 2). Original scripts for these surveys are provided at Zenodo repository (https://zenodo.org/records/10991068).

**Supplementary Figure Captions**

**Figure S1.** Extended phylogenetic analyses for the reconstruction of the eukaryotic tree of life (eToL) based on the concatenation of 317 phylogenetic markers. **A)** Maximum-likelihood reconstruction using IQ-TREE and ultrafast bootstraps (Ufboot2). **B)** Bayesian reconstruction of eToL using PhyloBayes. **C)** Maximum-likelihood phylogenies of eToL after applying gradual filtering of heterogeneous sites to the concatenation (from 0.1 to 0.9). These phylogenies are used for depicting Ufboot2 values in figure 1B. **D)** Expanded view of maximum-likelihood reconstruction using IQ-TREE and corrected Ufboot2 used in Figure 1A. Phylogenetic methods and length of the concatenations are indicated in the respective panels. BUSCO values and percentage of phylogenetic markers found in each eukaryotic proteome provided in **Table S1**.

**Figure S2.** Workflow for generating final phylogenies of central carbon metabolism enzymes for the identification of sister groups and the taxonomic composition of orthogroups. Lens indicate steps that were manually supervised. See methods for further explanation.

**Figure S3.** Phylogeny of P-loop NTPases involved in glucose phosphorylation. **A)** Phylogeny of glucokinase (GLK) rooted at the ROK subfamily (K00845). The multiple sequence alignment (MSA) was trimmed with trimAl (-gt 0.4) resulting in 348 positions. **B)** Phylogeny of ROK subfamily rooted arbitrarily at the separation between ROK and ROK/GLK subfamilies. The MSA consisted of 263 positions. **C)** Phylogeny of PPGK (K00886), and respective MSA of 227 positions. **D)** Phylogeny of Hexokinase (K00844) and respective MSA of 438 positions. The tree was rooted arbitrarily at a potential ancestral group of Firmicutes. **E)** Unrooted view of GLK/ROK (K00845) protein family, showing the distance between eukaryotic ROK and GLK. The same tree shown in **B)**. Bootstraps higher than 95% are denoted with black dots. **F)** Global phylogeny of the closest P-loop NTPases to the enzymes studied. The final MSA resulted in 254 positions. Both represent the same phylogeny but branches are colored according to the taxonomy (left panel) and the protein family (KO, right panel). Bootstraps higher than 95% are denoted with black dots. Asterisks in **D)** and **F)** panel indicate a group of Metamonada and bacteria with unstable topologies. Green highlight indicates a potential LECA clade.

**Figure S4.** Phylogeny of ADP-dependent glucokinase (ADPGK, K08074) and its closest family ADP-dependent phosphofructokinase/glucokinase (PFKC, K00918). The tree was rooted at midpoint and the MSA was trimmed with trimAl (-gt 0.4) resulting in 446 positions. Green highlight indicates a potential LECA clade.

**Figure S5. A)** Unrooted and **B)** rooted view of glucose-6-phosphate isomerase phylogeny (GPI, K01810). The final MSA resulted in 443 positions. The tree was arbitrarily rooted to ease visualization (arrow). **C)** Conservation of introns along GPI MSA using a selected set of eukaryotic genomes (*Eukarya4 (Vosseberg et al)*, see methods) mapped onto GPI phylogeny including relative prokaryotic sequences. Column labels denote the position in the MSA and the respective intron phases identified. Those highlighted positions indicate potential LECA introns. Bars at the right show the intron density for each coding gene. **D)** Phylogenies of the flanking region (+/- 20 positions) to the intron at position 1915 (left), and half C-terminus of the MSA from position 2117. This figure shows the different topologies of Cryptista sequences (and Chlamydia) that share introns with plastid paralog of Streptophyta. **E)** Ultrafast bootstrap dynamics of Cryptista-plastid and Cryptista-'LECA' monophyly along GPI's MSA partitions. Webologo for those partitions with potential Cryptista-plastid signals are shown.

**Figure S6. A)** Phylogeny of 6-phosphofructokinase 1 (PFKA, K00850) including closest homologs and protein domain architecture. PFKA phylogeny defines two paraphyletic groups, PFKA_1 with double tandem repeat PFK domain specific to eukaryotes and few bacteria (right panel for zoom in), and PFKA_2 with usually a single domain. **B)** Unrooted views of phylogenies using full protein MSA and extracted PFK Pfam domains MSAs (left and right panel respectively). Branch colors indicate gene family (upper panel) and taxonomy levels (lower panel). **C)** Full protein phylogeny of PFKA_1 rooted at PFKA_2. Left panel shows a phylogeny of the extracted PFK domain including prokaryotic sequences expanded searches, showing the spread of PFKA_1 double domain in Bacteria, in which Actinobacteria are closely related to Amoebozoa sequences. **D)** phylogeny of PFKA_2 rooted at PFKA_1. The final MSA consisted of 325 positions.

**Figure S7.** Phylogeny of analogous aldolases involved in glycolysis. Unrooted views of **A)** fructose-bisphosphate aldolase, class I (ALDO, K01623) and **B)** fructose-bisphosphate aldolase, class II (FBA, K01624) phylogenies and respective rooted views **C)** and **D)**. ALDO phylogeny was arbitrarily rooted at a distant bacterial group to the eukaryotic sequences and FBA phylogeny was rooted at midpoint. Some branches containing only prokaryotic sequences are collapsed to ease visualization, but can be identified in the unrooted views. The MSAs of ALDO and FBA reconstructions consisted of 343 and 282 positions respectively. The MSA of FBA was trimmed with trimAl (-gt 0.4).

**Figure S8.** Phylogenies of triosephosphate isomerase (TPI, K01803) **A)** including and **B)** and **C)** excluding archaeal sequences. The tree in **C)** was rooted arbitrarily in a group composed only by bacterial sequences. The arrow indicates the selected root. The MSAs consisted of 247 positions (full set, trimAl -gt 0.7) and 256 positions (bacteria-eukaryote set, trimAl -gt 0.4).

**Figure S9. A)** Unrooted and **B)** rooted views of glyceraldehyde 3-phosphate dehydrogenase phylogeny (GAPDH, K00134). The final MSA resulted in 443 positions. The tree was arbitrarily rooted to ease visualization (arrow). Green highlight indicates potential LECA clade.

**Figure S10. A)** Unrooted and **B)** rooted views of phosphoglycerate kinase phylogeny (PGK, K00927). The final MSA consisted of 392 positions. The tree was arbitrarily rooted to ease visualization (arrow).

**Figure S11.** Phylogenies of cofactor-dependent phosphoglycerate mutases. Unrooted views of **A)** GPMA (K01834) and **B)** GPMB (K15634) phylogenies. The respective MSAs consisted of 240 and 169 positions.

**Figure S12.** Phylogenies of cofactor-independent phosphoglycerate mutases. **A)** Unrooted and rooted phylogeny of GPMI (K15633). The rooted phylogeny was rooted at mid-point, and the respective MSA consisted of 508 positions. The tree was rooted at mid-point (arrow). **B)** Unrooted and rooted phylogeny of APGM (K15635). The rooted phylogeny was rooted arbitrarily at the distant groups of prokaryotes to the eukaryotic group containing bacterial and eukaryotic sequences. The respective MSA consisted of 400 positions. The tree was arbitrarily rooted to ease visualization (arrow).

**Figure S13. A)** Unrooted view of Enolase phylogeny (ENO, K01689) and rooted view of two eukaryotic ENO clades identified, **B)** ENO_1 and **C)** ENO_2. The tree was arbitrarily rooted between the split of eukaryotic-archaeal and bacterial sequences. The final MSA consisted of 419 positions.

**Figure S14. A)** Unrooted phylogeny of pyruvate kinase (PK, K00873) highlighting the eukaryotic groups discussed in this study (PK_1 to PK_6). **B-E)** Rooted phylogeny of different PK_1-6 groups. All rooted trees are rooted at the same midpoint node (arrow). Left panel in **D)** shows the same phylogenetic group (PK_5) including sequences from searches against all eukaryotic proteomes of EUKPROTv3. The final MSA consisted of 469 positions.

**Figure S15. A)** Unrooted and **B)** rooted view of glucose-6-phosphate 1-dehydrogenase (G6PD, K00036) and hexose-6-phosphate dehydrogenase (H6PD, K13937) phylogeny including domain architecture. The final MSA consisted of 479 positions. **C)** Unrooted and **D)** rooted view of 6-phosphogluconolactonase (PGLS, K01057) phylogeny. The final MSA consisted of 234 positions. Independent fusions of G6PD and PGLS are indicated. The trees were arbitrarily rooted to ease visualization (arrow). Asterisks indicate an independent fusion in Metamonada.

**Figure S16.** Phylogeny of NAD+ dependent glucose-6-phosphate dehydrogenase (AZF, K19243). The final MSA consisted of 238 positions.

**Figure S17. A)** Unrooted and **B)** rooted view of glucose-6-phosphate 1-dehydrogenase (PGL, K07404) phylogeny. The tree was arbitrarily rooted to ease visualization (arrow). The final MSA consisted of 240 positions.

**Figure S18. A)** Unrooted and **B)** rooted view of 6-phosphogluconate dehydrogenase phylogeny (PGD, K00033). The tree was rooted at mid point (arrow). The final MSA consisted of 311 positions. **C)**

Conservation of introns along PGD MSA using a selected set of eukaryotic genomes (*Eukarya4 (Vosseberg et al)*, see methods) mapped onto PGD phylogeny including relative prokaryotic sequences. Column labels denote the position in the MSA and the respective intron phases identified. Bars at the right show the intron density for each coding gene.

**Figure S19. A)** Unrooted and **B)** rooted view of ribulose-phosphate 3-epimerase phylogeny (RPE, K01783). The tree was arbitrarily rooted to ease visualization (arrow). The final MSA consisted of 214 positions. **C)** Refined phylogeny of RPE subtree showing the external clustering of bacterial sequences that cluster within LECA clade in **B)** (see arrow). **D)** Phylogeny of RPE with empirical models (LG+R10) showing clear signal of plastid paralog with Cyanoabcteria.

**Figure S20. A)** Unrooted view of ribose 5-phosphate isomerase A phylogeny (RPIA, K01807). The tree was arbitrarily rooted to ease visualization (arrow). Rooted subtrees of eukaryotic **B)** RPIA_1 and **C)** RPIA_2 clades. The final MSA consisted of 227 positions.

**Figure S21. A)** Unrooted and **B)** rooted view of ribose 5-phosphate isomerase B phylogeny (RPIB, K01808). The tree was arbitrarily rooted to ease visualization (arrow). The final MSA consisted of 146 positions.

**Figure S22.** Unrooted views of **A)** fructose-bisphosphate aldolase, class I (TKTA/B, K00615) and **B)** transaldolase (TALA/B, K00616) phylogenies and respective rooted views of refined phylogenies **C)** and **D)**. Trees were rooted arbitrarily to ease visualization. The MSAs for final phylogenies (**C**, and **D**) consisted of 656 and 319 positions respectively.

**Figure S23. A)** Unrooted and **B)** rooted view of phosphogluconate dehydratase phylogeny (EDD, K01690) and closest relative gene families IlvD (K00845) and XylD (K00886). Upper and lower panels in **A**), are the same phylogeny, coloring the branches by taxonomy and gene family respectively. The tree in **B)** was arbitrarily rooted to ease visualization (arrow). The final MSA consisted of 549 positions. **C)** Rooted phylogeny of 2-dehydro-3-deoxyphosphogluconate aldolase (EDA, K01625). The tree was rooted at midpoint (arrow). The final MSA consisted of 211 positions.

**Figure S24. A)** Unrooted phylogeny of ribose-phosphate pyrophosphokinase (PRPS, K00948) highlighting the eukaryotic groups discussed in this study (from PRPS_1 to PRPS_4). **B-D)** Rooted phylogenies of different PRPS_1-4 groups. All trees are rooted at the same node which was selected arbitrarily to ease visualization. The final MSA consisted of 296 positions.

**Figure S25.** Phylogenies of pyruvate dehydrogenase subunits. Unrooted (upper panels) and rooted (lower panels) views of **A)** PDHA, **B)** PDHB, **C)** PDHC and PDHX, and **D)** PDHD (dihydrolipoyl dehydrogenase). The trees were arbitrarily rooted to ease visualization. The final MSAs consisted of 330, 325, 387 and 458 positions respectively. Tree at the bottom in **D)** with an asterisk shows a refined phylogeny of mitochondrial PDHD using a selection of eukaryotes and bacterial sequences. Labels in bold are those sequences whose topology changes, branching within or outside (with alphaproteobacteria) the LECA clade. Bars shows the P-values for the chi2 test, computed by IQ-TREE. Those sequences with P-value lower than 5% might contain compositional bias.

**Figure S26. A)** Unrooted and **B)** rooted view of pyruvate-ferredoxin/flavodoxin oxidoreductase phylogeny (POR, K03737). The tree was arbitrarily rooted to ease visualization (arrow). The final MSA consisted of 1166 positions.

**Figure S27. A)** Unrooted and **B)** rooted view of formate C-acetyltransferase (PFL, K00656) phylogeny. The tree was rooted at midpoint visualization (arrow). The final MSA consisted of 690 positions.

**Figure S28. A)** Unrooted and **B)** rooted view of acetyl-CoA synthetase phylogeny (ACS, K01895). The tree was arbitrarily rooted to ease visualization (arrow). The final MSA consisted of 627 positions. **C)** Conservation of introns along ACS MSA using a selected set of eukaryotic genomes (*Eukarya4 (Vosseberg et al)*, see methods) mapped onto ACS phylogeny including relative prokaryotic sequences. Column labels denote the position in the MSA and the respective intron phases identified. Bars at the right show the intron density for each coding gene. Those positions that share introns between two ACS paralogs are shown.

**Figure S29.** Phylogenies and domain architecture of acetate---CoA ligase (ADP-forming) (ACDA/B/AB, K22224, K01905 and K24012). **A)** Rooted phylogeny of extracted ATP-grasp Pfam domains from ACDB subunit and citrate lyase (ACLA/Y). Discussed eukaryotic clades are indicated. The tree was rooted in the ACL-ATP-grasp subfamily. The MSA was trimmed with trimAl (-gt 0.1) and consisted of 253 positions. **B)** Rooted phylogeny of ACDA subunit indicating the respective eukaryotic clades from ACDB subunit. The tree was rooted in the counterpart most basal clade of ACD(B)_1. The MSA was trimmed with trimAl (-gt 0.20) and consisted of 456 positions. See legend for graphical explanation of selected regions for constructing the MSAs. Pruned clades of the respective groups of interest of **C)** ACDB and **D)** ACDA subunits.

**Figure S30.** Rooted phylogenies and domain architecture of ATP-citrate lyase subunits **A)** alpha (ACLA/B, K15230) and **B)** beta (ACLB, K15231) and the respective region of the fused version of ATP-citrate (pro-S)-lyase (ACLY, K01648). The trees were rooted at the same clade, containing divergent sequences that branch basal in rooted phylogeny of ATP-grasp domain including ACD subfamily (**Figure S29**). The respective MSA consisted of 389 (ACLA) and 602 (ACLB) positions. **C)** Phylogeny of ACLA and ACLB subunits concatenation. Those taxa that only contain one ACLA/B ortholog were included in the concatenation. Yellow highlight indicates an unstable group of Haptista.

**Figure S31. A)** Unrooted view of citrate synthase phylogeny (CS, K01647) indicating the eukaryotic groups discussed in this study (from CS_1 to CS_3). **B-D)** Rooted phylogenies of different CS_1-3 groups. All trees are rooted at the same node which was selected arbitrarily to ease visualization. The final MSA consisted of 370 positions.

**Figure S32. A)** Unrooted view of aconitate hydratase protein family (ACO1, K01681 and ACO2, K17450). The dataset was reduced by removing redundant sequences (identity higher than 80) by taxonomic class. The final MSA consisted of 729 positions. **B)** Refined phylogeny of ACO1 rooted arbitrarily to ease visualization. The final MSA consisted of 650 positions. **C)** Rooted phylogeny ACO2 rooted at ACO1 subfamily.

**Figure S33. A)** Unrooted view of isocitrate dehydrogenase protein family phylogeny (IDH1-2, K00031 and IDH3, K00030) indicating the eukaryotic groups discussed in this study. Branch colors indicate gene family (upper panel) and taxonomy levels (lower panel). **B-F)** Rooted phylogenies of different IDH groups. **B)** and **E)** are refined phylogenies of the respective subfamilies and final MSAs consisted of 408 and 302 positions, respectively. Trees were rooted arbitrarily to ease visualization.

**Figure S34.** Unrooted (upper panel) and rooted (lower panel) phylogenies of **A)** 2-oxoglutarate dehydrogenase E1 component (SucA, K00164), and **B)** 2-oxoglutarate dehydrogenase E2 component (dihydrolipoamide succinyltransferase, SucB, K00658). The trees were arbitrarily rooted to ease visualization (arrow). The final MSAs consisted of 905 and 352 positions, respectively.

**Figure S35.** Unrooted view of succinyl-CoA synthetase **A)** alpha and **B)** beta subunit phylogenies , and respective rooted views **C)** and **D)**. The trees were arbitrarily rooted to ease visualization (arrow). The final MSAs consisted of 316 and 411 positions respectively.

**Figure S36.** Unrooted (upper panel) and respective pruned (lower panel) phylogenies of **A)** succinate dehydrogenase (ubiquinone) flavoprotein subunit (SDH1, K00234), **B)** succinate dehydrogenase (ubiquinone) iron-sulfur subunit (SDH2, K00235), **C)** succinate dehydrogenase (ubiquinone) cytochrome b560 subunit together with succinate dehydrogenase (ubiquinone) membrane anchor subunit (SDH3, K00236 and SHD4, K00237). The final MSAs consisted of 599, 253 and 107 positions, respectively. Inset tree in **C)** shows the symmetric distributions of SDH3 and SDH4 subunits, in which arcs connect sequences from the same organisms and discontinuous line indicates the potential duplication node of both subunits.

**Figure S37. A)** Refined phylogeny of fumarate hydratase, class I (FUMA/B, K01676). Inset tree indicates unstable topology of Kinetoplastid sequences branching within and outside the LECA clade when including or excluding a divergent outgroup. The final MSA consisted of 537 positions. **B)** Unrooted (upper panel) and rooted (lower panel) view of fumarate hydratase, class II, phylogeny (FUMC, K01679). The rooted tree (below) is a refined phylogeny including closely related sequences to the LECA clade, in which Metamonada and other sequences branch with eukaryotic sequences. The tree was rooted at the most basal bacterial branch in the unrooted tree. The final MSA consisted of 461 positions.

**Figure S38. A)** Unrooted and **B)** rooted phylogeny of lactate/malate dehydrogenase protein family. The tree includes lactate dehydrogenases (LDH, K00016), and malate dehydrogenases (MDH, K00024, MDH1, K00025 and MDH2, K00026). The tree was rooted in the LDH protein family (arrow). The final MSA consisted of 311 positions. Conservation of introns along LDH/MDH MSA using a selected set of eukaryotic genomes (*Eukarya4 (Vosseberg et al)*, see methods) mapped onto **C)** LDH and **D)** MDH phylogenies including relative prokaryotic sequences. Column labels denote the position in the MSA and the respective intron phases identified. Bars at the right show the intron density for each coding gene.

**Figure S39. A)** Unrooted and **B)** rooted view of isocitrate lyase phylogeny (ACEA, K01637). The tree was rooted at midpoint and the final MSA consisted of 419 positions.

**Figure S40.** Extended view of phylogenetic profile for the proposed origins of selected orthogroups mapped onto the eukaryotic tree of life. Bold labels of columns indicate those that are proposed to be present in LECA. Different shapes of cells indicate presence of one or diverse sequences with the respective targeting signal obtained with TargetP (*see legend*). This is an extended view of Figure 3 in the main text, see figure caption for additional information.

**Figure S41.** Clustermap representing the correlations of the orthogroup distribution of CCM enzymes. The correlation was inferred using the phi coefficient (*matthews_corrcoef* function in python) and converting the matrix to 0 (absence) and 1 (presence) values. Note that the distribution of characteristic traits were included in the analysis (Anaerobic, Primary and Secondary endosymbionts). Red and blue cells indicate correlated and anticorrelated distributions respectively. The raw data for this analysis is provided in **DATA S6**.

**Figure S42.** Distribution of parallel glycolysis in the cytoplasm and in the **A)** chloroplast or **B)** mitochondria using all proteomes of EukProt v3. Colored cells indicate organisms that have one sequence with no targeting signal (cytoplasm) and another with the respective organelle targeting. Only those organisms with more than 3 enzymes with parallel targeting are shown.

**Figure S43.** Origins and evolution of cytoplasmic and chloroplast glycolysis and PPP mapped onto an schematic representation of the eukaryotic tree of life (zoomed in the Archaeplastida clade). The boxes encompass enzymes predicted to be present in the corresponding ancestral nodes. Tree reconciliations focus on major events, representing an approximation inferred manually. For instance, duplications within the Archaeplastida ancestor might represent independent duplications in distinct lineages. The prefix symbol minus (-) denotes potential gene loss.

**Figure S44.** Distribution of targeted glycolytic enzymes using all proteomes of EukProt v3. **A)** Quantification of targeted enzymes by eukaryotic group. Left panel shows the number of sequences with/without targeting signal and the right panel shows the number of eukaryotic sequences with a combination of targeting. Plots within the blue box represent those enzymes using substrates with more than three carbons while the purple box include those enzymes using substrates of three carbons (triose). Phylogeny of **B)** GAPDH, **C)** PGK, and **D)** ENO mapping the respective distribution of mitochondrial and chloroplast targeting. Branches are colored according to the eukaryotic groups, and peripheral prokaryotic groups are collapsed.

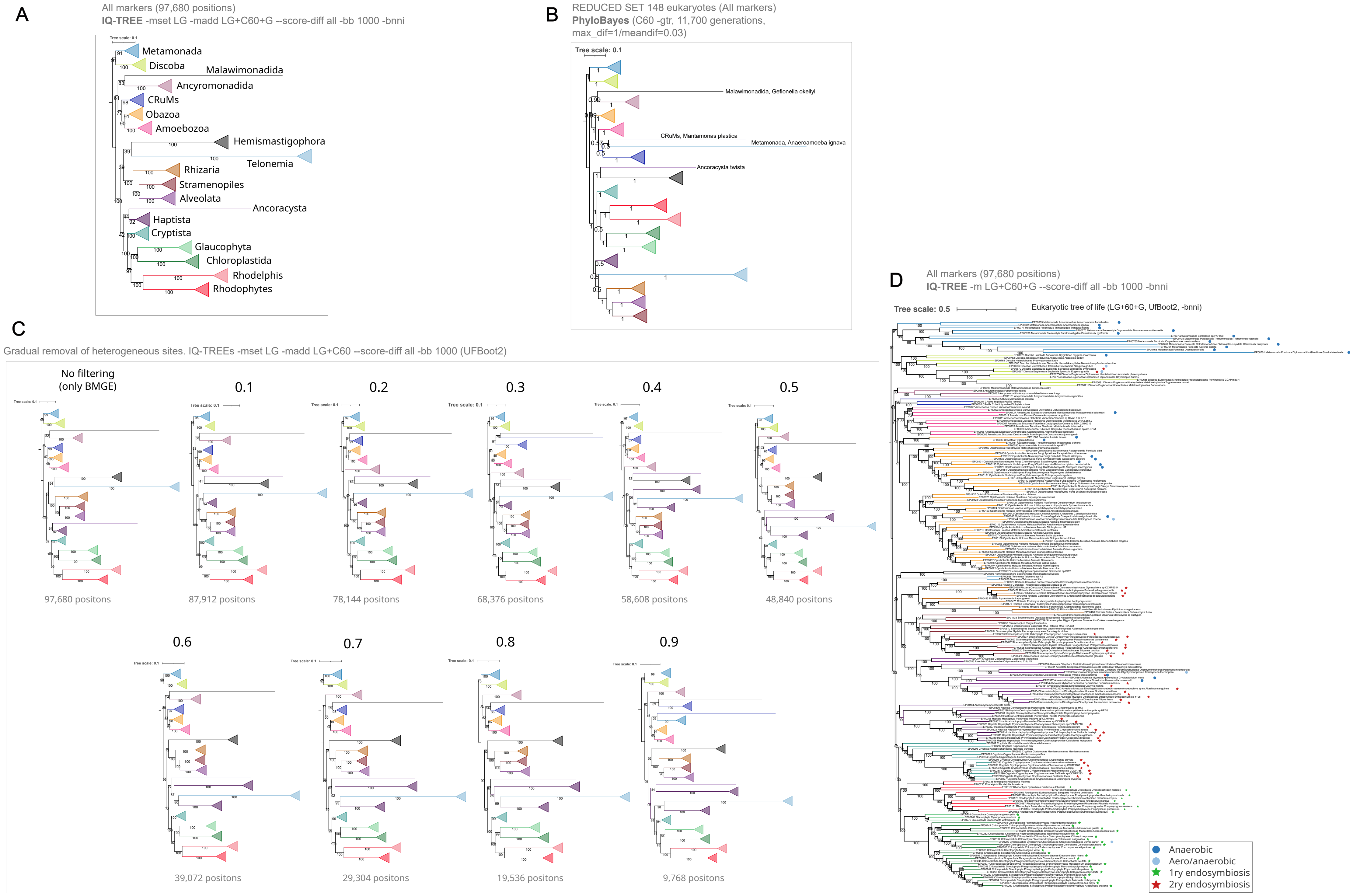

**Supplementary Figure 1.**

### General workflow for phylogenetic reconstructions of CCM enzymes

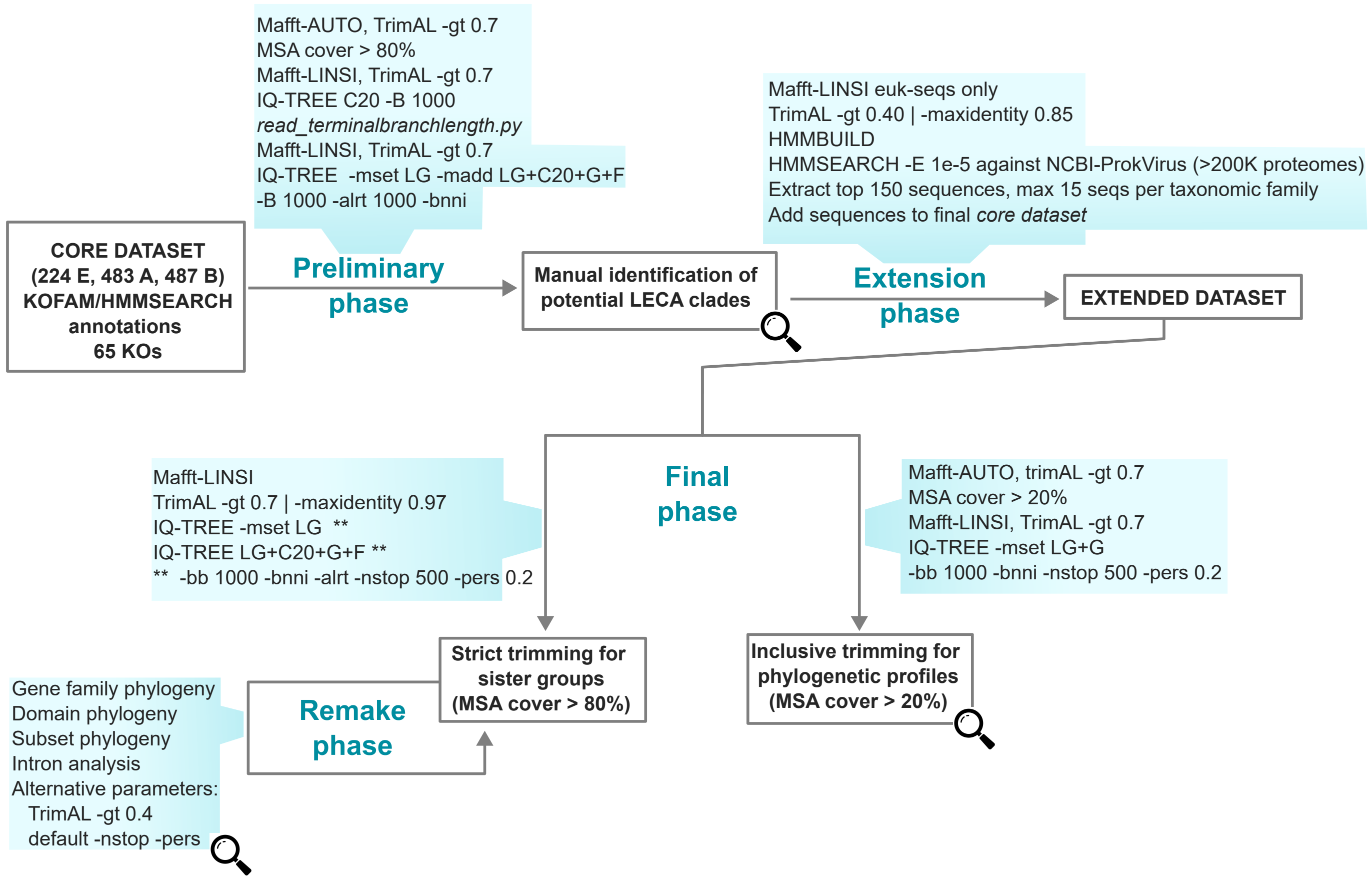

**Supplementary Figure 2.**

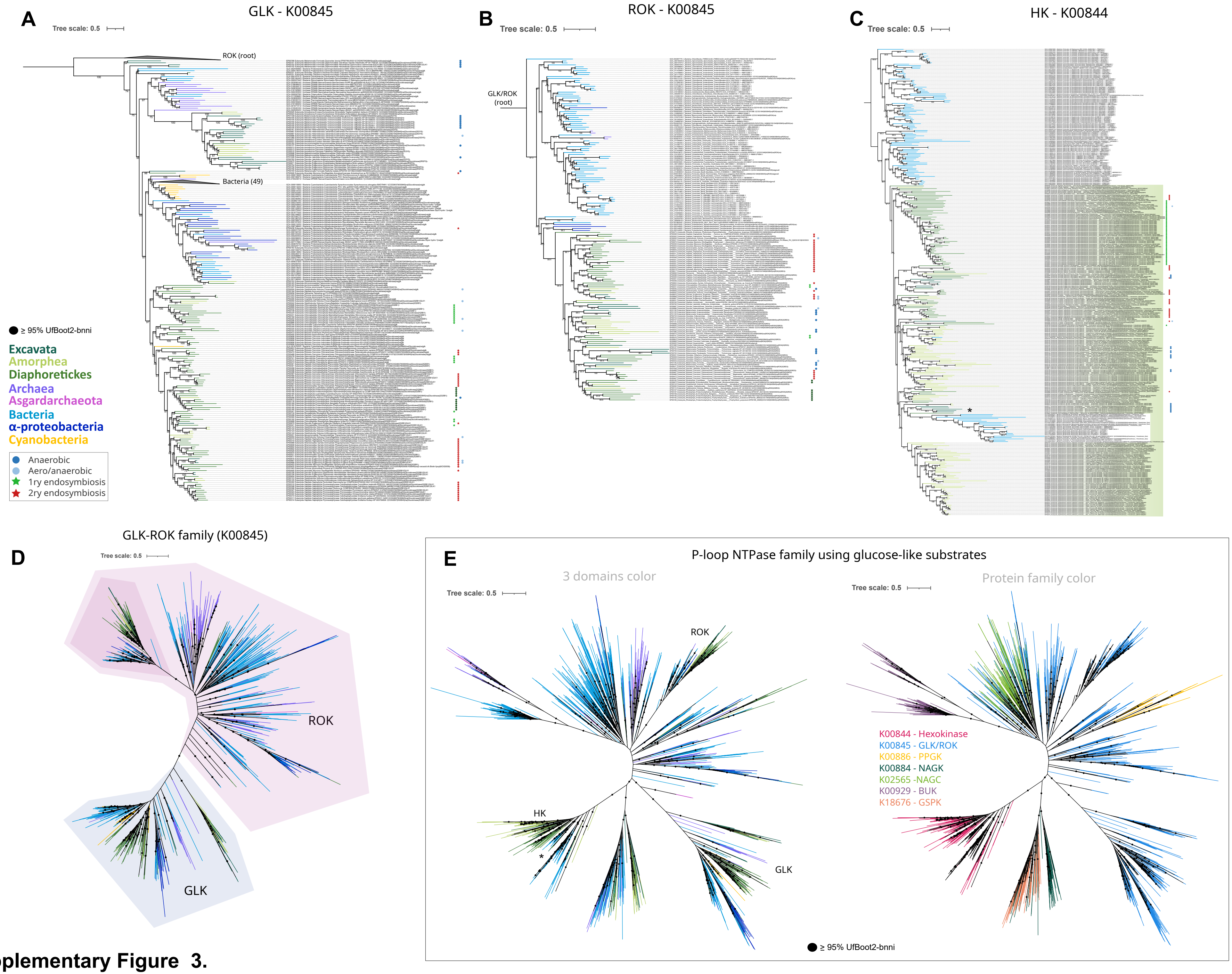

**Supplementary Figure 3.**

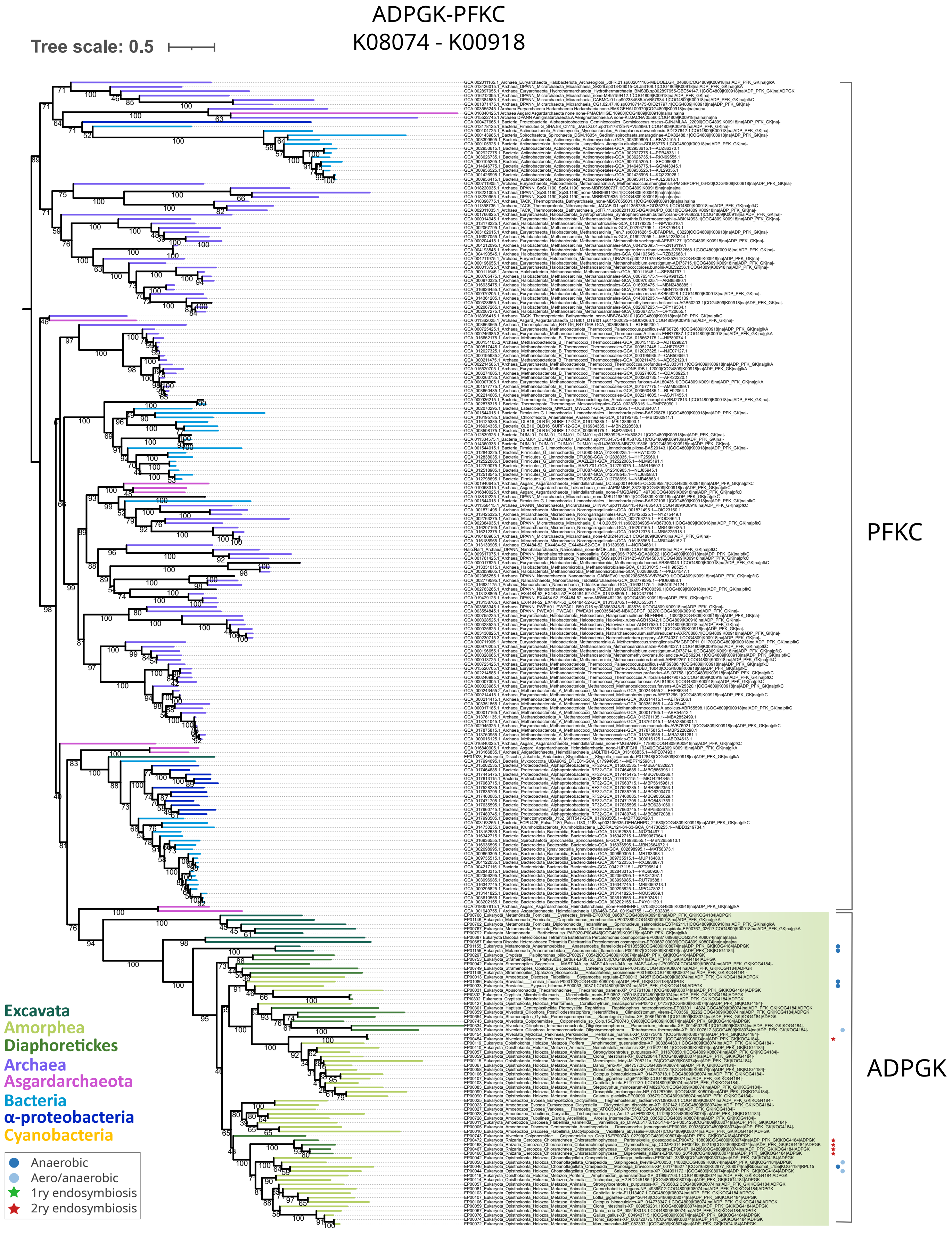

**Supplementary Figure 4.**

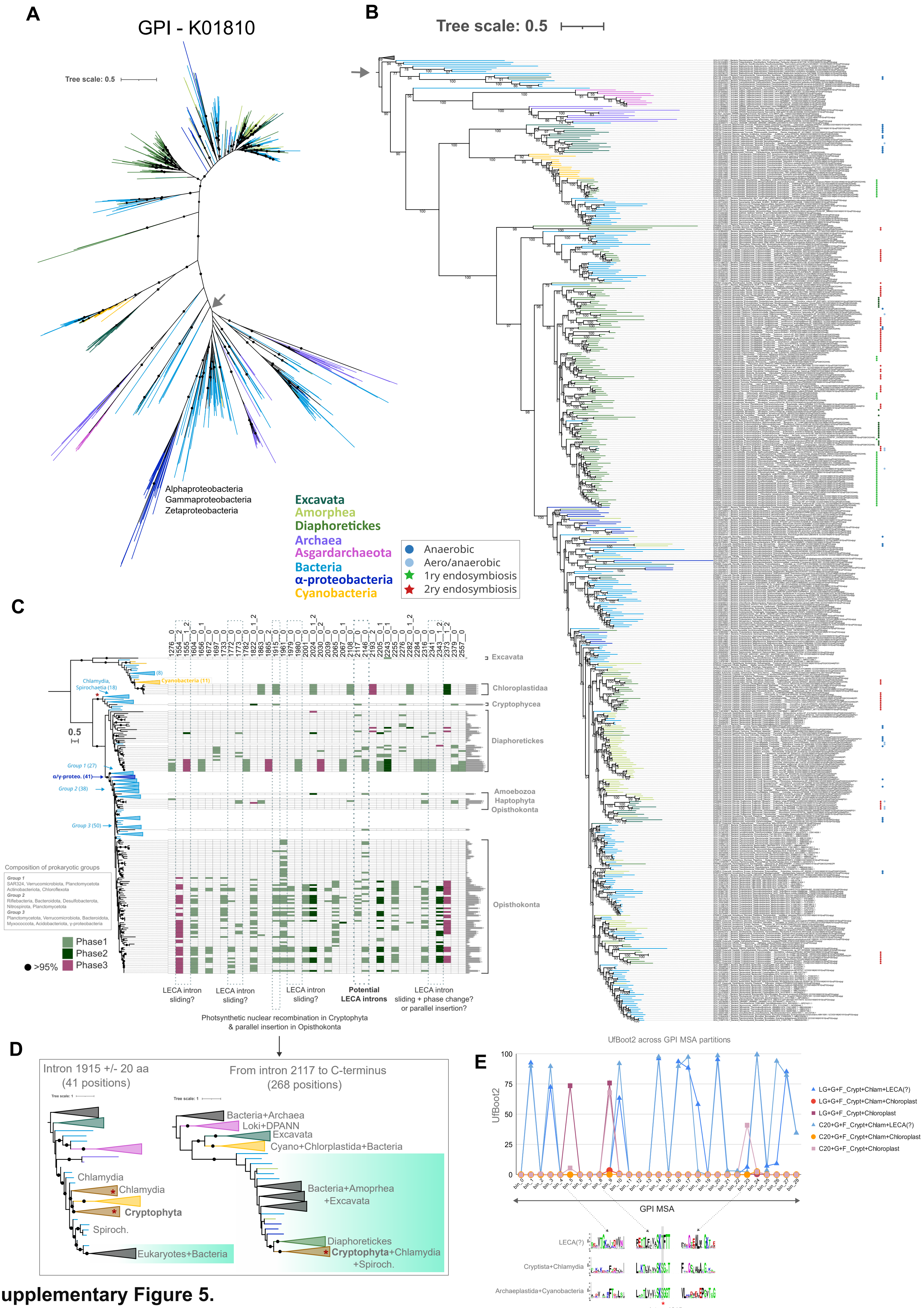

**Supplementary Figure 5.**

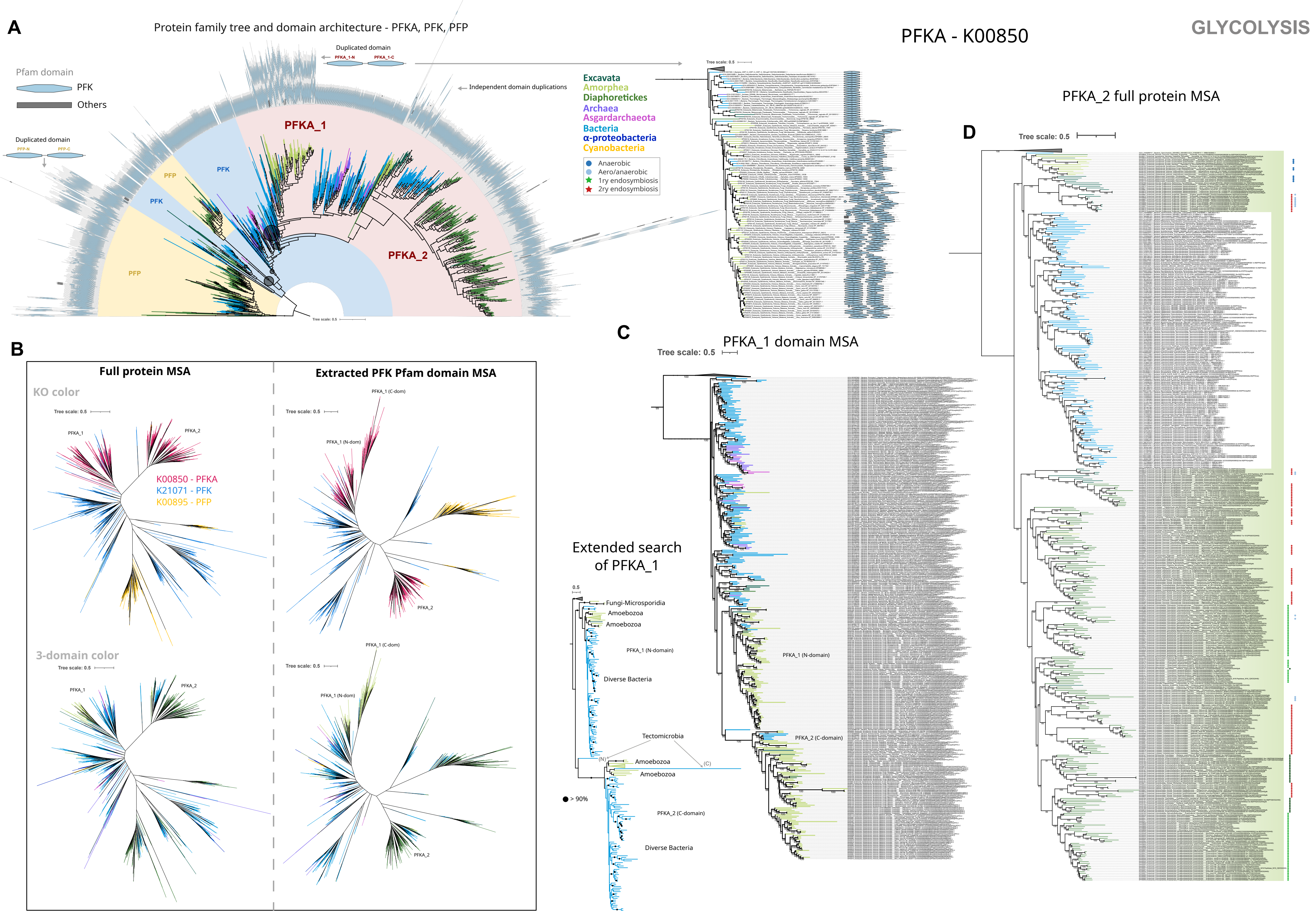

**Supplementary Figure 6.**

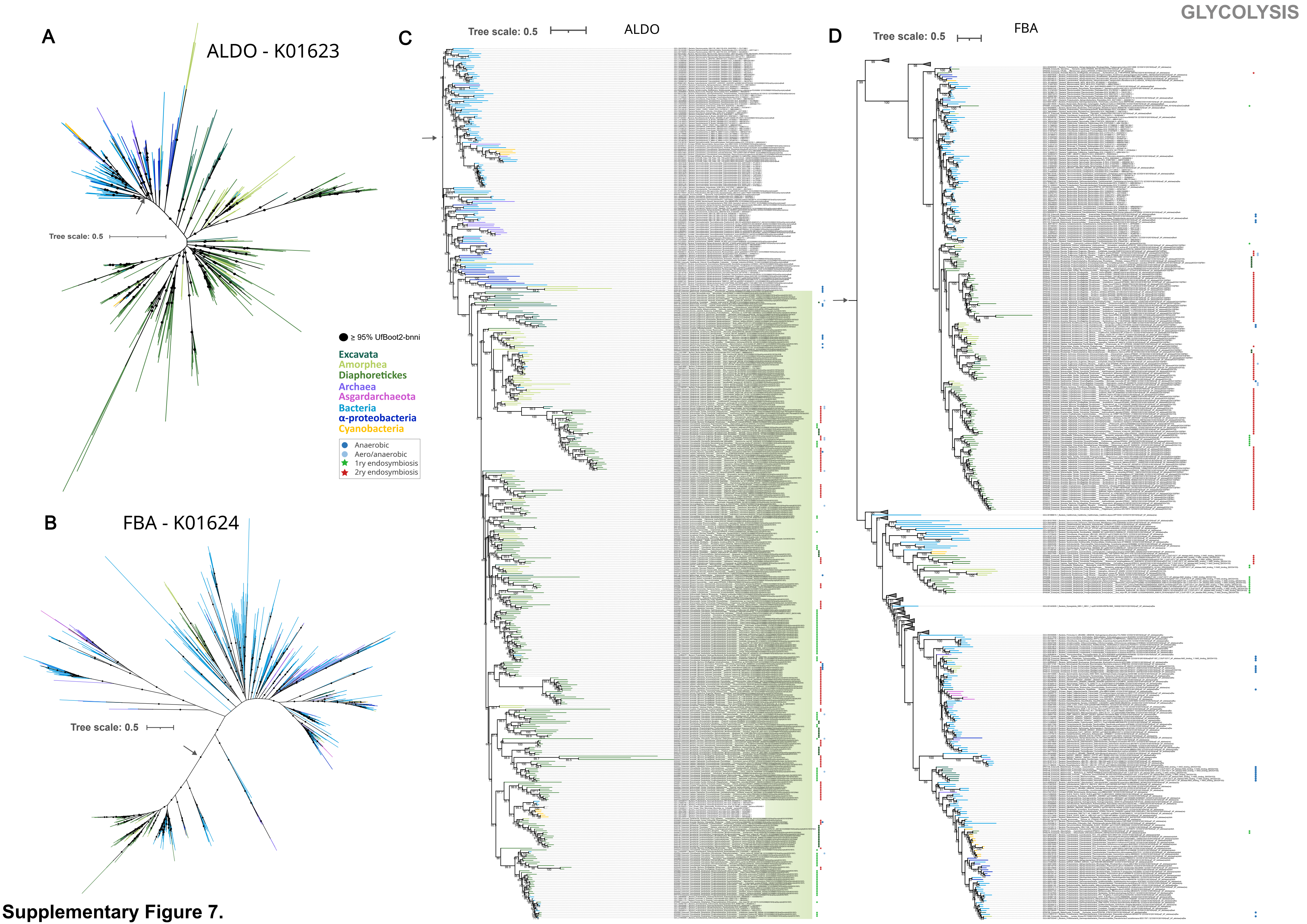

**Supplementary Figure 7.**

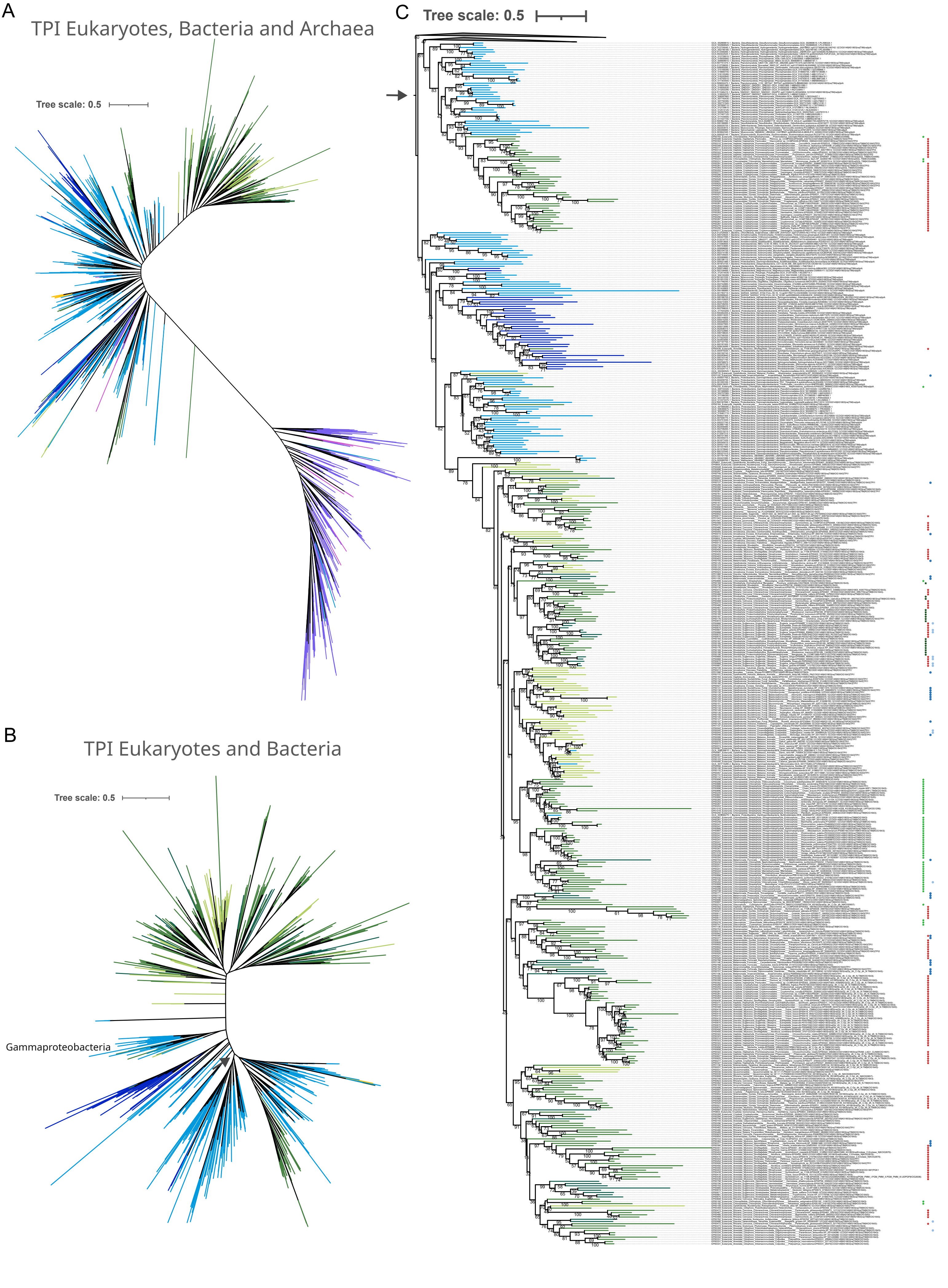

**Supplementary Figure 6.**

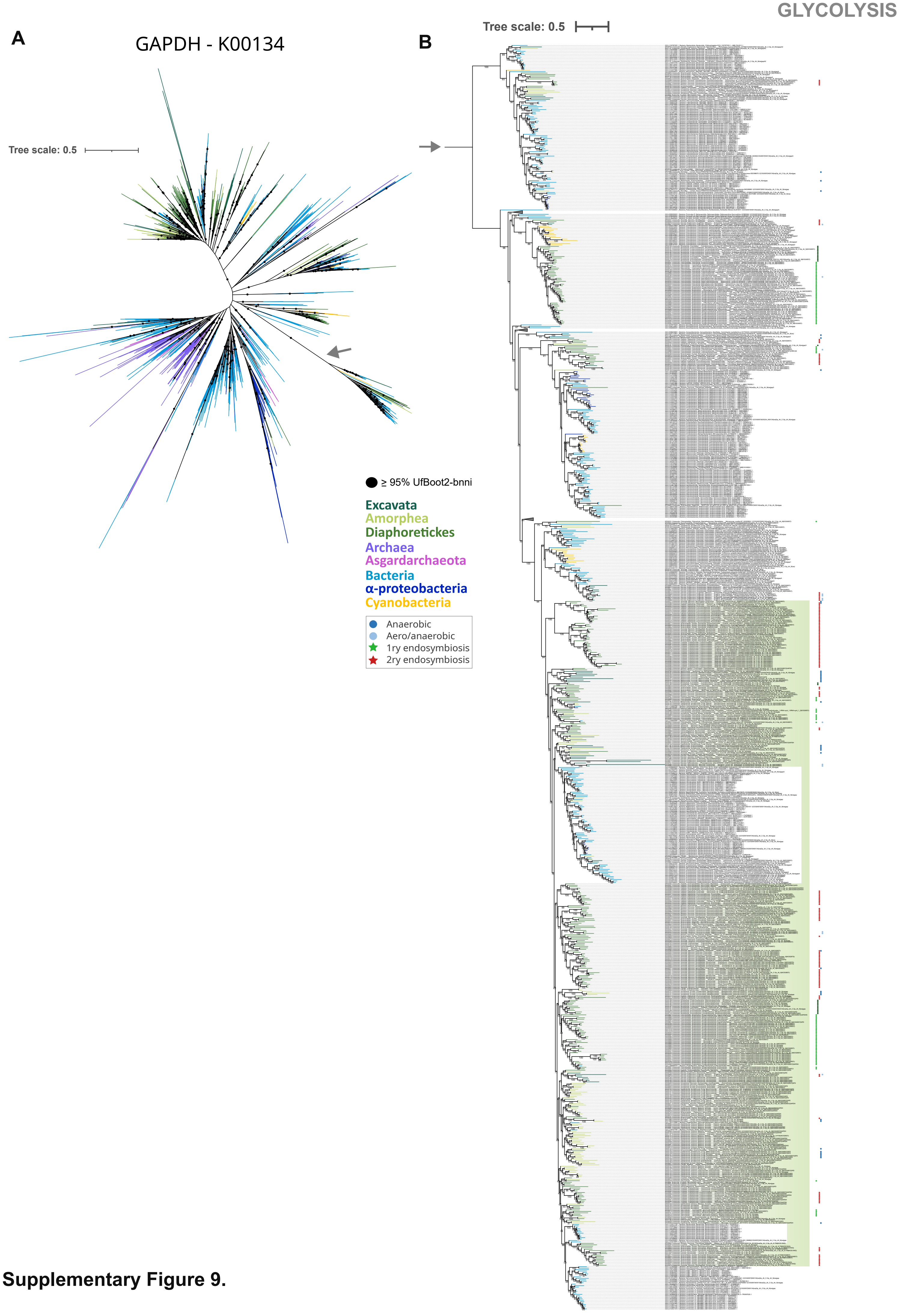

**Supplementary Figure 9.**

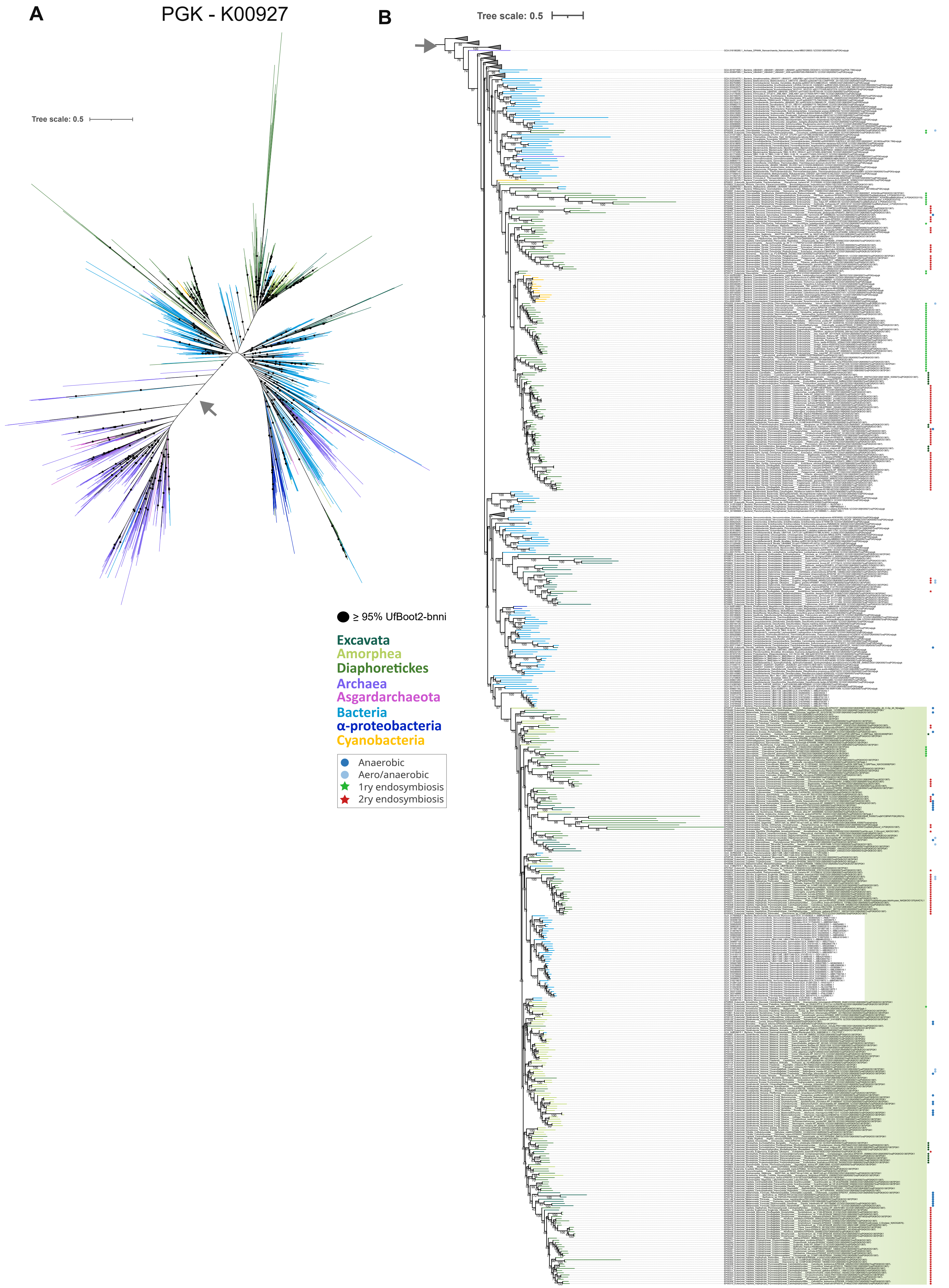

**Supplementary Figure 10.**

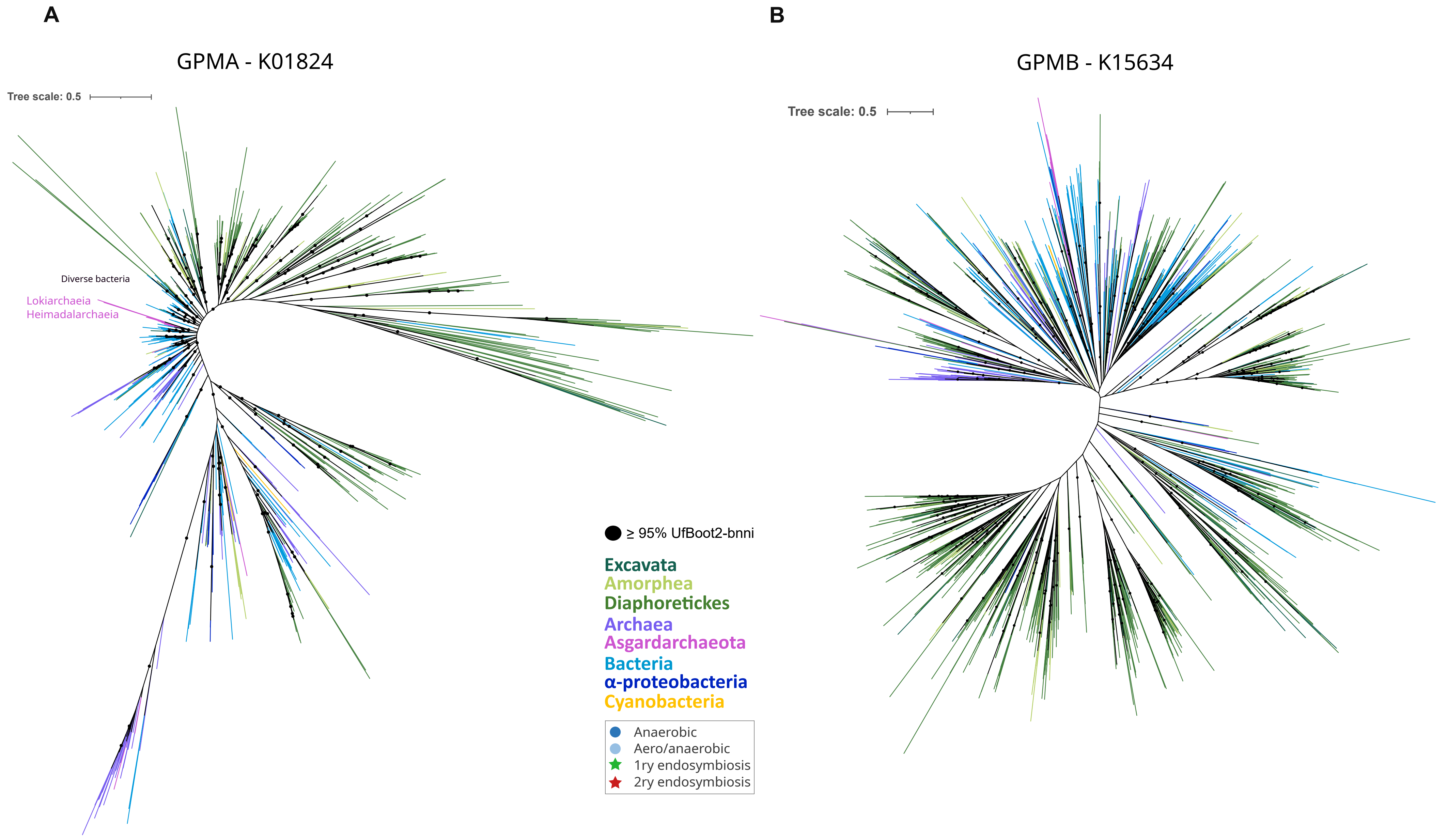

**Supplementary Figure 11.**

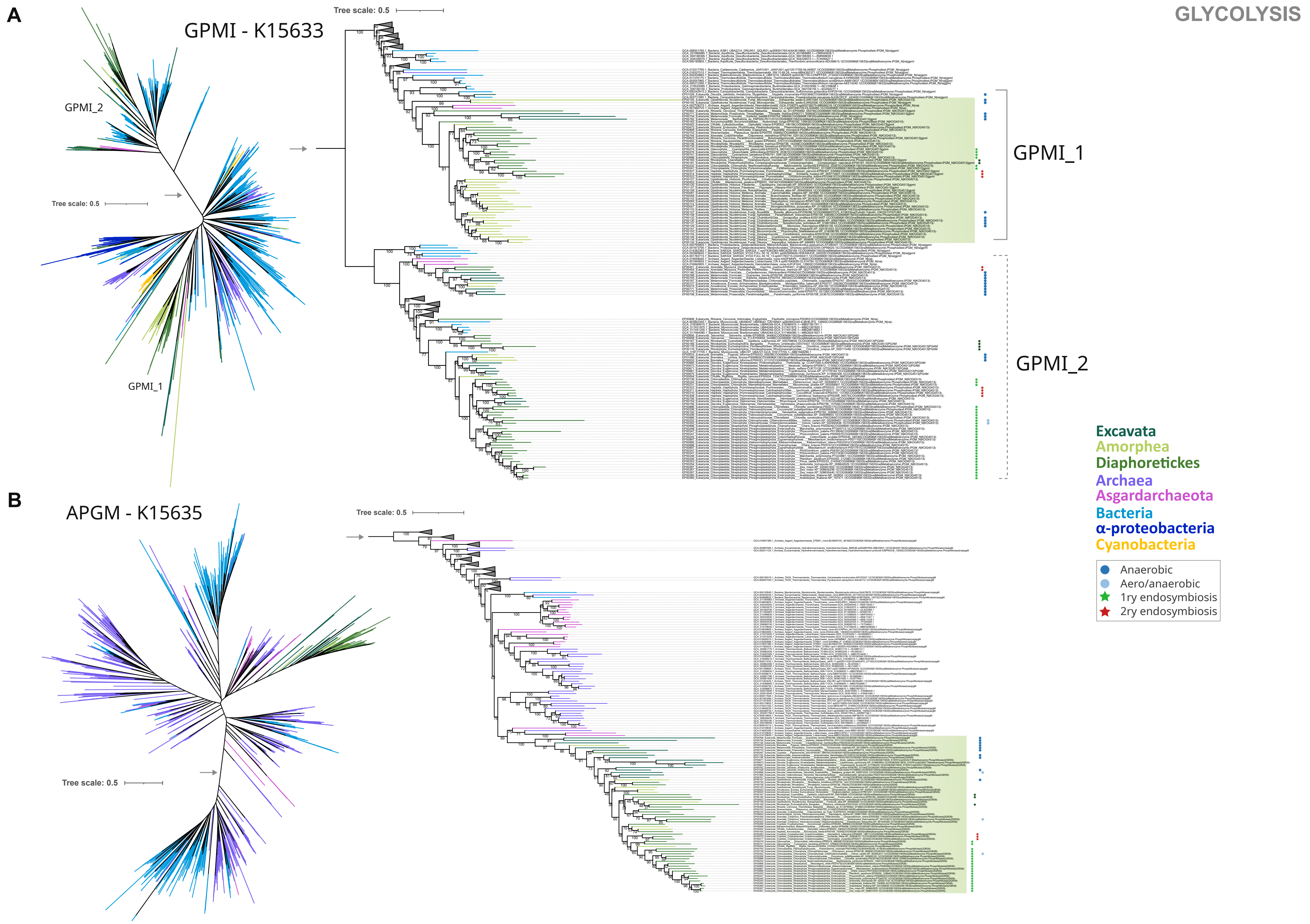

**Supplementary Figure 12.**

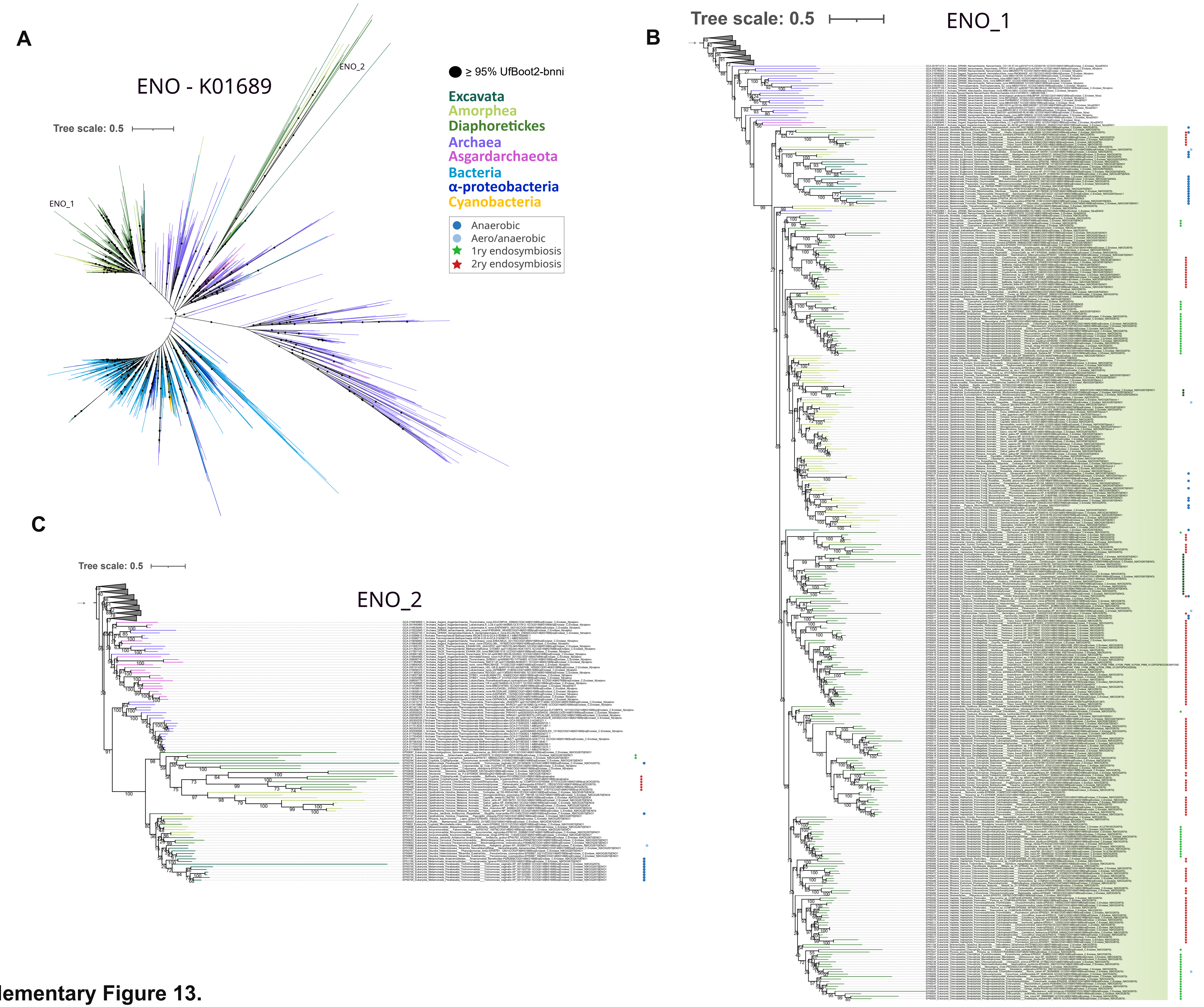

**Supplementary Figure 13.**

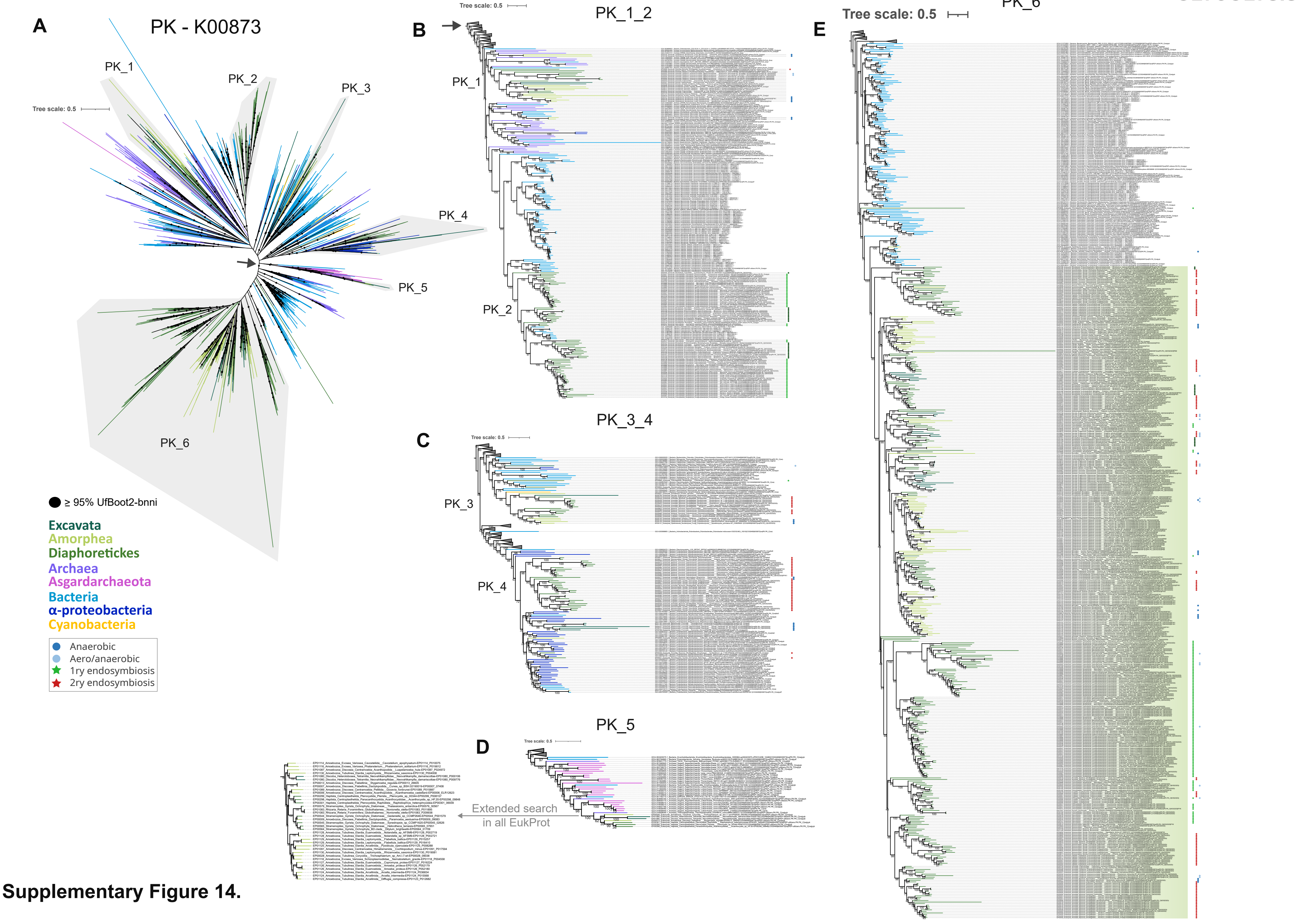

**Supplementary Figure 14.**

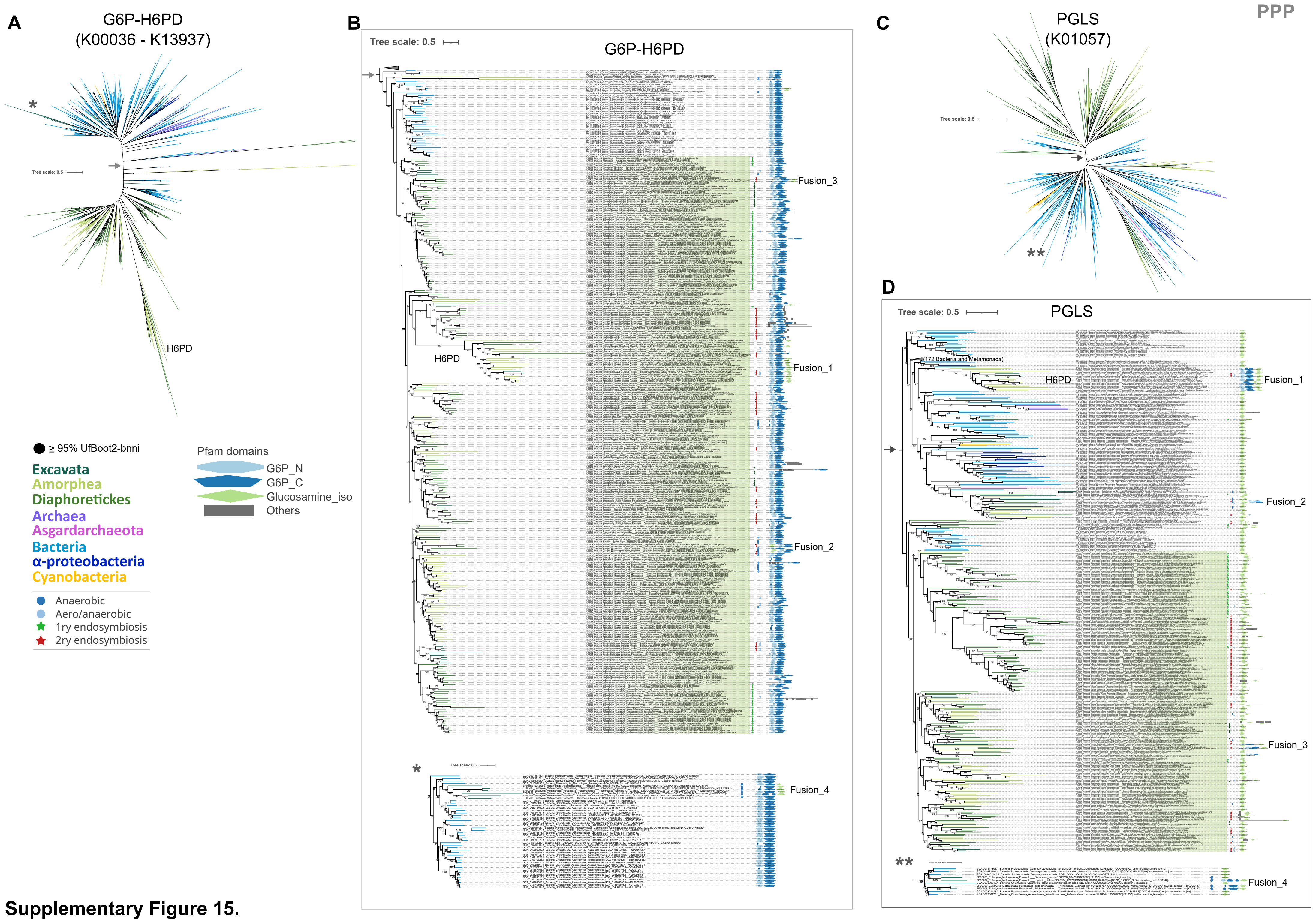

**Supplementary Figure 15.**

### AZF - K19243

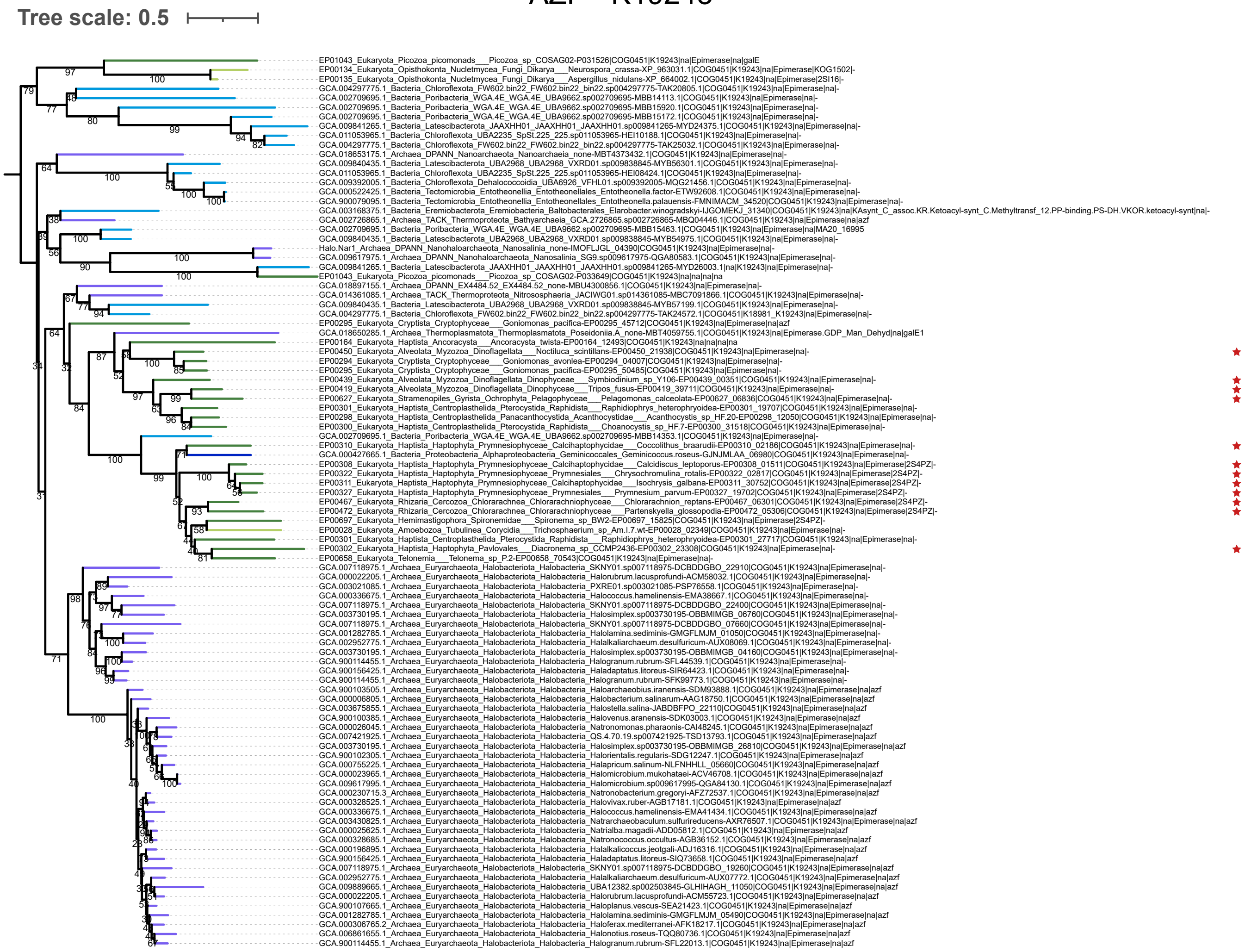

**Supplementary Figure 16.**

A

PGL - K07404

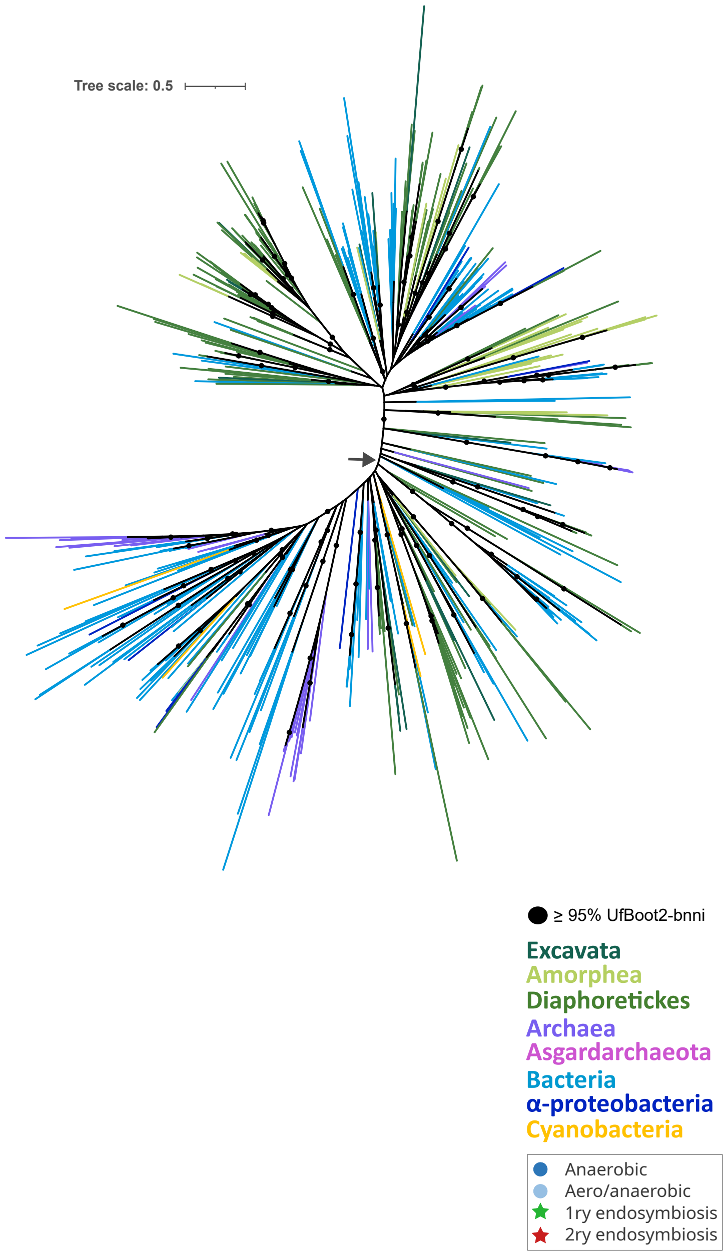

B

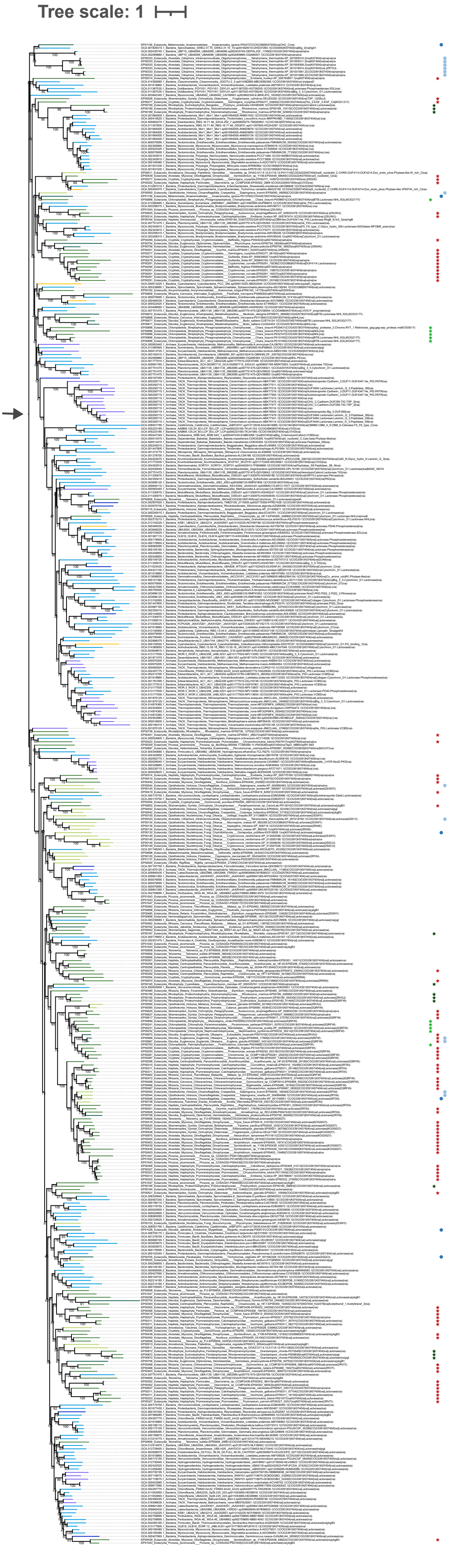

**Supplementary Figure 17.**

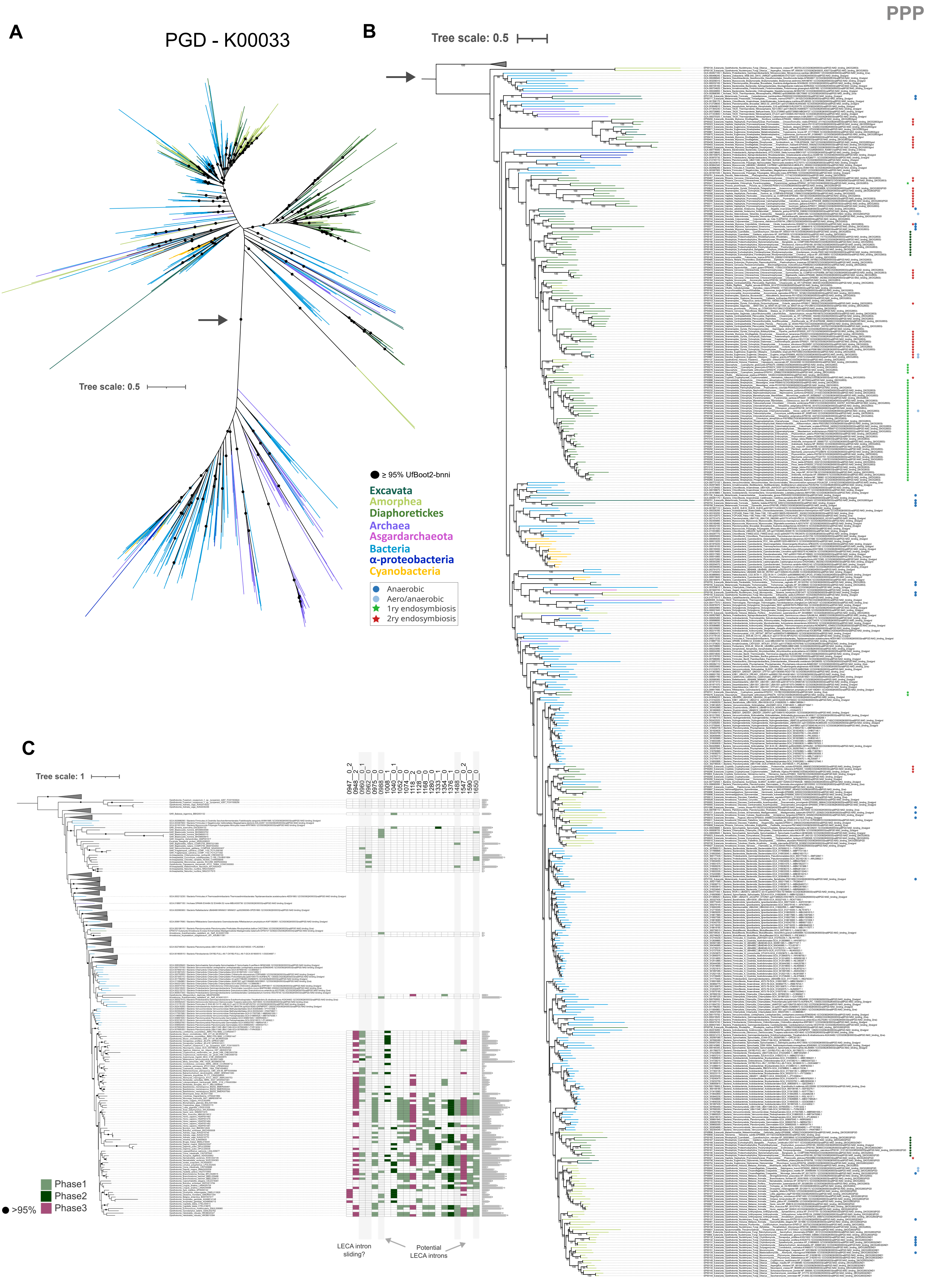

**Supplementary Figure 18.**

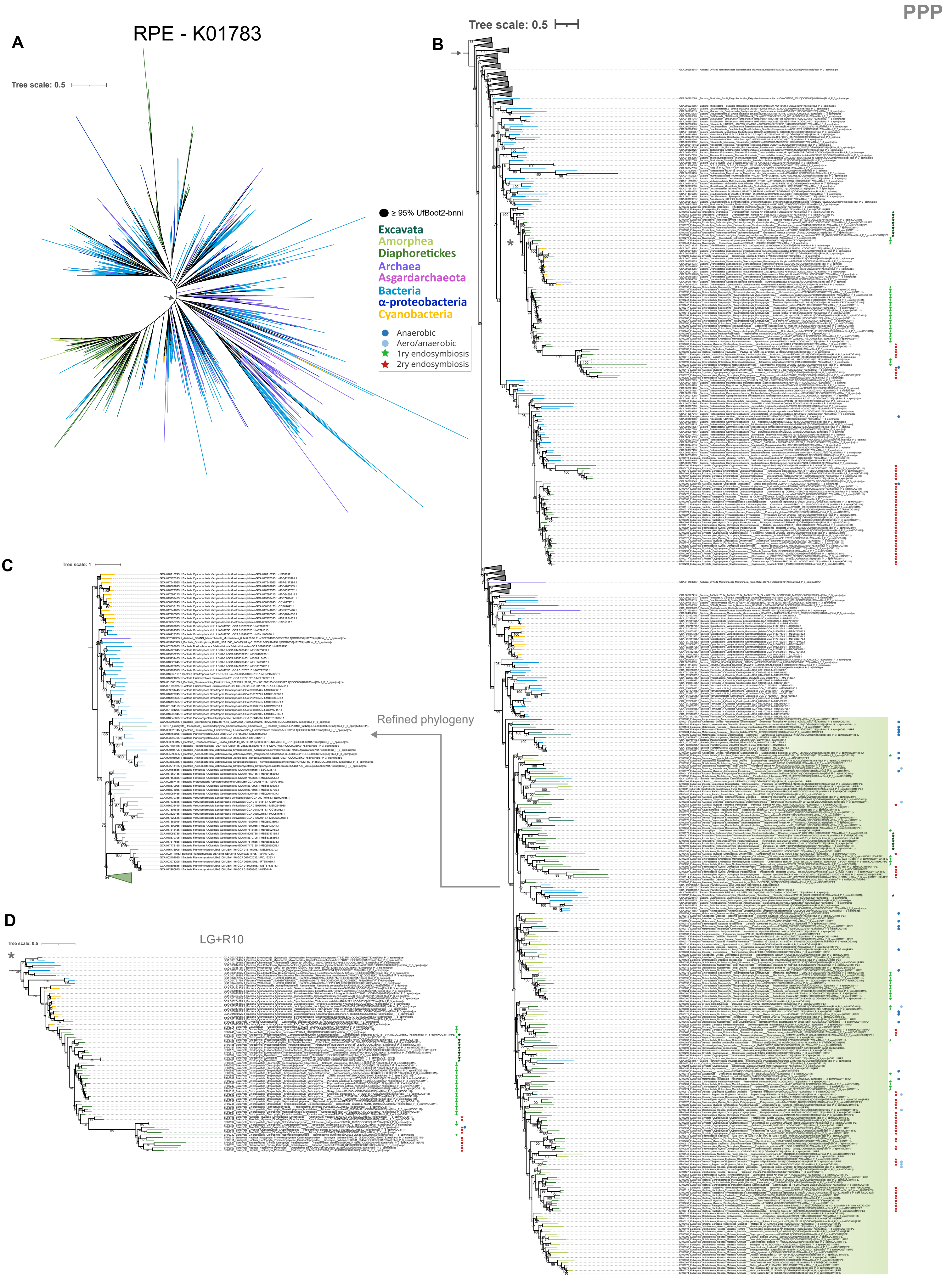

**Supplementary Figure 19.**

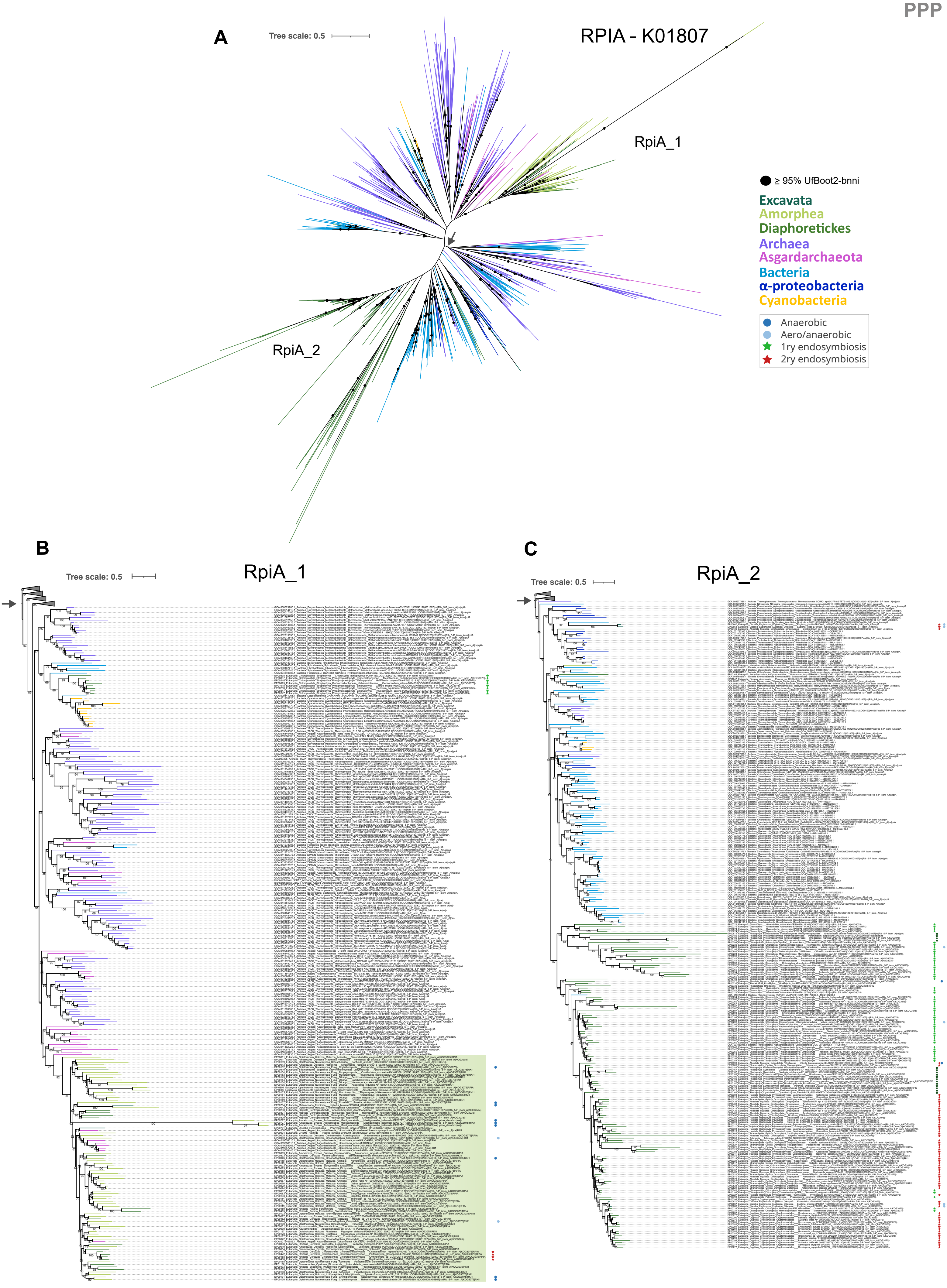

**Supplementary Figure 20.**

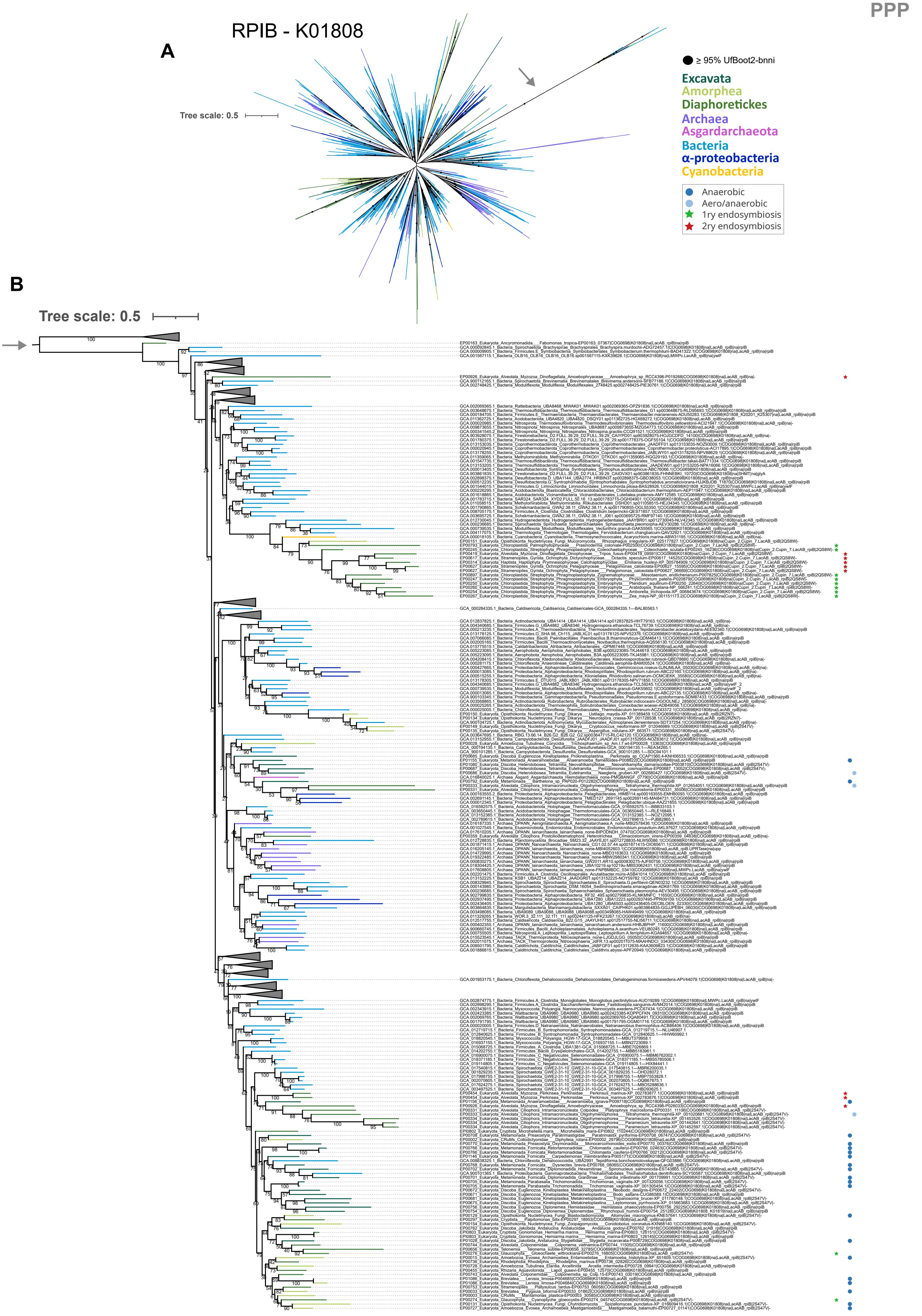

**Supplementary Figure 21.**

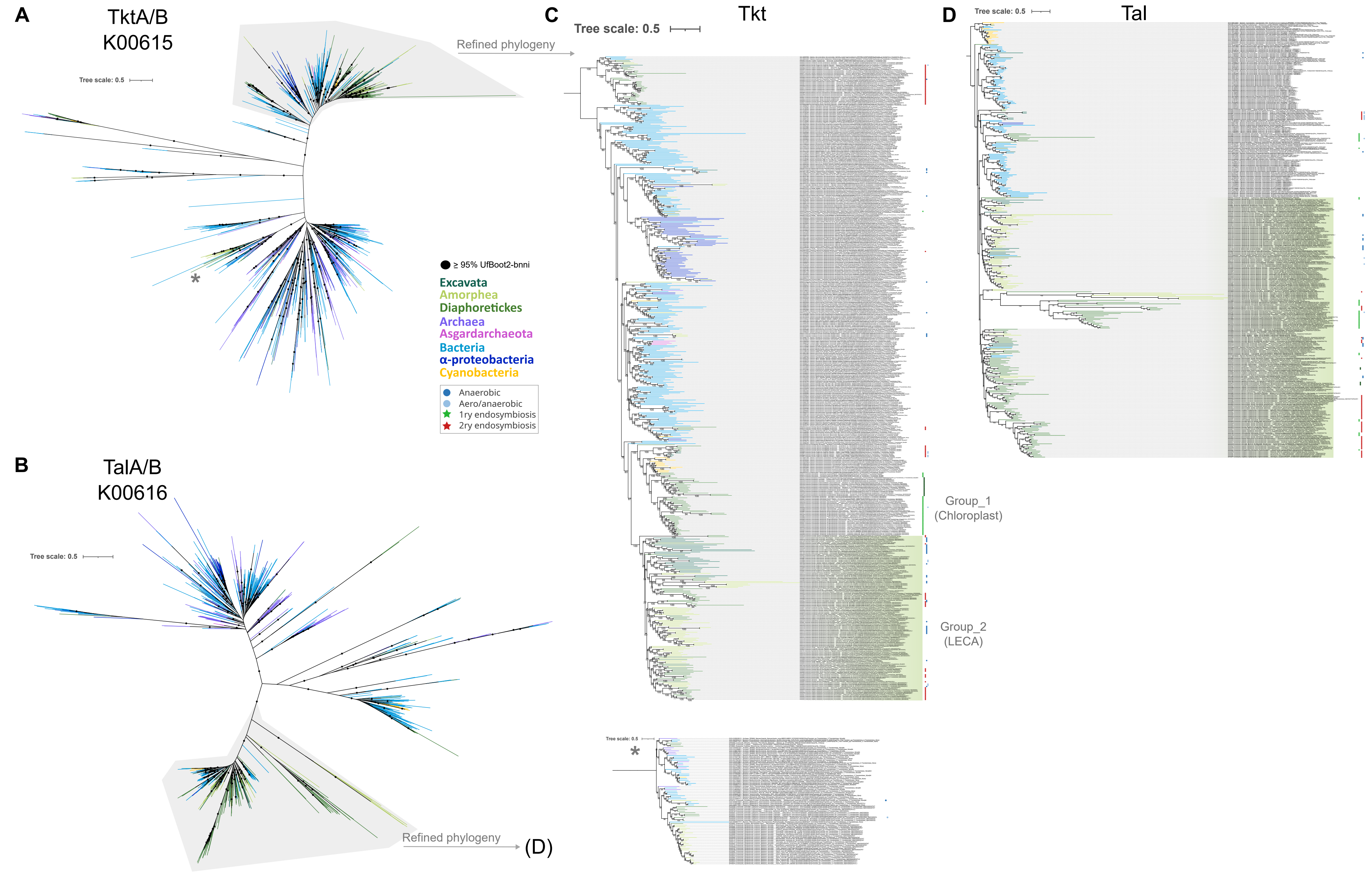

**Supplementary Figure 22.**

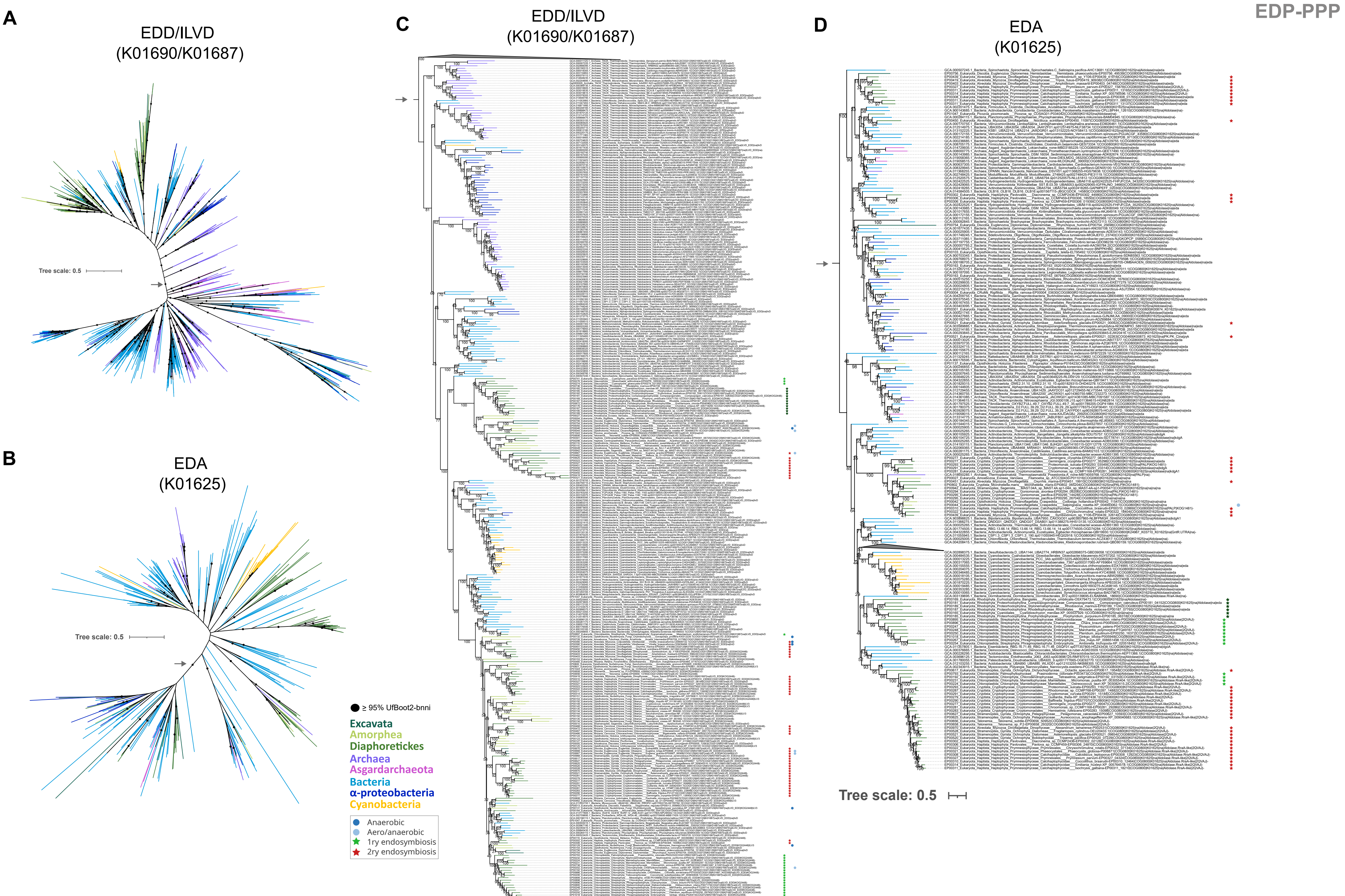

**Supplementary Figure 23.**

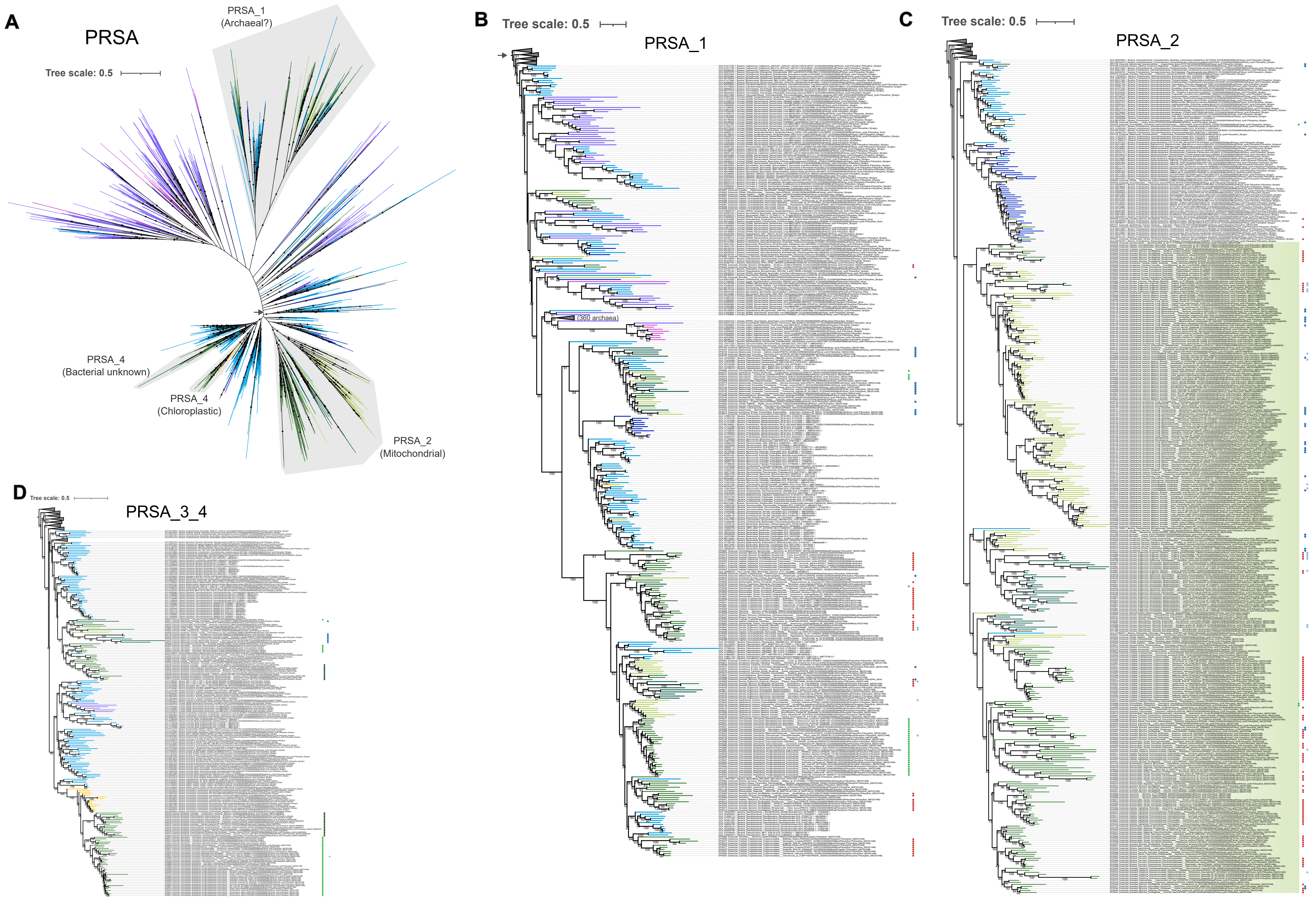

**Supplementary Figure 24.**

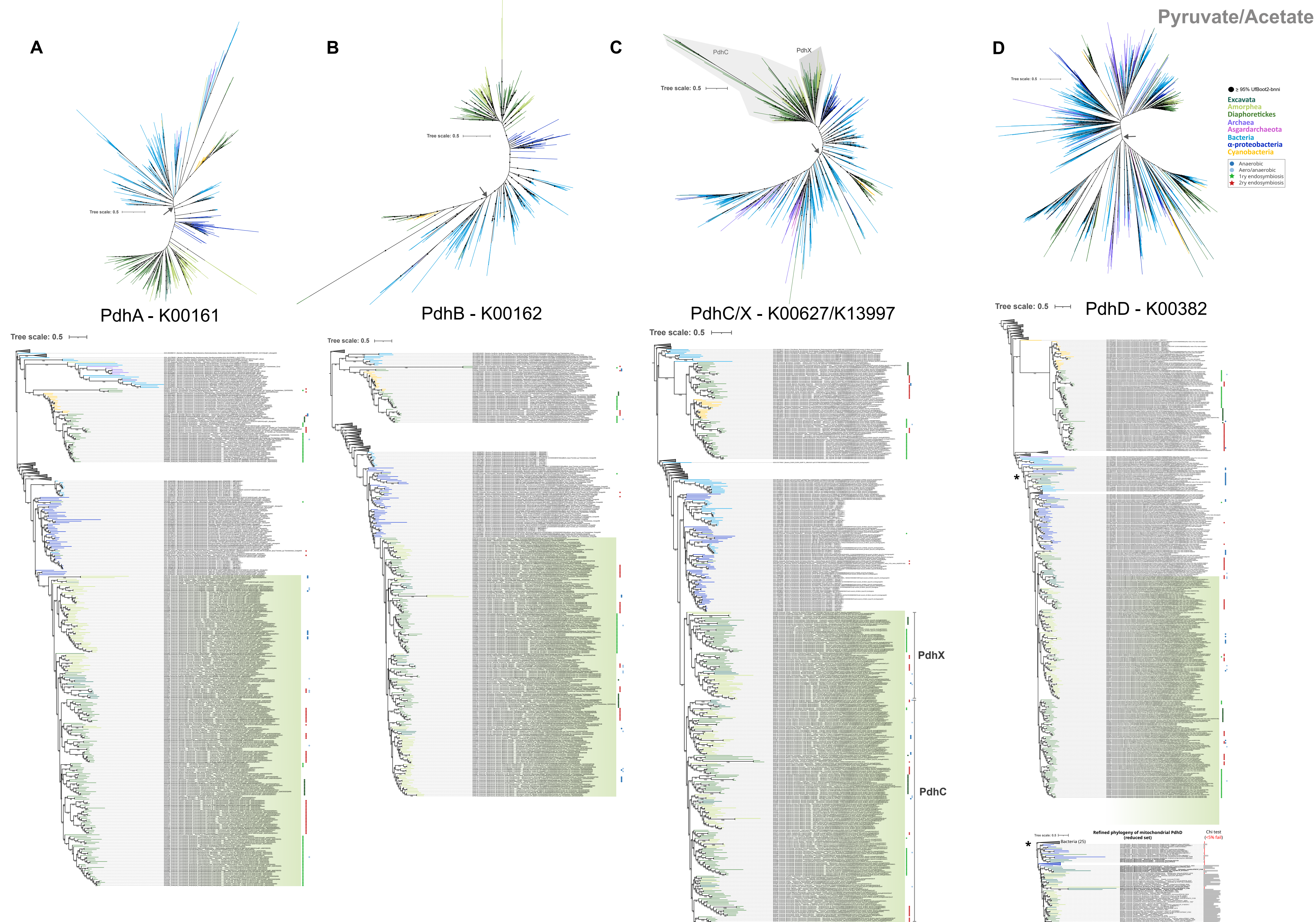

**Supplementary Figure 25.**

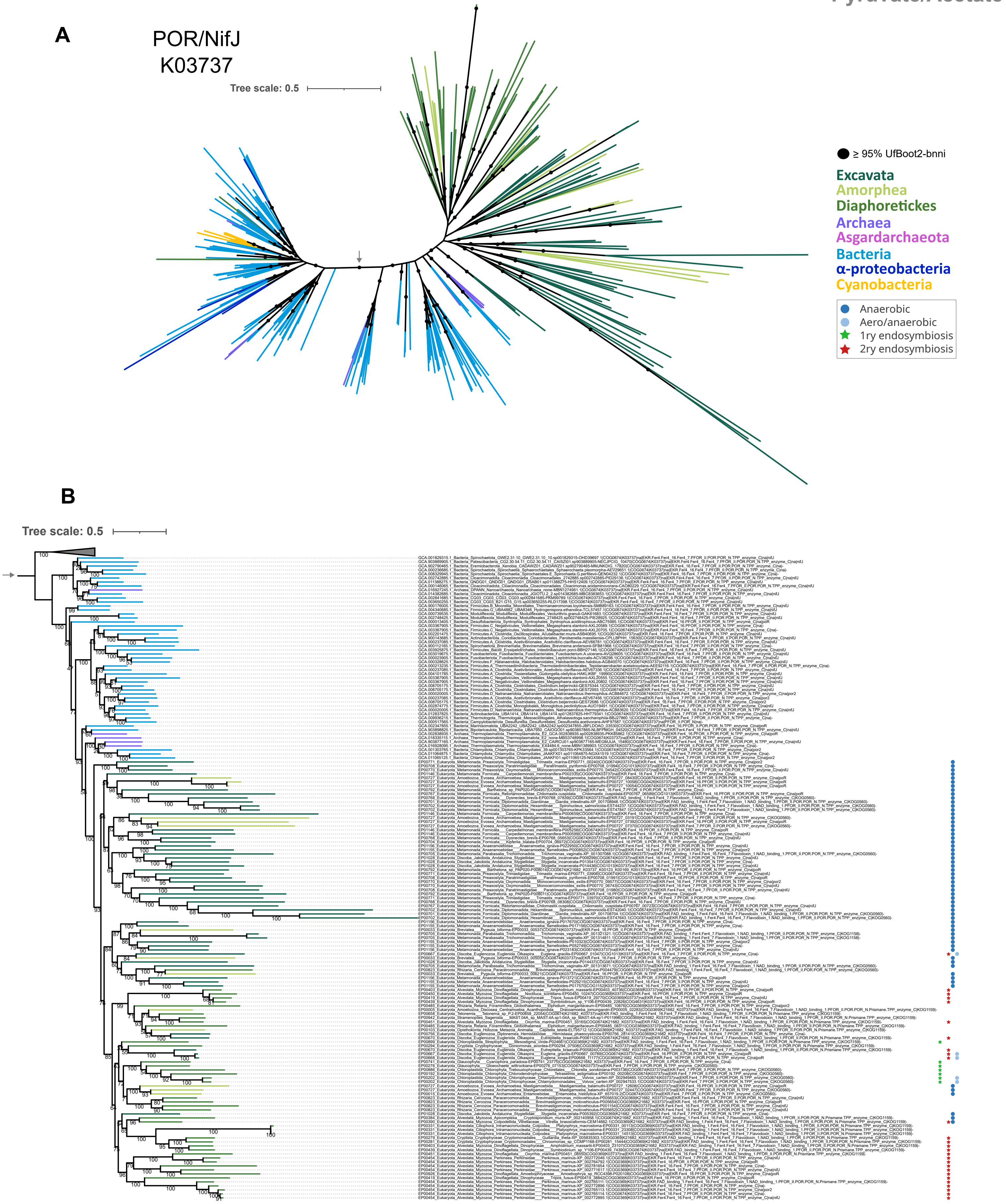

Supplementary Figure 26.

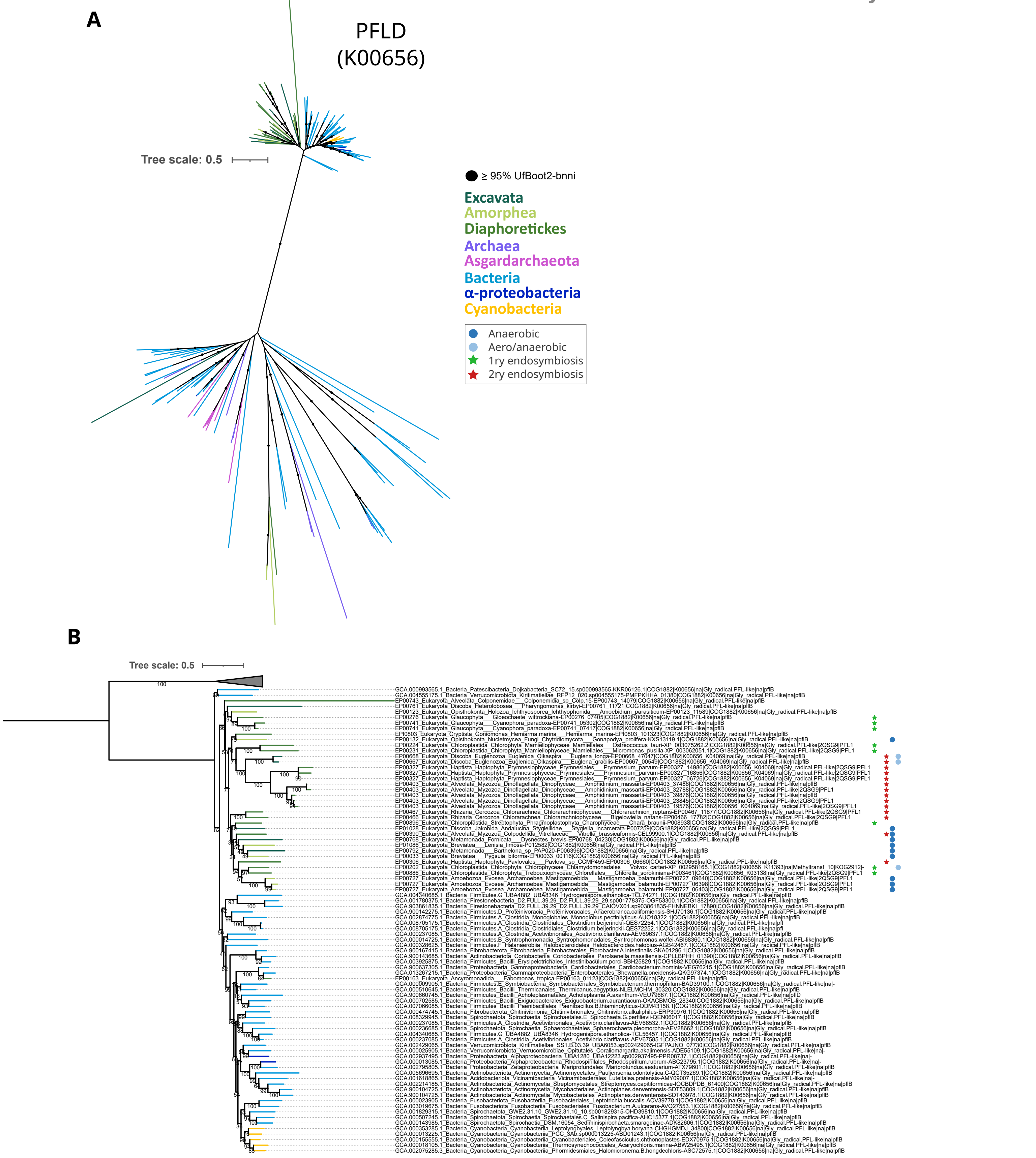

**Supplementary Figure 27.**

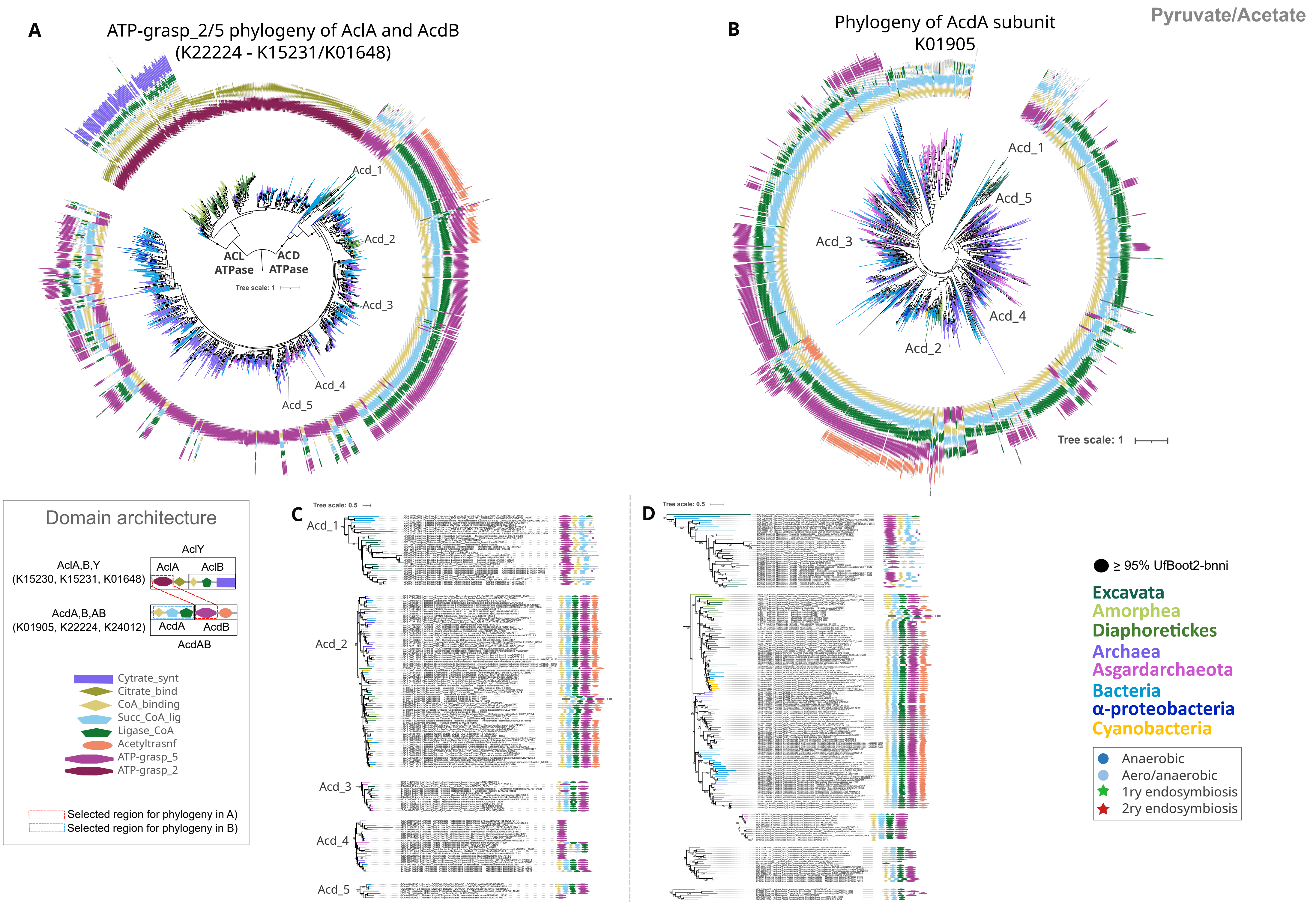

**Supplementary Figure 29.**

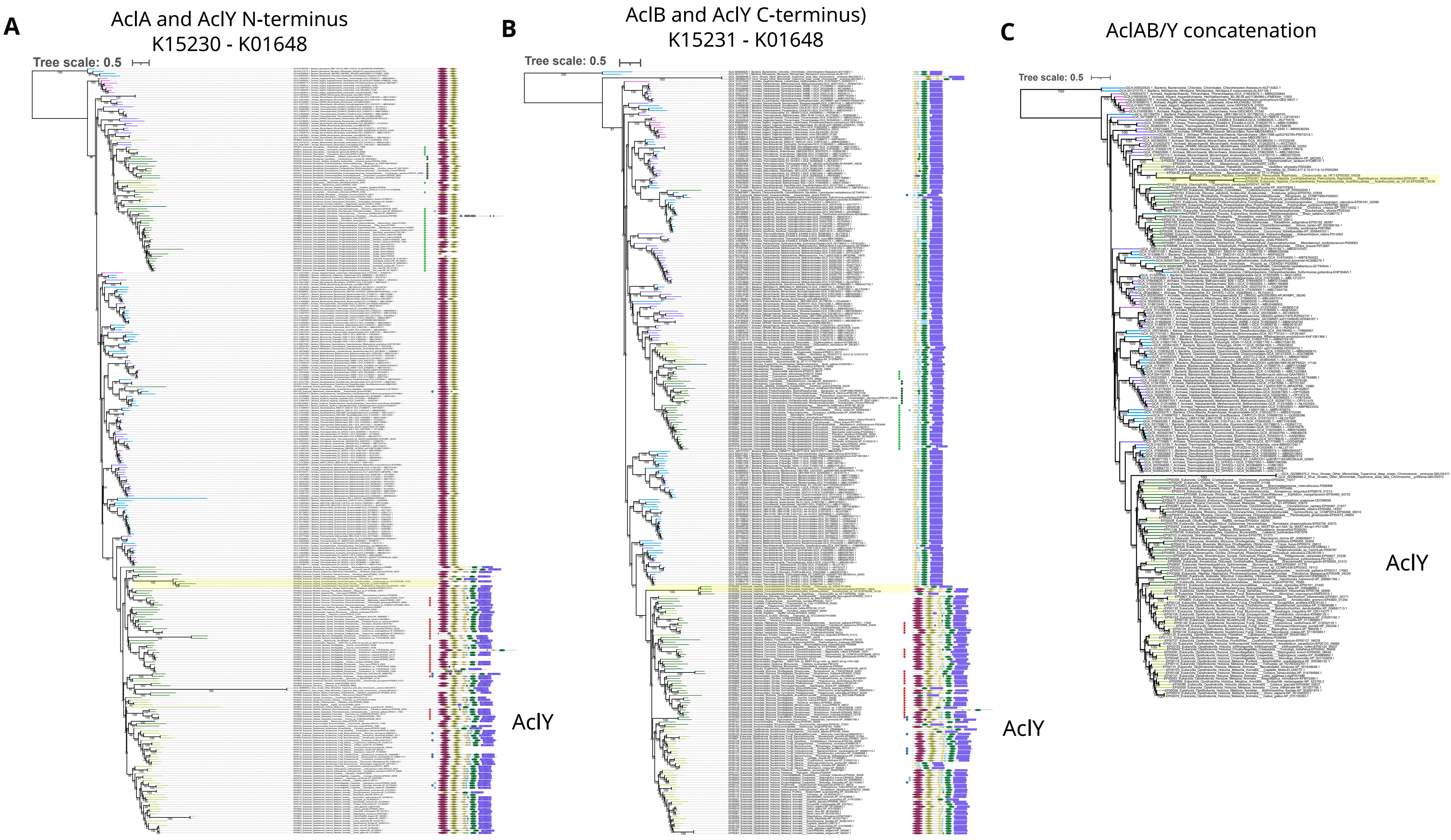

**Supplementary Figure 30.**

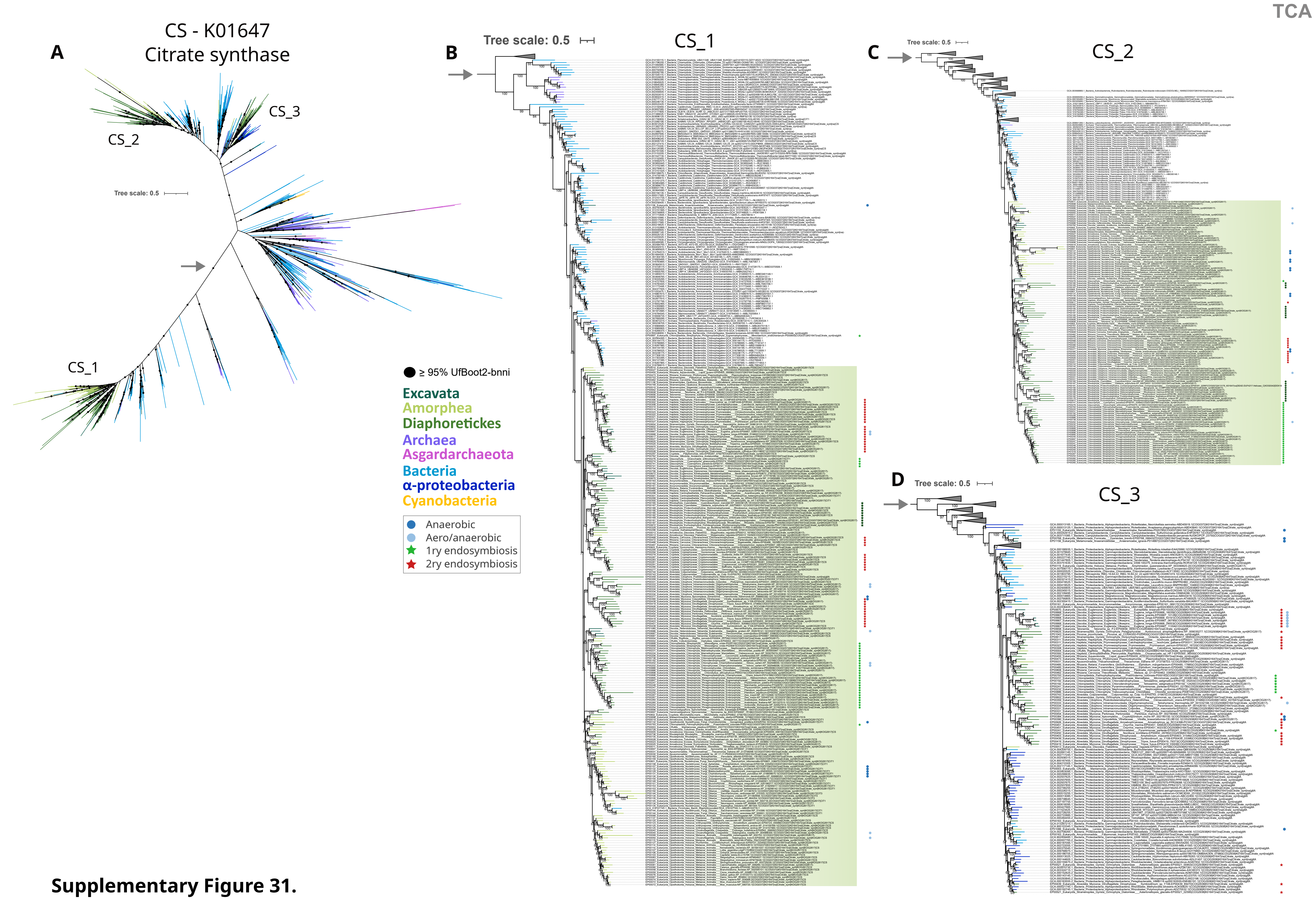

**Supplementary Figure 31.**

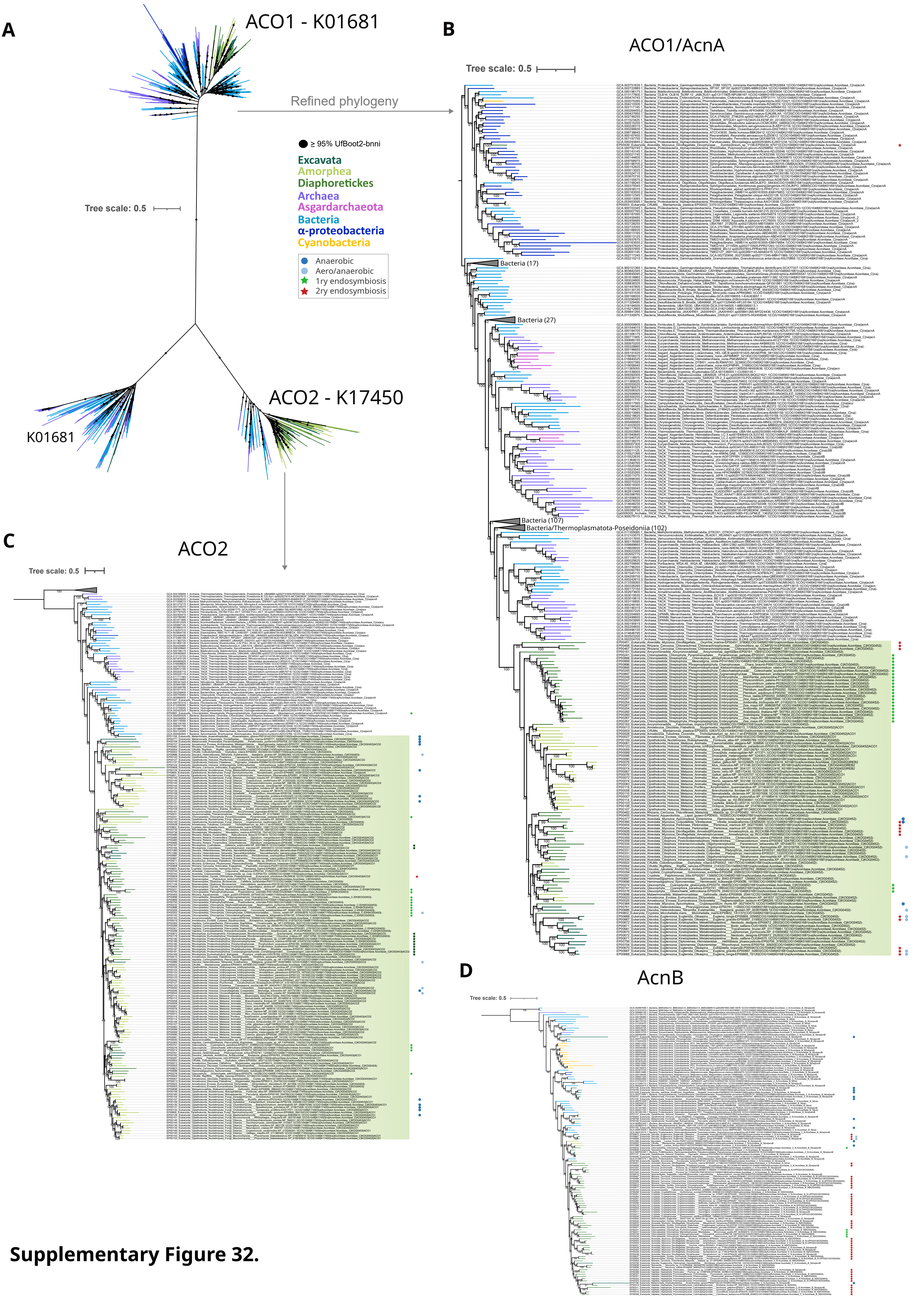

**Supplementary Figure 32.**

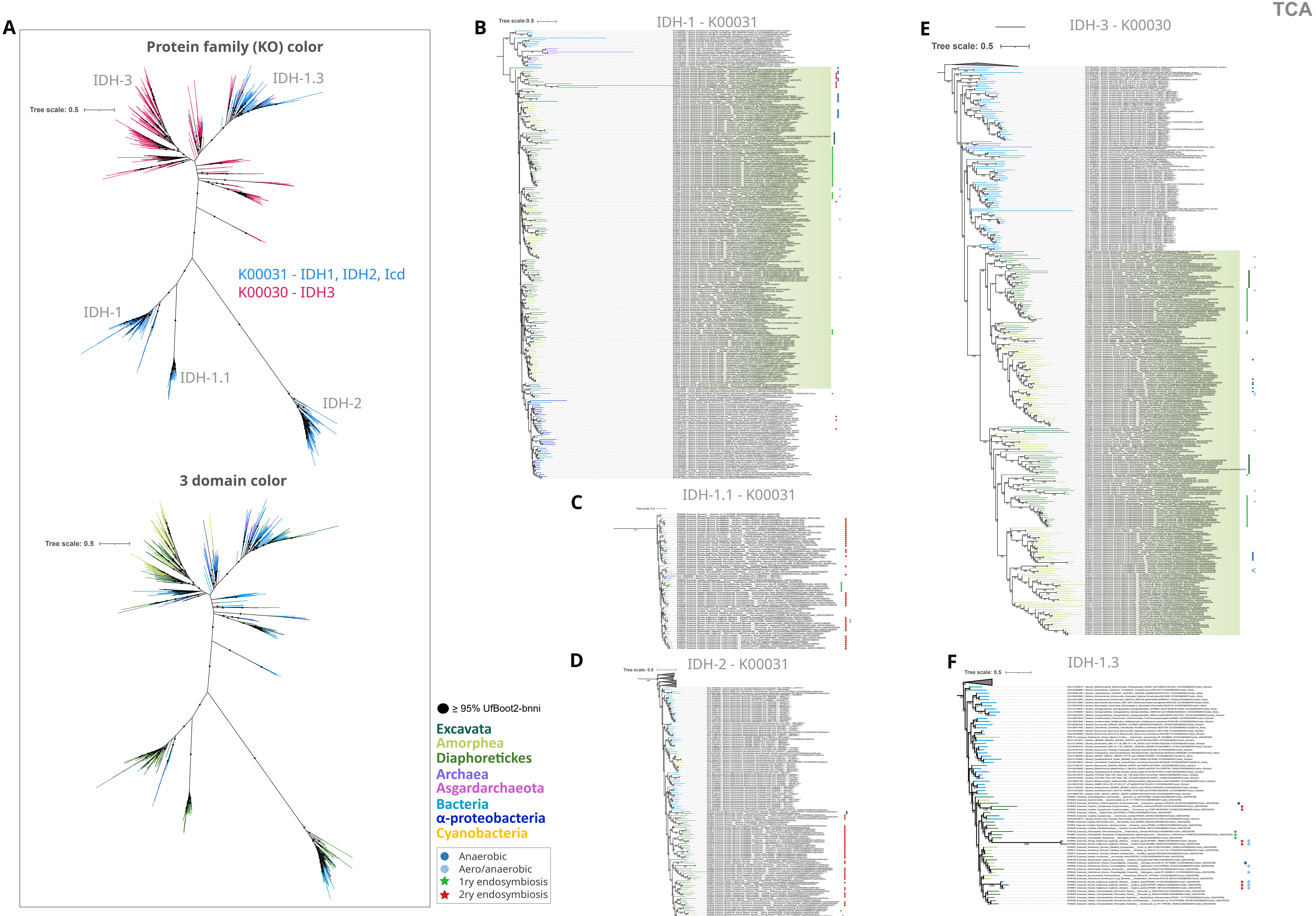

**Supplementary Figure 33.**

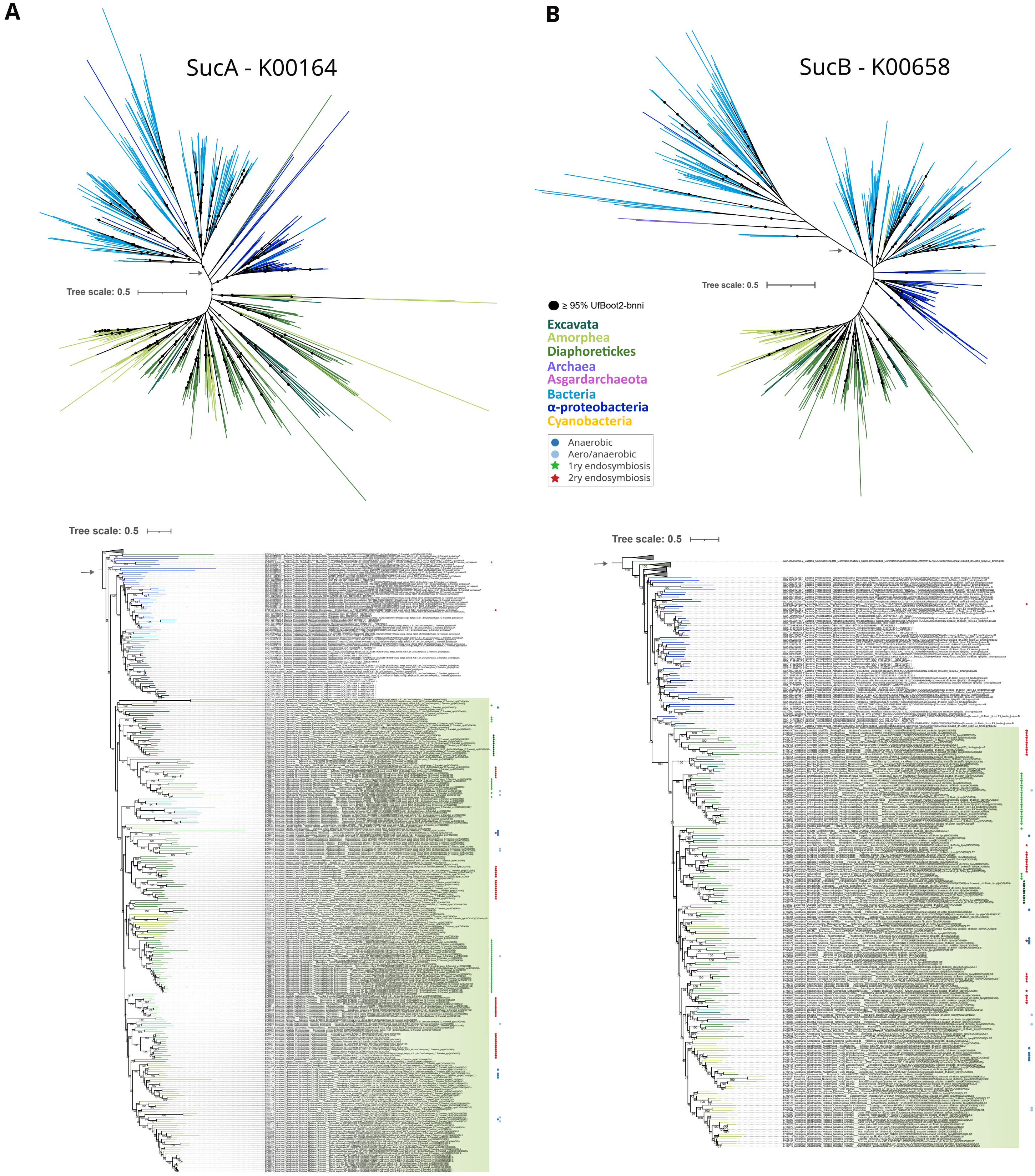

**Supplementary Figure 34.**

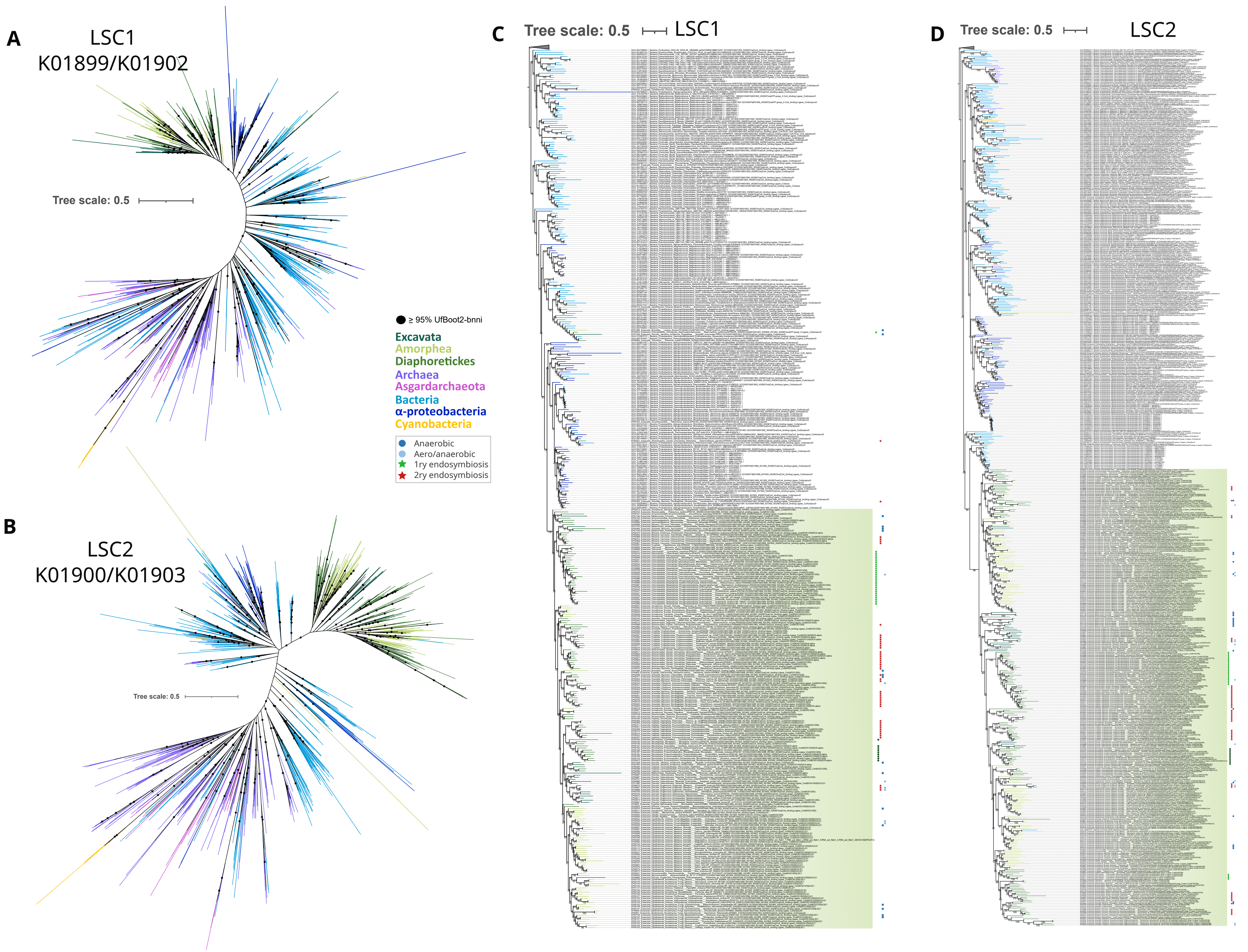

**Supplementary Figure 35.**

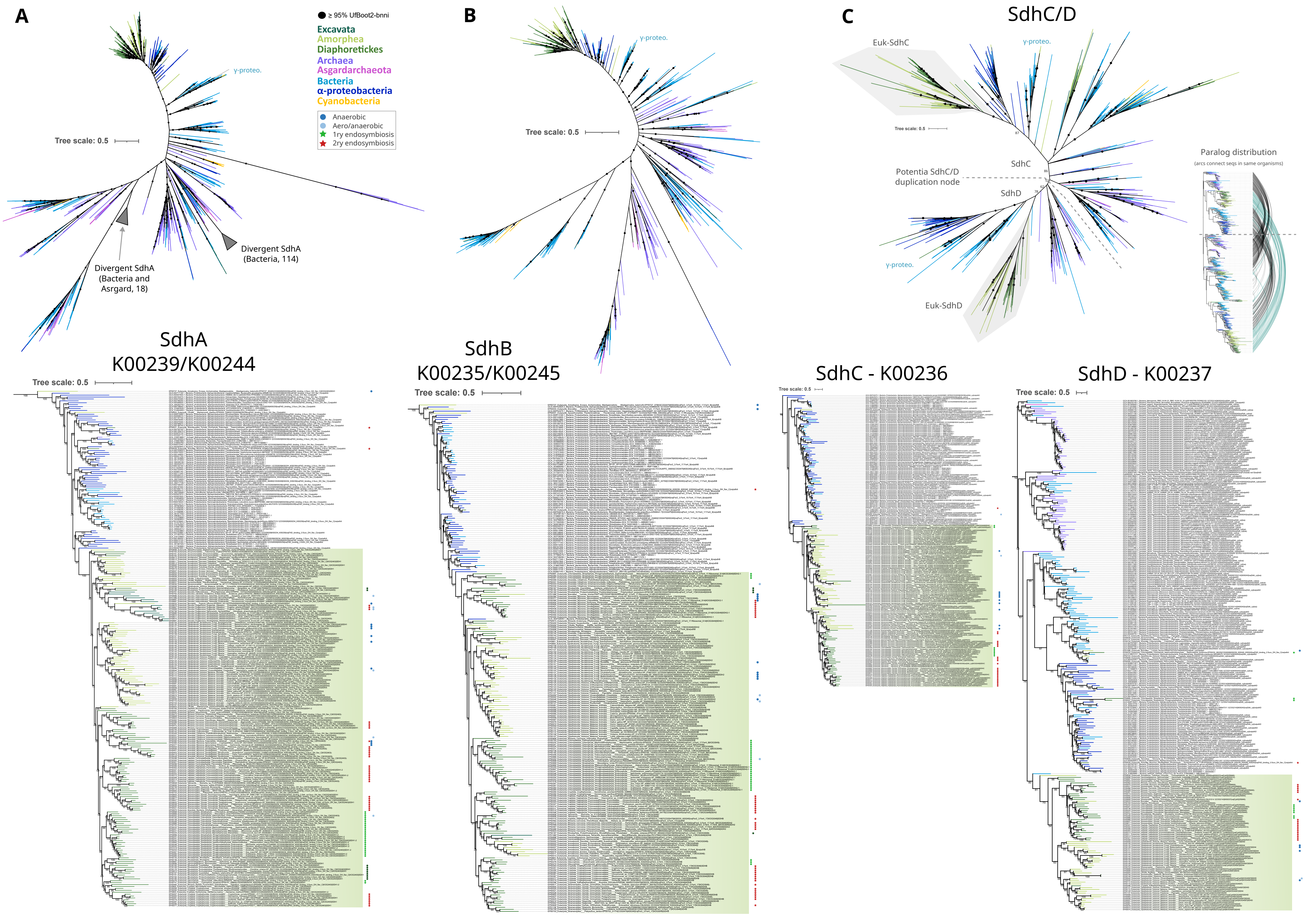

**Supplementary Figure 36.**

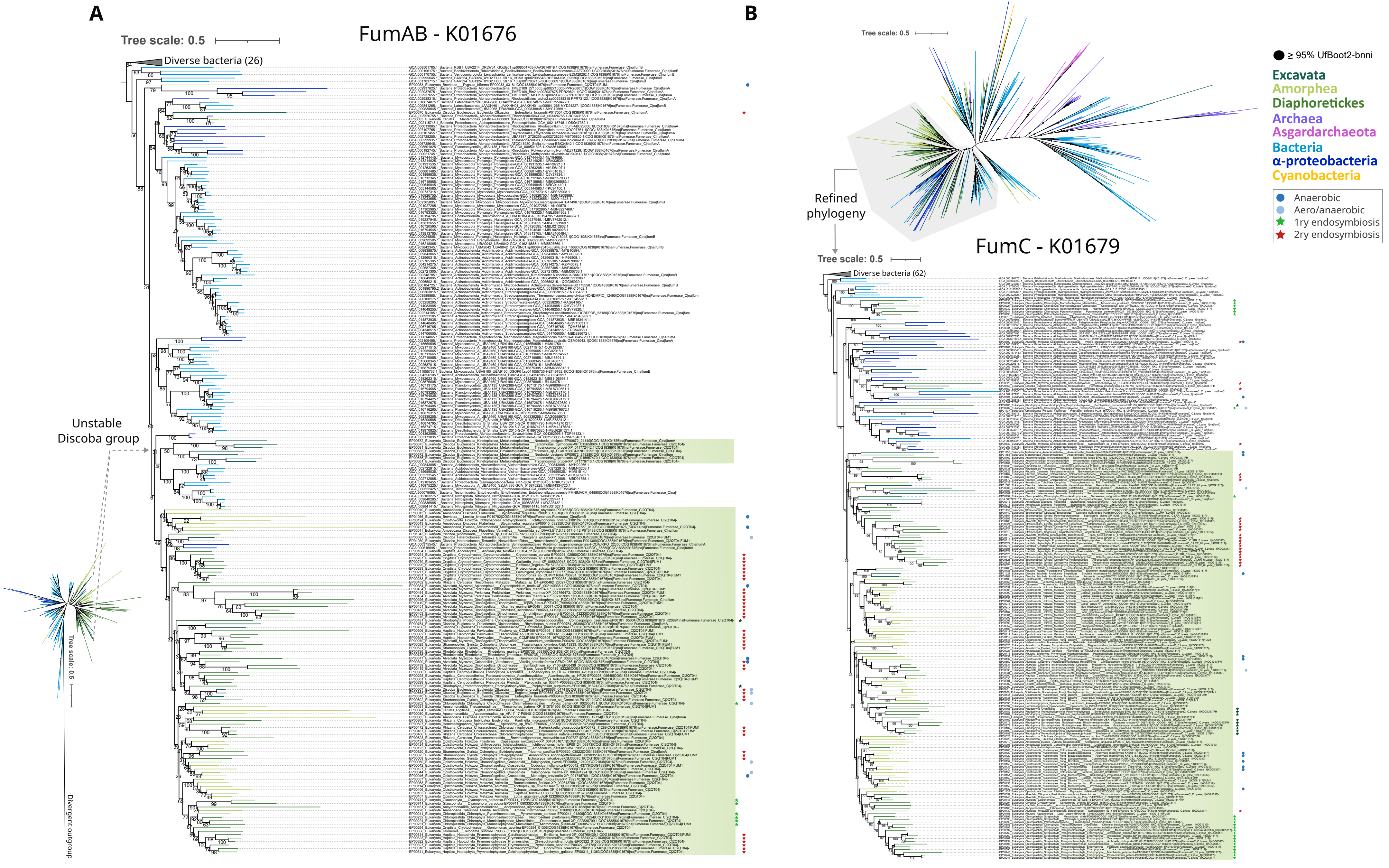

**Supplementary Fiugre S37.**

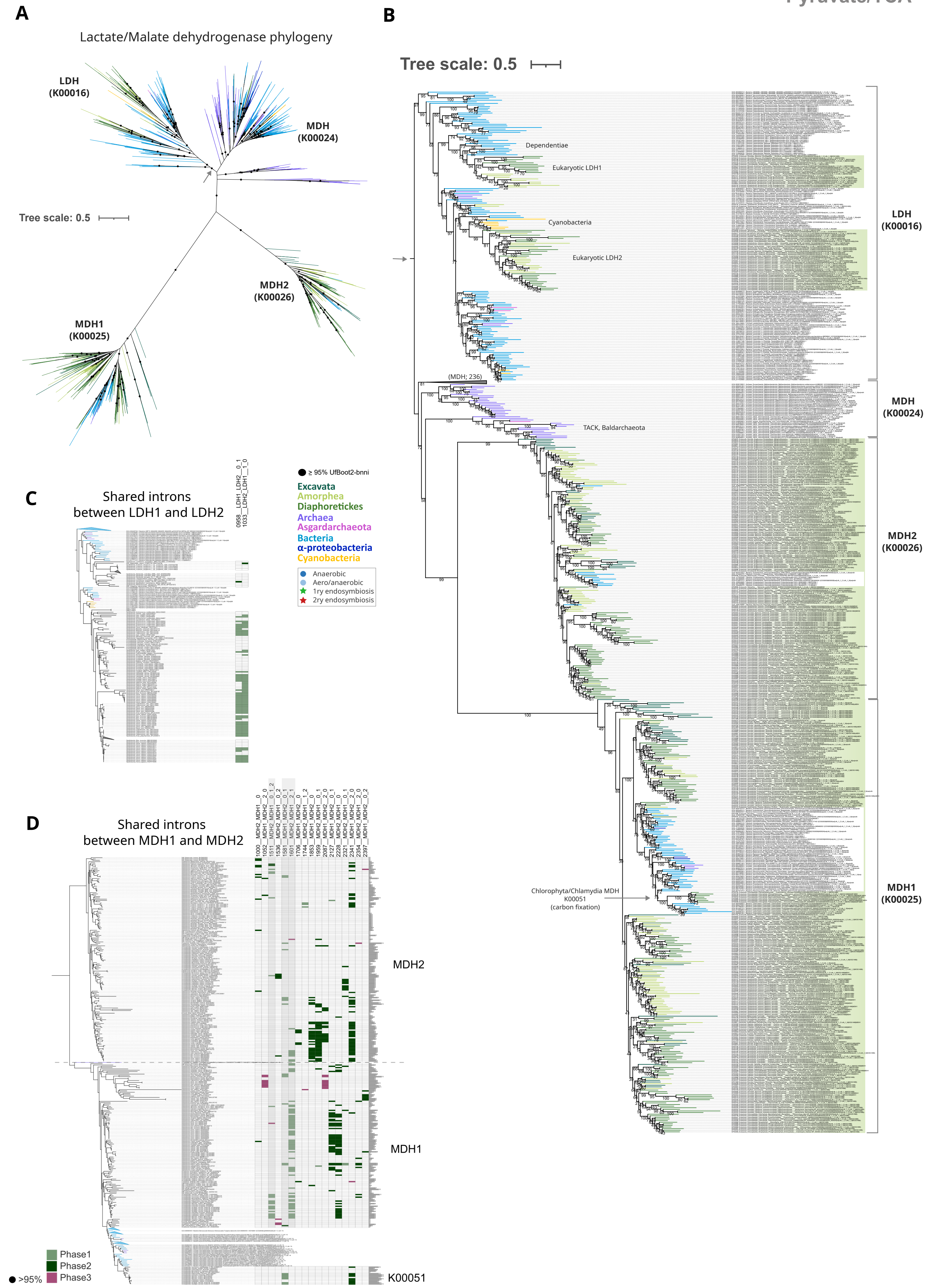

Supplementary Figure 38.

TCA

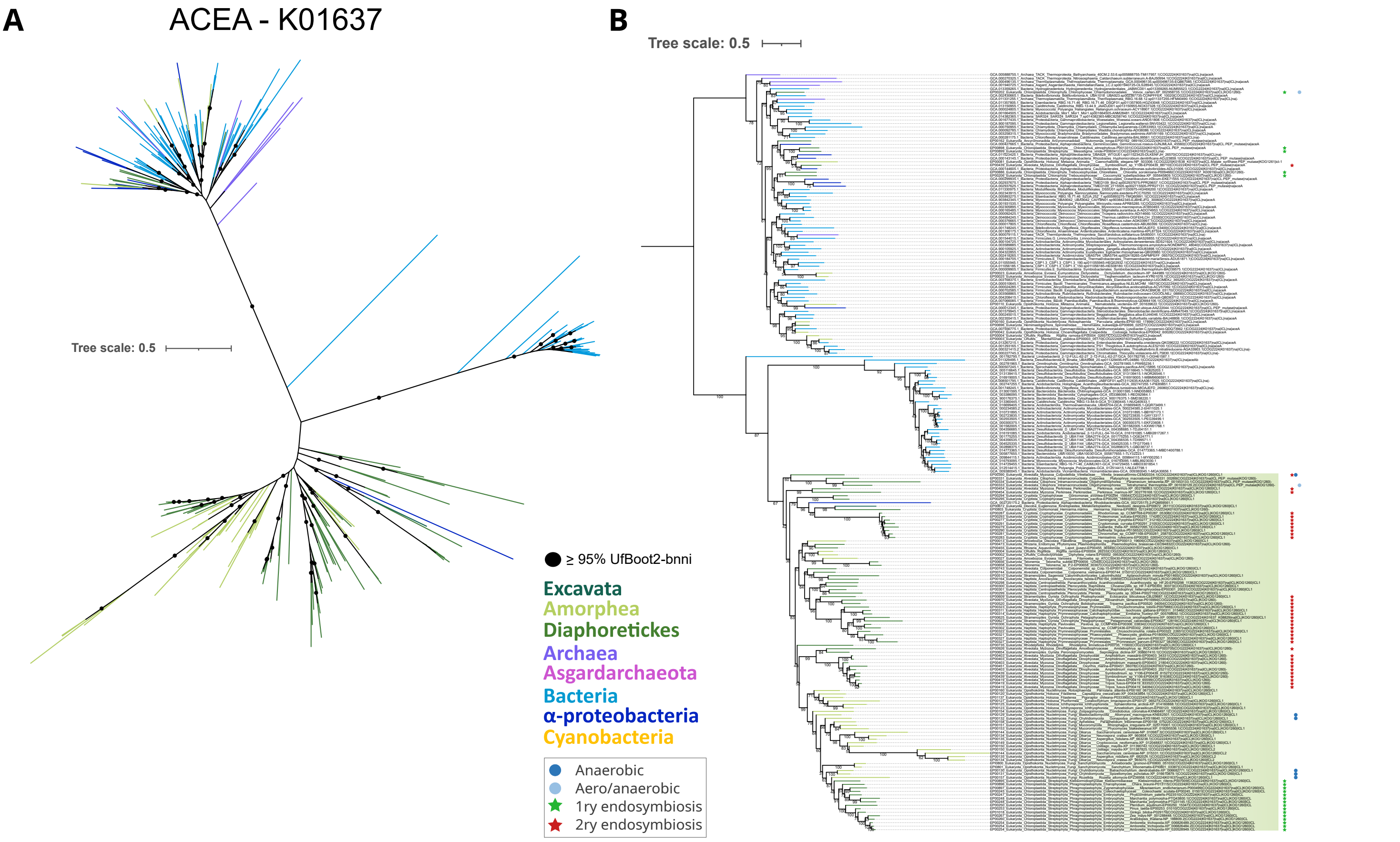

**Supplementary Figure 39.**

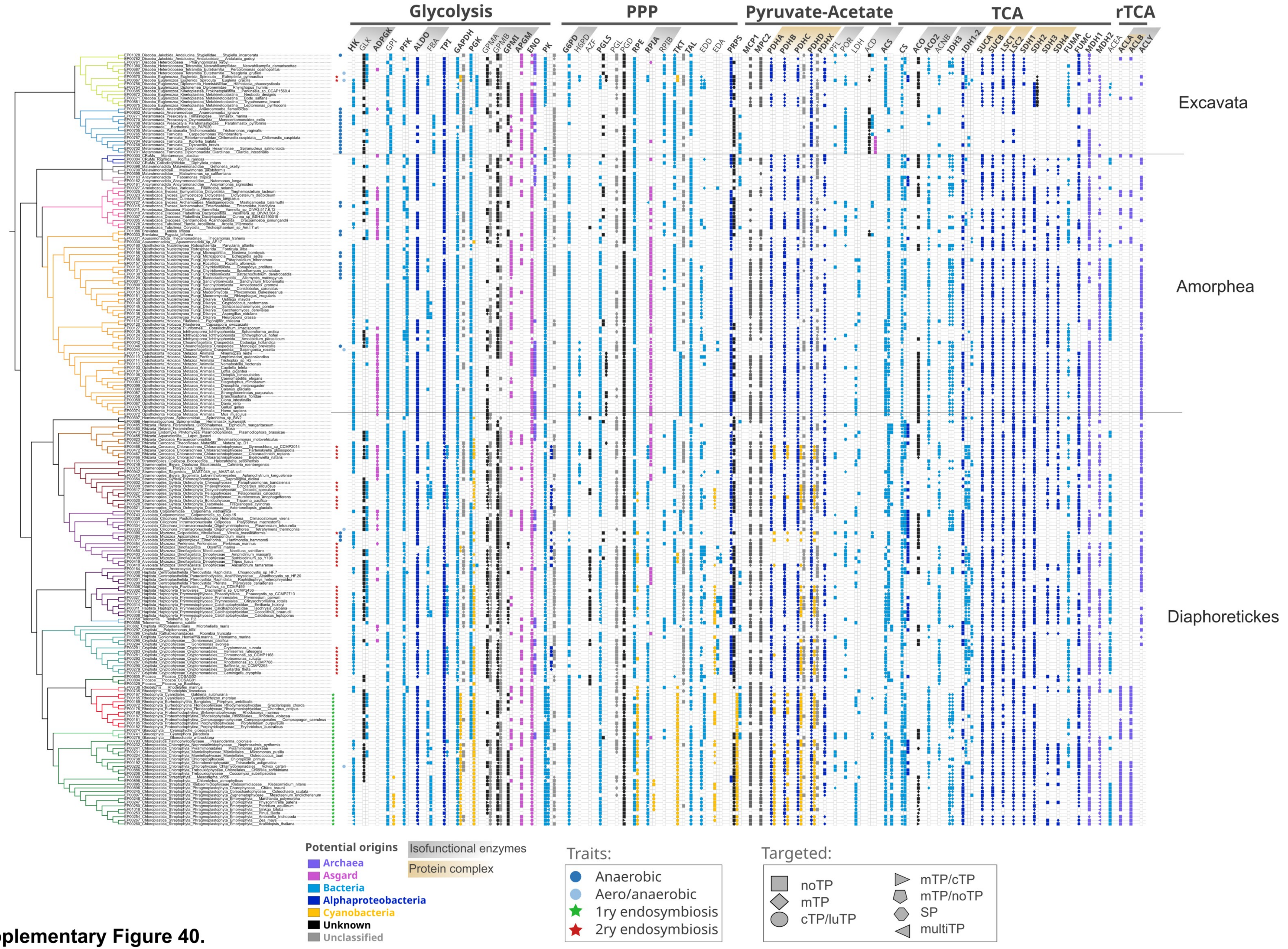

**Supplementary Figure 40.**

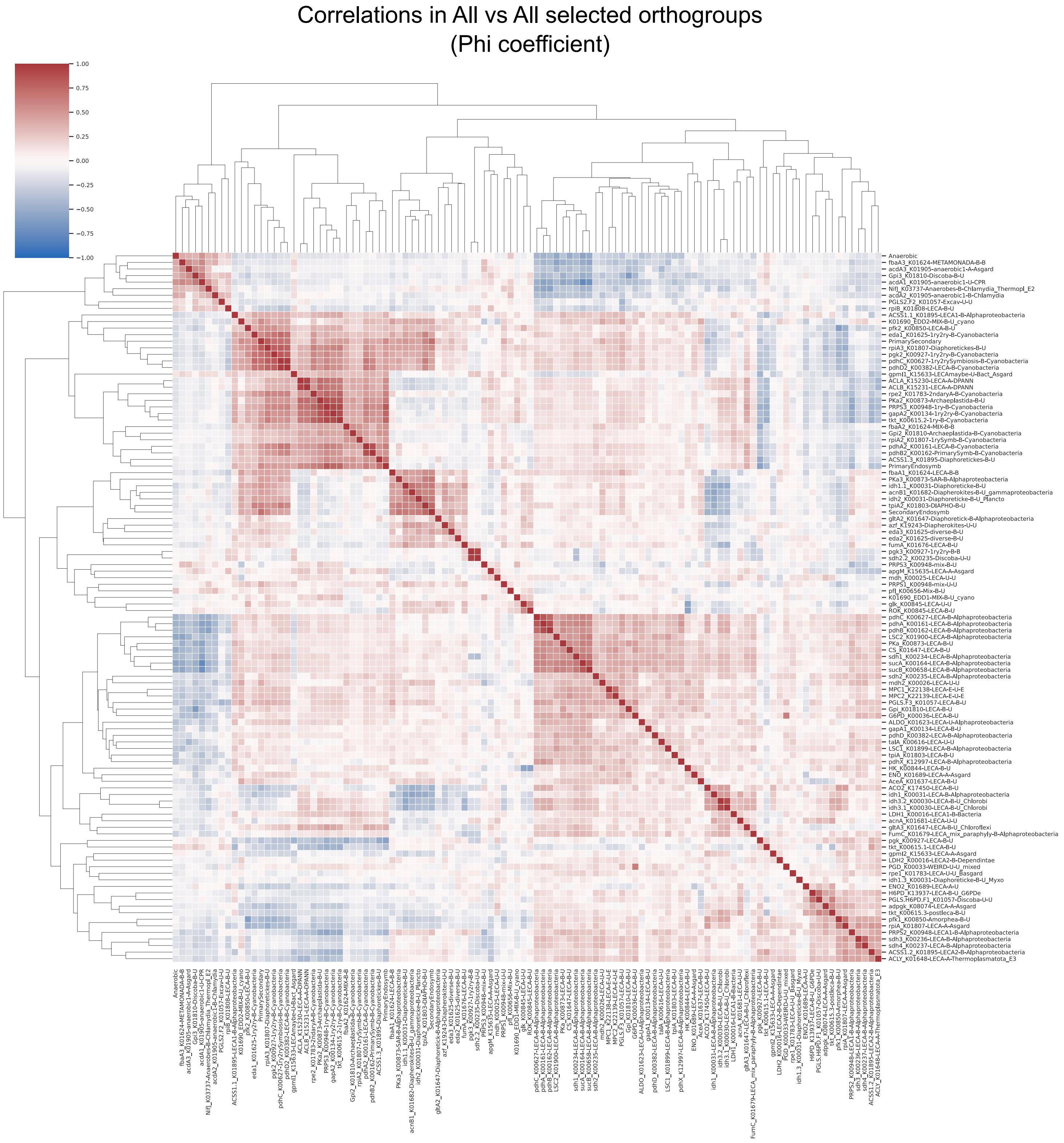

**Supplementary Figure 41.**

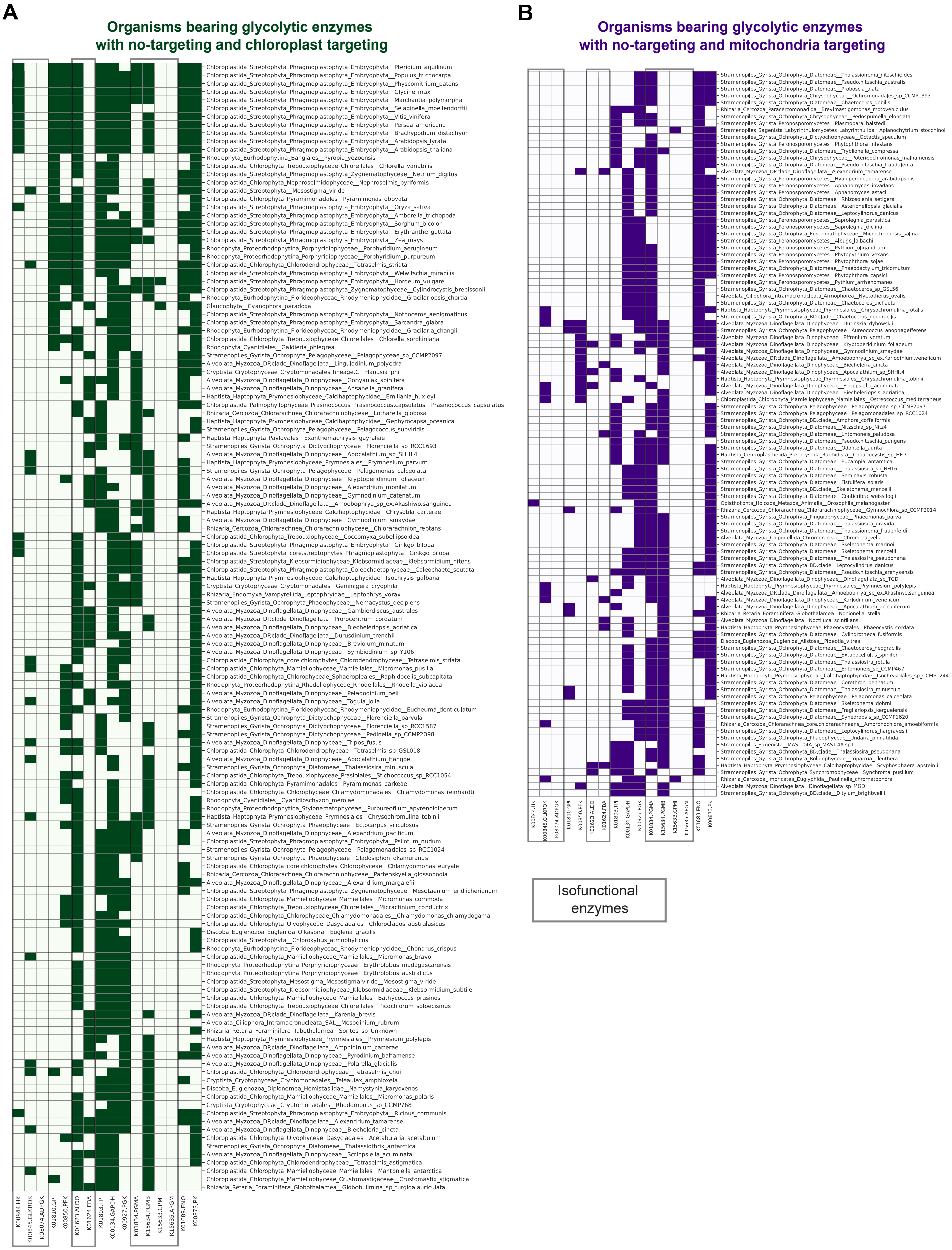

**Supplemnetary Figure 42.**

Evolutionary reconstruction EMP-Glycolysis and PPP in Archaeplastida

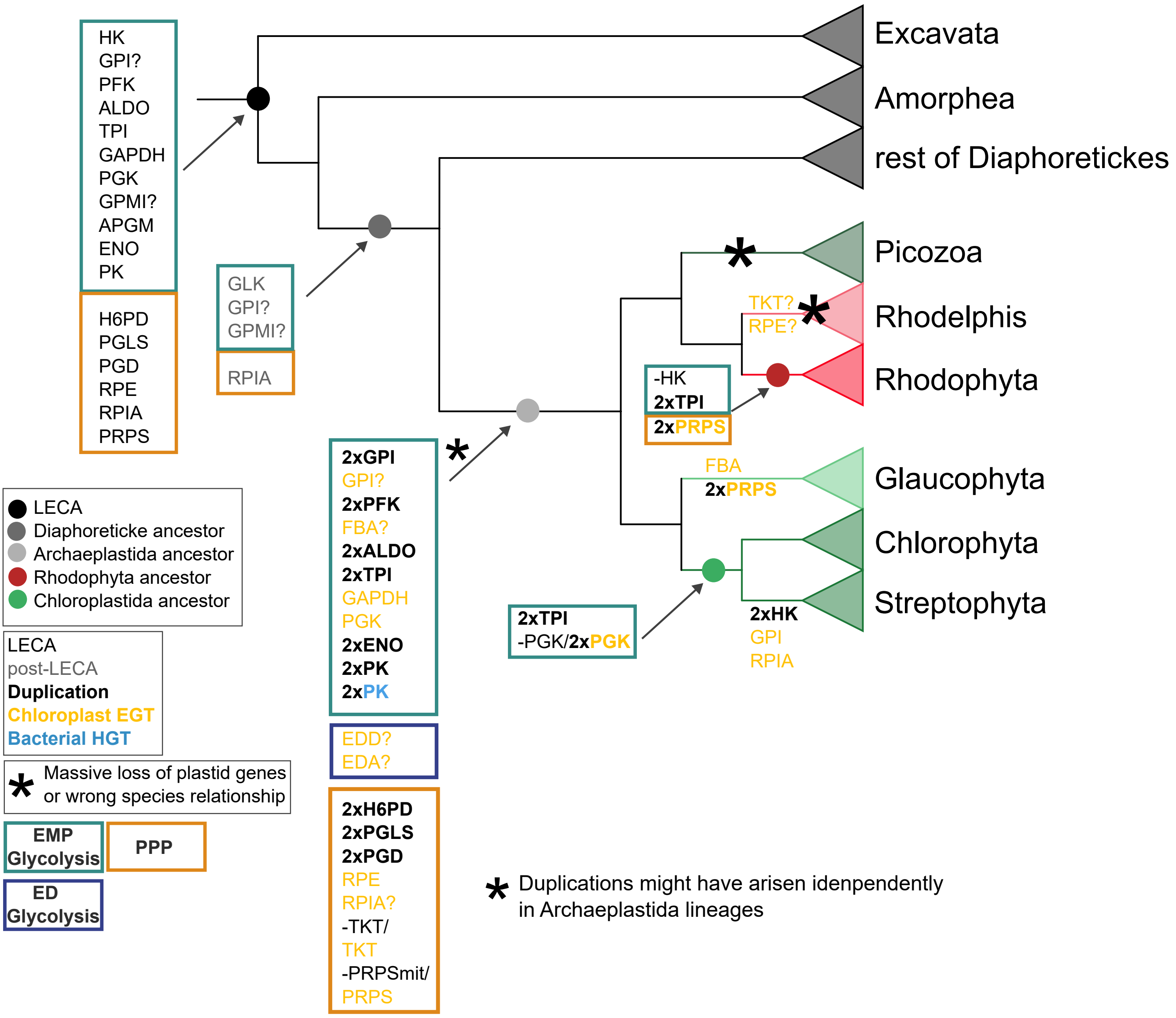

**Supplementary Figure 43.**

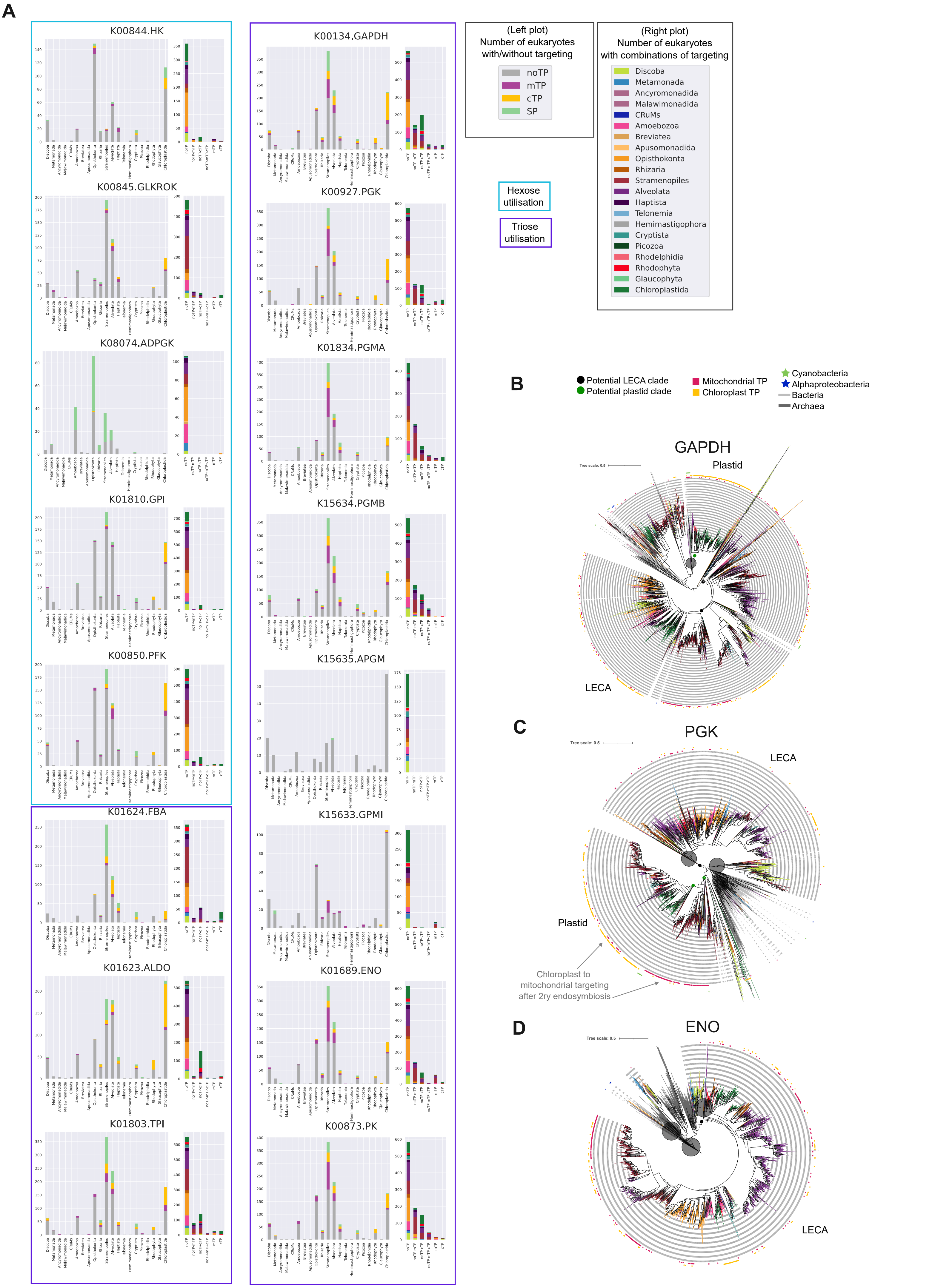

**Supplementary Figure 44.**
